# Type 1 interferon perturbates clonal competition by reshaping human blood development

**DOI:** 10.1101/2022.09.28.509751

**Authors:** Chhiring Lama, Danielle Isakov, Shira Rosenberg, Miguel Quijada-Álamo, Mirca S. Saurty-Seerunghen, Sara Moein, Tsega-Ab Abera, Olivia Sakaguchi, Mansi Totwani, Grace Freed, Chi-Lam Poon, Neelang Parghi, Andrea Kubas-Meyer, Amy X. Xie, Mohamed Omar, Daniel Choi, Franco Castillo-Tokumori, Ghaith Abu-Zeinah, Alicia Dillard, Nathaniel D. Omans, Neville Dusaj, Paulina Chamely, Eleni Mimitou, Peter Smibert, Heidi E. Kosiorek, Amylou C. Dueck, Rona Weinberg, Ronan Chaligne, Bridget Marcellino, Luigi Marchionni, Sanjay Patel, Paul Simonson, Dan A. Landau, Elvin Wagenblast, Ronald Hoffman, Anna S. Nam

## Abstract

Inflammation perturbs evolutionary dynamics of hematopoietic stem cell (HSC) clones in clonal hematopoiesis and myeloid neoplasms. We studied HSCs, progenitors and immune cells from patients with myeloproliferative neoplasm (MPN) at baseline and following interferon-⍺ (IFN⍺) treatment, the only MPN therapy to deplete clonal stem cells. We focused on essential thrombocythemia, an informative model of early-phase neoplastic hematopoiesis. We integrated somatic genotyping, transcriptomes, immunophenotyping, and chromatin accessibility across single cells. IFN⍺ simultaneously activated HSCs into two polarized states, a lymphoid progenitor expansion associated with an anti-inflammatory state and an IFN⍺-specific inflammatory granulocytic progenitor (IGP) state derived directly from HSCs. The augmented lymphoid differentiation balanced the typical MPN-induced myeloid bias, associated with normalized blood counts. Clonal fitness upon IFN⍺ exposure was due to resistance of clonal stem cells to differentiate into IGPs. These results support a paradigm wherein inflammation perturbs clonal dynamics by HSC induction into the precipitous IGP differentiation program.

**One-Sentence Summary:** Inflammation accelerates clonal evolution by driving stem cell differentiation into an alternate interferon-⍺-induced progenitor state.

## Introduction

Systemic inflammation increases with aging and is implicated in accelerating the development of clonal hematopoiesis (CH) and myeloid neoplasms (*1–8*). Clonal stem cells may be resistant to inflammatory signaling that leads to functional defects in hematopoietic stem cells (HSC) (*4, 6–8*). On the other hand, inflammatory cytokines, such as interferon-⍺ (IFN⍺), directly activate HSCs into cell cycle entry in mice (*9–11*), an observation that has been proposed to undergird both enhanced clonal expansion upon inflammation (*5*) and clonal depletion in the setting of IFN⍺ therapy for myeloproliferative neoplasms (MPN) (*9, 12, 13*). Indeed, the ability of IFN⍺ to modulate clonal dynamics in patients with MPN presents a unique opportunity to assess the effects of chronic IFN⍺ signaling on clonal HSC fitness in human.

MPNs are driven by somatic mutations in *CALR*, *JAK2* or *MPL* that override the highly regulated process of hematopoiesis resulting in an overproduction of one or more myeloid lineages, such as increased platelet production in essential thrombocythemia (ET) (*14*). IFN⍺ is the only clonally selective MPN treatment, often effecting molecular response (*12, 13, 15*). Even in the absence of molecular response, IFN⍺ treatment frequently induces normalization of the patients’ blood counts (*12, 13, 15*). To define the downstream effects of IFN⍺ therapy that undergird the phenotypic response and clonal dynamics in human, we require methods that can isolate the differential IFN⍺ effects on mutated stem cells from the wildtype. However, as clonal cells cannot be distinguished from the admixed wildtype cells via cell surface markers, we leveraged single-cell multi-omics platforms that detect the mutational status and whole transcriptomes (*16*) with immunophenotyping or chromatin accessibility data, within thousands of individual cells. These methods allowed us to overlay two hematopoietic differentiation landscapes—one mutated and the other wildtype—from the same individual, thus facilitating a direct comparison between mutated and wildtype cells both at baseline and following treatment. We focused on *CALR*-mutated ET due to the heterogeneous molecular response despite clinical response in most patients (*13, 17*) and applied these multi-modality single-cell methods to CD34^+^ hematopoietic stem and progenitor cells (HSPC) and immune cells from serial bone marrow sampling from patients treated with IFN⍺ for at least one year. This approach enabled us to assess the phenotypic and epigenetic alterations, jointly together with clonal dynamics, induced by IFN⍺ in human neoplasm.

## Results

### IFN⍺ paves an alternate route of granulocytic differentiation

To define the effects of IFN⍺ on wildtype and neoplastic hematopoiesis in human, we leveraged the Genotyping of Transcriptomes (GoT) technology that simultaneously captures the mutation status and whole transcriptomes in thousands of single cells (*16*). To overcome inter-patient variability in baseline hematopoiesis, we applied GoT to FACS-isolated CD34^+^ cells from serial (i.e., baseline and treated) bone marrow from individuals who were diagnosed with *CALR*-mutated ET (*18*) (**Fig. 1A**). As serial bone marrow biopsies are not typically performed in the absence of suspected disease progression, we utilized cryopreserved specimens from the MPN-RC-111 and -112 clinical trials wherein patients were treated weekly with a pegylated form of IFN⍺ (*19, 20*) (**Fig. 1A**, n = 10 individuals, 8 baseline samples, 13 treated samples; additional 3 baseline samples included from our previous work (*16*); see **table S1** for patient and sample information). Patients with samples available for this study exhibited partial or complete clinical response (i.e., improvement in platelet counts) and were representative of the other patients with *CALR* mutations in these clinical trials (**fig. S1A**, **table S1**). We incorporated time-point specifying barcoded antibodies (*21*) that enabled multiplexing baseline and IFN⍺-treated samples into the same GoT experiments, in order to obviate technical batch effects (e.g., sequencing depth) between serial samples (**Fig. 1B**). We also advanced the GoT method by incorporating immunophenotyping (*22*) (GoT-IM) to link transcriptional and immunophenotypic cell identities (**Fig. 1B**). GoT-IM provided genotyping data for the canonical *CALR* frameshift mutations for 72% of CD34^+^ HSPCs (n = 46,883 cells of total 65,452 cells), consistent with our previously reported genotyping rates (*16*). In this way, we obtained somatic genotyping, whole transcriptomes, immunophenotyping and treatment status for thousands of cells from the same GoT-IM experiment.

**Figure 1.**
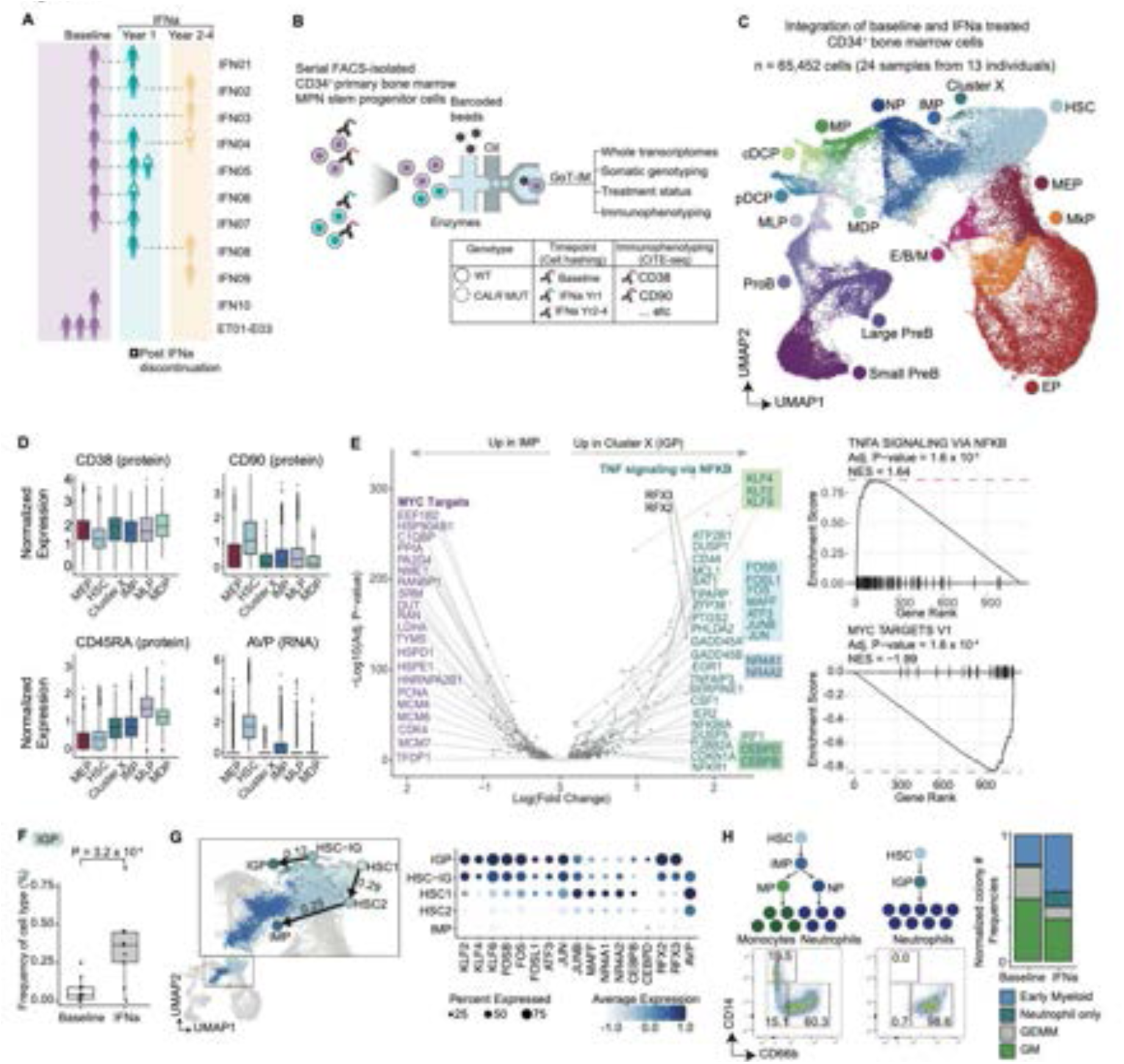
Integration of Genotyping of Transcriptomes with immunophenotyping and IFN⍺-treatment status identifies an alternate IFN⍺-specific cell state in MPN stem and progenitor cells. **A.** Primary bone marrow samples at baseline and after IFN⍺ treatment included for Genotyping of Transcriptomes with Immunophenotyping (GoT-IM) on CD34^+^ cells. **B.** Schematic of GoT-IM via CITE-seq and Cell Hashing. MPN, myeloproliferative neoplasms; WT, wildtype; MUT, mutated. **C.** Uniform manifold approximation and projection (UMAP) of CD34^+^ cells (n = 65,452 cells) from MPN samples (n = 24 samples from 13 individuals), overlaid with cell type assignment. HSC, hematopoietic stem cells; IMP, immature myeloid progenitors; NP, neutrophilic progenitors; MP, monocytic progenitors; cDCP, classic dendritic progenitors; pDCP, plasmacytoid dendritic progenitors; MDP, monocytic dendritic progenitors; MLP, multipotent lymphoid progenitors; E/B/M, eosinophil/basophil/mast cell progenitors; MkP, megakaryocytic progenitors; EP, erythroid progenitors; MEP, megakaryocytic-erythroid progenitors. **D.** Box plots showing normalized expression of HSC-defining protein and RNA markers. **E.** Volcano plot showing genes differentially expressed (DE) between Cluster X (i.e., IGPs) and IMPs identified via linear mixed modeling (LMM) with/without cluster identity (left, **methods**). Highlighted are genes enriched in the MYC pathway (purple) and TNF⍺ signaling via NF-κB (blue); Boxes represent transcription factors (TF) of the AP-1 (blue), KLF (light green), NR4A (purple), CEBP (dark green) families. Pre-ranked gene set enrichment analysis using the MSigDB Hallmark collection (right). **F.** Box plot showing normalized IGP frequency at baseline and IFN⍺ treatment (n = 11 baseline samples, n = 9 treated samples). P-values from likelihood ratio test of LMM with/without IFN⍺ treatment (**methods**). **G.** Integrated UMAP highlighting IGP, IMP and the HSC subclusters with RNA velocity-based cell state trajectory for IFN⍺-treated cells (left, trajectory presented corresponds only to IMP, IGP and the HSC subclusters, see **table S5, methods**). Dot plot showing gene expression levels of upregulated TFs in IGPs (right). **H.** Representative flow cytometric analysis of colonies from the single-cell differentiation of individually sorted CD34^+^, CD90^+^ HSCs (left). Normalized colony frequency from HSCs sorted from bone marrow samples from patients at baseline and following IFN⍺ (right, n = 2 baseline; n = 3 IFN⍺-treated). The number of colonies were down-sampled to the same minimum count for each replicate for equal representation. GEMM, granulocyte, erythrocyte, monocyte and megakaryocyte; GM, granulocyte-monocyte colonies .

We hypothesized that IFN⍺ may alter cell states but not induce novel cellular identities. We integrated across the individual 24 samples to define the cell identities of the CD34^+^ HSPCs consistently across individual sampling (after analytically segregating the cells by time-point from the same experiments, **Fig. 1C**, **fig. S1B-F, methods**) (*23*). As single-cell gene expression provides high-resolution mapping of the HSPC identities, we clustered the cells based on gene expression data and annotated the clusters based on canonical cell markers (**fig. S2A-C**, see **table S2** for cell numbers)(*16, 24–26*). To identify HSCs, we leveraged the jointly captured immunophenotyping to identify the CD38^-^, CD45RA^-^, CD90^+^ HSCs (with high RNA expression of the HSC marker *AVP* (*27*), **Fig. 1D**, **fig. S2D**). We observed the expected cell types, such as megakaryocytic progenitors (MkP) and immature myeloid progenitors (IMP, consisting predominantly of phenotypically defined common myeloid progenitors (CMP) and granulo-monocytic progenitors (GMP) (*24*), **Fig. 1C**, **fig. S2A-C**).

Contrary to our hypothesis that IFN⍺ may not induce novel cellular identities, we identified an unknown cluster (Cluster X), previously not described in studies of normal or MPN bone marrow CD34^+^ cells (*16, 24–26, 28–30*) (**Fig. 1C**). Cluster X was immunophenotypically similar to the IMPs based on CD38^mid^, CD45RA^mid^, and CD90^-^ expression (**Fig. 1D**). To elucidate the identity of Cluster X, we performed differential expression analysis between Cluster X and IMPs (**Fig. 1E**, **left, table S3,** linear mixed model that explicitly models the effects of patient batch and treatment status, see **methods**). We observed a striking upregulation of the immediate early response transcription factors (TF) of the AP-1 (*JUN, FOS, JUNB, FOSB, ATF3, FOSL1, MAFF*), KLF (*KLF2, KLF4, KLF6*), and NR4A (*NR4A1, NR4A2*) families (**Fig. 1E**, **left, table S3**). Other TFs included interferon regulatory factor 1 (*IRF1*), indicating an inflammatory response. In addition, we observed a robust upregulation of *RFX2* and *RFX3* TFs. While the *RFX2/3* factors are not well characterized in HSPCs, RFX2 activity was identified as one of the key pro-survival transcription factors activated in neutrophils during an inflammatory challenge in mice, particularly in the transition from bone marrow to blood (*31*). Upregulation of *CEBPB* and *CEBPD*, implicated in emergency granulopoiesis (*32*) and granulopoiesis under cellular stress (*33*), respectively, further suggested a neutrophilic differentiation trajectory. Gene set enrichment analysis identified the upregulation of TNF⍺ signaling via NF-κB pathway (Adj. P-val = 1.6 x 10^-4^), and downregulation of the MYC targets (Adj. P-val = 1.6 x 10^-4^, **Fig. 1E**, **right, table S4**). Incorporation of other HSPC immunophenotypic markers revealed that these cells were also positive for CD44, CD117, dim CD66b, and negative for HLA-DR, similar to IMPs and neutrophil progenitors (**fig. S2B)**. Based on the transcription factors, immunophenotypes and pathways activated in Cluster X, we termed these cells inflammatory granulocytic progenitors (IGP). Consistent with the identification of the IGPs in this cohort of patients, the IGPs derived predominately from the IFN⍺-treated samples with elevated frequencies in the IFN⍺-treated versus baseline CD34^+^ cells (**Fig. 1F**, **fig. S2E**). As the frequencies of IGPs were low, we confirmed that the elevated IGP frequencies in the IFN⍺ treatment samples were not simply due to a greater number of total CD34^+^ cells captured in the treated samples (**fig. S2E-F**). Separately integrating the samples based on treatment status also confirmed the specificity of the IGPs to IFN⍺-treated bone marrow (**fig. S2G**). The presence of a distinct IGP cluster in dimensional reduction of cells from individual experiments without any batch correction reassured that IGPs were not a technical artifact of integration (**fig. S2H**). Altogether, these data revealed that IFN⍺ induces an alternate IGP state.

The differentially upregulated TFs in the IGPs were also highly enriched in a subset of quiescent HSCs with elevated *AVP* (*27*) and CD90 expression, we labeled HSC-IG (**Fig. 1G**, **fig. S2D, S3A**). The transcriptional similarities of the IGPs and HSC-IG (as revealed by their proximity on the UMAP space) suggested that the IGPs may derive from HSC-IG. RNA velocity measurements (*34, 35*) combined with partition-based graph abstraction (*36*) predicted cell state transitions from HSC-IG to IGPs (**Fig. 1G**, **left, table S5**). To define the transcriptional state transitions from HSC-IG to IGPs, we compared IGPs to HSC-IG and identified a reinforcement of the RFX3, AP-1, CEBPB/D, and KLF family TF expressions and downregulation of NR4A2 (**Fig. 1G**, **right, fig. S3B, table S3;** only IFN⍺-treated cells included in the differential expression analysis). As NR4A1/2 have been reported to maintain HSC quiescence (*37, 38*), their downregulation in IGPs relative to HSC-IG was consistent with the upregulation of differentiation and cell cycle-related genes in the IGPs (**Fig. 1G**, **right, fig. S3B-D, table S4**). Upregulation of *MPO* (encoding myeloperoxidase in primary granules) and *CSF3R* (encoding the G-CSF receptor), and downregulation of the MHC class II genes (*CD74, HLA-DPA1, HLA-DRB1, HLA-DPB1*) further provided evidence for its differentiation into the neutrophil lineage (**fig. S3B, table S4**).

To test the ability of HSCs to directly give rise to neutrophils, bypassing the conventional CMP and GMP oligo-potent progenitor states, we utilized a similar strategy by which the direct derivation of MkPs from HSCs (without traversing through the megakaryocytic-erythroid progenitor state) was demonstrated in human cells (*39*). We performed single cell colony forming unit assays by which we could track the differentiation of individual CD34^+^, CD90^high^ bone marrow HSCs from MPN patients at baseline and on IFN⍺ therapy (**Fig. 1H**, **fig. S3E, methods**). We identified that HSCs gave rise to mixed multilineage and monocyte-neutrophil colonies, consistent with a passage through the oligo-potent progenitor states, but HSCs from IFN⍺-treated patients also frequently gave rise to neutrophil-only colonies (CD66b^+^, CD16^+^, CD14^-^), supporting a direct passage to neutrophil development (**Fig. 1H**, **fig. S3E**). These data suggested that IFN⍺ induces an alternate and precipitous neutrophil developmental pathway that bypasses the typical granulo-monocytic bi-potent progenitor states.

### Inflammatory neutrophils are enriched in IFN⍺-treated bone marrow

To determine the identity of the immune cells that are downstream of the IGPs in an unbiased manner, we performed GoT-IM on the CD34^-^ compartments of the serial IFN⍺-treated samples (**Fig. 2A**, **fig. S4A-B**, see **table S6** for cell numbers, **methods**). Similar to the CD34^+^ cells, we clustered the cells based on gene expression and annotated the cell types based on canonical gene and protein markers (**Fig. 2B**, **fig. S4C**). We identified that the IGP gene signature was the highest in a distinct group of neutrophils (Neu1, **Fig. 2C**). Consistently, unsupervised co-embedding of the myeloid progenitors with the mature myeloid compartment revealed that the IGPs clustered with the Neu1 subset of ‘inflammatory’ neutrophils (**fig. S4D**). Comparison of the gene expression of the Neu1 subset to the other neutrophil group, Neu2 (at an equivalent stage of maturation based on the expression level of CD66b, CD11b, CD16, **Fig. 2B**) revealed that Neu1 expressed inflammatory cytokines such as *CXCL8* and *CXCL2* and the transcription factors observed in the IGPs including *RFX2/3*, *AP-1*, *KLF2/4/6*, and *CEBPB/D* (**Fig. 2D**, **left, table S7**). Gene set enrichment of the differentially expressed genes also highlighted TNF⍺ signaling via NF-κB and inflammatory pathways in the Neu1 subset (**Fig. 2D**, **right, table S7**), suggesting the propagation of the inflammatory state of the IGPs to the mature progeny. These inflammatory neutrophils derived predominantly from the IFN⍺-treated samples (**Fig. 2E**). Overall, these data demonstrated that IFN⍺ induced the development of inflammatory neutrophils.

**Figure 2.**
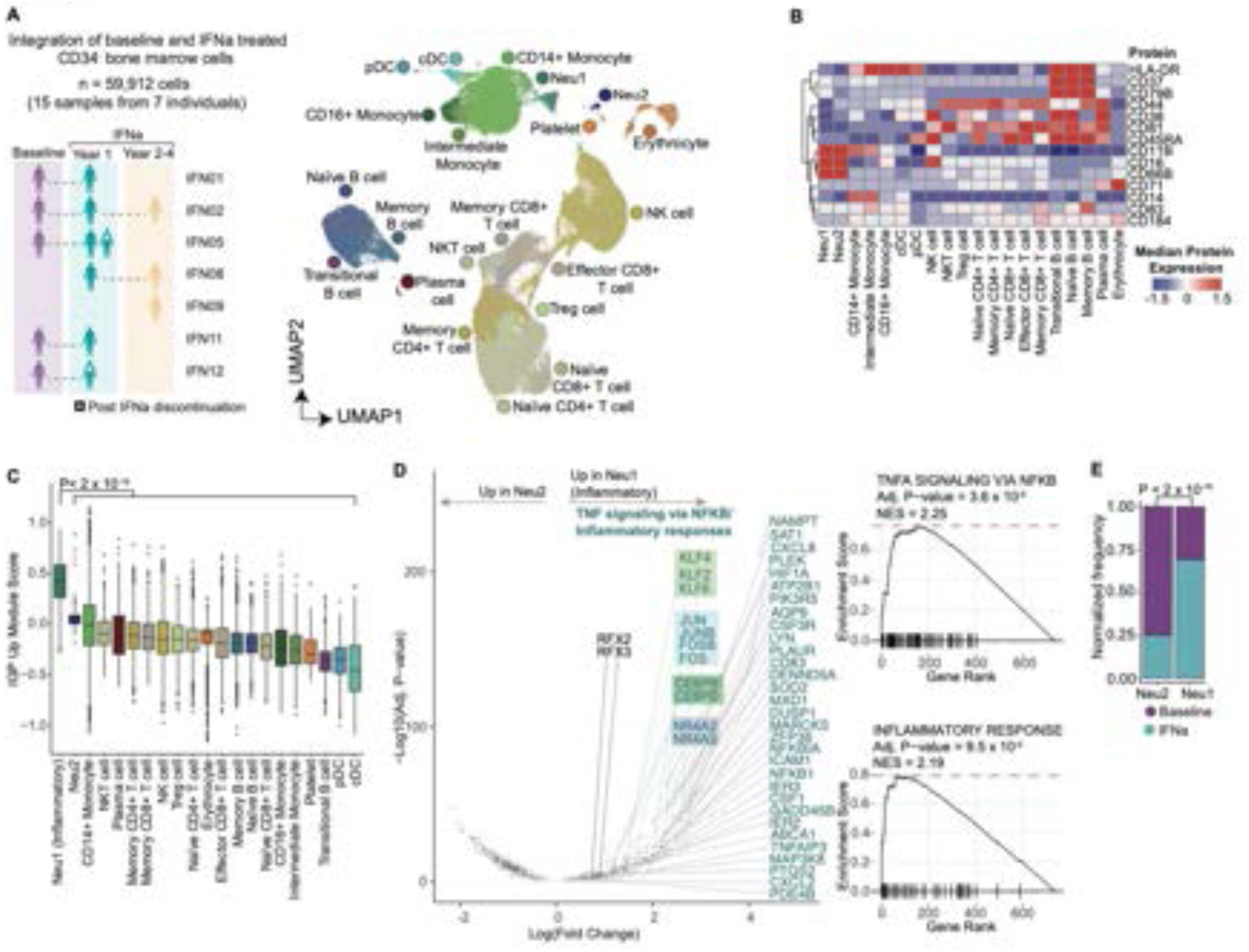
Inflammatory neutrophils are enriched in IFN⍺-treated bone marrow. **A.** Primary bone marrow samples at baseline and after IFN⍺ treatment included in GoT-IM on CD34^-^ cells (left). UMAP of CD34^-^ cells (n = 59,912 cells; 15 samples from 7 individuals), overlaid with cell type assignment (right). Neu1, Neutrophil subset 1; Neu2, Neutrophil subset 2; cDC, classic dendritic cells; pDC, plasmacytoid dendritic cells, Treg cell; Regulatory T cells, NKT cells; Natural Killer T cells. **B.** Heatmap showing median scaled expression of canonical immune cell protein markers from a representative patient IFN12. **C.** Box plots showing IGP-specific upregulated signature score (Fig. 1E**, table S3**) in CD34^-^ cell type clusters. P-value from likelihood ratio test of linear-mixed modeling (LMM) with/without cluster identity (**methods**). **D.** Volcano plot showing genes differentially expressed (DE) between Neu1 and Neu2 identified via LMM with/without cluster identity (left, **methods**). Highlighted are genes enriched in the TNF⍺ signaling via NF-κB (blue, box representation is same as Fig. 1E). Pre-ranked gene set enrichment analysis using the MSigDB Hallmark collection (right). **E.** Normalized frequency of baseline and IFN⍺-treated cells in Neu1 and Neu2 subsets. P-value from Fisher’s exact test, two-sided.

### IFN⍺ concurrently coordinates anti-and pro-inflammatory programs

To assess the global transcriptional impact of IFN⍺, we examined the transcriptional distance of HSCs between treatment time-points for individual patients. In an example case of patient IFN04 who showed partial clinical response (without evidence of disease progression, **table S1**), samples from three timepoints were available – (1) baseline, (2) at one year on active treatment, and (3) at four years but off therapy for 3 weeks at the time of collection. HSCs at year 1 displayed a strikingly distinct transcriptional profile compared to baseline cells, whereas cells that had been collected following discontinuation of therapy at year 4 were more similar to baseline HSCs (**Fig. 3A**), consistent with clearance of pegylated-IFN⍺ at ∼2-3 weeks (*40*). This contrasted with HSCs from samples with two timepoints under active IFN⍺ therapy (at years 1 and 2), which were similarly distinct from the baseline HSCs (**fig. S5A**). Projection of the progenitor identity assignments revealed that the HSPCs clustered based on cell identity as well as treatment status (**Fig. 3A**, **fig. S2H**). The magnitude of the transcriptional impact of IFN⍺ was in contrast to the subtler effects of somatic mutations, such as those in *CALR* (*16*)*, JAK2* (*29, 41*), and *DNMT3A* (*42*), resulting in co-mingling of mutated and wildtype cells, which could not be distinguished by scRNA-seq data alone, as revealed by methods that incorporate genotyping and scRNA-seq (*16, 29, 42, 43*). Thus, we first examined the impact of IFN⍺ on the overall hematopoiesis agnostic to genotype status.

**Figure 3.**
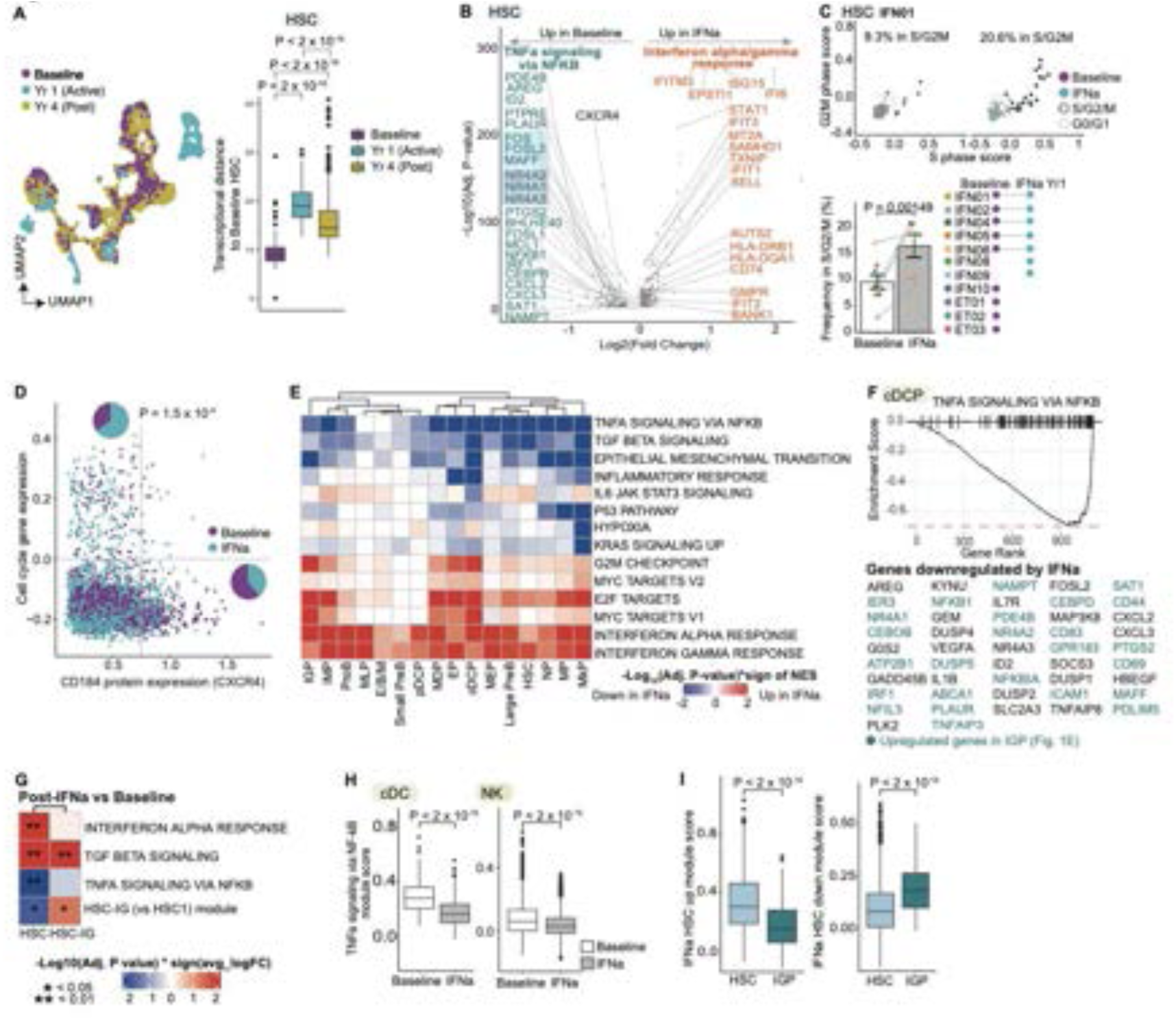
IFN⍺ concurrently regulates anti-and pro-inflammatory programs. **A.** UMAP showing a representative experiment that includes three time-points from patient IFN04 (n = 7,282 cells, left). Box plot showing transcriptional distance measurements between HSCs from each time-point and HSCs from baseline (right). Transcriptional distance corresponds to Euclidean distance of the first thirty principal components. P-values from Wilcoxon rank sum test, two-sided. **B.** Volcano plot showing genes differentially expressed (DE) between baseline and IFN⍺-treated HSCs via linear mixed modeling (LMM) with/without treatment status (**methods**). Genes highlighted in blue are those in the TNF⍺ signaling via NF-κB and those in orange enriched in the IFN⍺/γ response, identified by pre-ranked gene set enrichment analysis using the MSigDB Hallmark collection; box representation is same as Fig. 1E. **C.** Cell cycle gene expression (representative patient IFN01, n = 97 baseline and 97 IFN⍺-treated HSCs, top). Frequencies of cells in S/G2/M phase as assessed in top subpanel at baseline (n = 11 samples) and at year 1 of IFN⍺ treatment (n = 9 samples). For IFN05 which has two IFN⍺ year 1 samples, the active IFN⍺ time-point was selected. P-values from likelihood ratio test of LMM with/without treatment status (**methods**). **D.** CXCR4 vs. cell cycle gene expression in HSCs before and after IFN⍺ treatment. Pie charts show frequencies of baseline versus treated cells in cell cycle-low, CXCR4-high and those in cell cycle-high, CXCR4-low populations. P-value from two-sided Fisher’s exact test. **E.** Heatmap showing results of the pre-ranked gene set enrichment analysis comparing baseline and during IFN⍺ treatment across HSC and progenitor subsets. Values show the sign of the normalized enrichment score (NES) multiplied by -log10(Adjusted P-value). **F.** Pre-ranked gene set enrichment analysis comparing before and after treatment with IFN⍺ in cDCPs, showing downregulation of TNF⍺ signaling via NF-κB and the leading-edge genes. Genes highlighted in blue represent those upregulated in the IGP versus IMP (**table S3**, Fig. 1E). **G.** Heatmap showing results of the pre-ranked gene set enrichment analysis comparing baseline and upon IFN⍺ treatment discontinuation in HSCs (combined HSC1 and HSC2 subsets from Fig. 1G) and HSC-IGs. Values show the sign of the normalized enrichment score (NES) multiplied by - log10(Adjusted P-value). HSC-IG (vs HSC1) module was calculated using net score of genes upregulated and downregulated in HSC-IG (**table S3**). **H.** Boxplot showing module expression for genes involved in TNF⍺ signaling via NF-κB at baseline and during IFN⍺ treatment in cDCs (left) and NK cells (right). P-value from likelihood ratio test of LMM with/without treatment status. **I.** Box plots showing HSC-specific IFN⍺-induced signature score in IFN⍺-treated HSCs and IGPs (**methods**). Score calculated using upregulated (left) or downregulated (right) genes (**table S8**). P- values from likelihood ratio tests of LMM with/without cell type.

To define the transcriptional perturbations by IFN⍺, differential expression analyses were performed between baseline and treated CD34^+^ cells, as a function of cell identity. We identified genes commonly regulated across multiple progenitor subsets upon IFN⍺ administration, including the canonical IFN⍺ genes, such as *ISG15, IFITM3, IFI6* and *EPSTI1* (**Fig. 3B**, **fig. S5B, table S8**). To test whether IFN⍺ induces human HSCs into cell cycle entry, as reported in mice (*9*), we examined the gene signatures for cell cycle phases shown to be an accurate assessment of cell cycle status (*44*). Indeed, HSC rates of cell cycle entry were enhanced upon IFN⍺ treatment (**Fig. 3C**, **fig. S5C-D)**. A positive regulator of HSC quiescence (*45*), *CXCR4*, was downregulated upon IFN⍺ therapy, suggesting that *CXCR4* downregulation may help coordinate HSC activation (**Fig. 3B**). We incorporated protein detection for CXCR4 (CD184) via GoT-IM and identified that IFN⍺ reduced surface CD184 expression (**fig. S5E)**. Consistently, CD184 downregulation was observed in HSCs in cell cycle and enriched in IFN⍺-treated populations and, conversely, HSCs with elevated CD184 expression were less likely to be in cell cycle and enriched in baseline samples (P = 1.5 x 10^-9^, Fisher’s exact test, **Fig. 3D**). In light of enhanced rates cell cycle entry of HSCs with abrogation of *Cxcr4* in mice (*45*), these data suggested that CXCR4 downregulation may play a role in permitting HSC cell cycle entry upon IFN⍺ exposure.

In the treated MkPs, *CD9* and *VWF*, closely associated with MkP differentiation (*46–49*), were downregulated (**fig. S5B, table S8**). We also observed a downregulation of *TGFB1* (**fig. S5B**), which encodes the pro-fibrotic cytokine TGFβ established as one of the main inducers of marrow fibrosis in patients with myelofibrosis (*50, 51*). We and others have shown that MPN megakaryocytes exhibit increased *TGFB1* expression (*16, 50, 52*), and thus downregulation of *TGFB1* by IFN⍺ indicates a potential mechanism of disease amelioration by IFN⍺. To ensure that the transcriptional changes we observed were specific to IFN⍺ therapy, we performed scRNA-seq on CD34^+^ HSPCs from individuals with *CALR*-mutated ET who were treated with hydroxyurea, at baseline and at one year of treatment, and observed no significant overlap between the genes differentially regulated by IFN⍺ and hydroxyurea (P = 0.599, Fisher’s exact test, **fig. S5F-G**).

To determine which canonical pathways may be modulated by IFN⍺, we performed gene set enrichment analysis using the Hallmark gene sets in the IFN⍺-treated versus baseline cells and identified an upregulation of the IFN⍺ signaling pathway across the cell subsets as expected (**Fig. 3E**, **table S9**). The analysis also confirmed the upregulation of cell cycle-related pathways (G2M check point and E2F targets) as well as MYC targets (**Fig. 3E**, **table S9**), corroborating a previous study that reported enhanced MYC protein expression during IFN⍺-induced cell cycle entry of mouse HSCs (*53*). Consistent with the downregulation of *TGFB1* gene itself, gene set enrichment analysis between baseline and IFN⍺-treated cells revealed a decrease in TGFβ signaling across several HSPCs, including downregulation of *THBS1* and *SERPINE1* (**Fig. 3E**, **fig. S5B, table S8,9)**. TGFβ signaling was particularly downregulated in the HSCs (**Fig. 3E**), as was observed in mice and associated with HSC exit from quiescence (*10*).

Furthermore, in contrast to the pro-inflammatory state of the IFN⍺-associated IGPs, we observed a downregulation of pro-inflammatory pathways, including the TNF⍺ signaling via NF-κB and inflammatory response pathways, across the stem and progenitor cells and especially pronounced in the cDCPs (**Fig. 3E-F**, **table S9)**. Downregulated genes in the TNF⍺ signaling via NF-κB pathway included AP-1 subunits, an NF-κB subunit *NFKB1*, and *IL1B*, which encodes the pro-inflammatory cytokine IL-1b (**Fig. 3F**, **table S8,9)**. Expression of *IL1R1* and *CXCL8* genes from the inflammatory response pathway were also downregulated (**table S8)**. Previously, IFN⍺ treatment in mice has yielded mixed results demonstrating either an upregulation (*10*) or downregulation (*54*) of TNF⍺ by IFN⍺. Our findings indicated that in human, isolated IFN⍺ exerts an overall anti-inflammatory response, consistent with a previous report demonstrating decreased *TNF* mRNA levels following IFN⍺ therapy in patient samples (*55*).

To determine whether the IFN⍺-regulated pathways may be partially retained in the HSCs after discontinuation of therapy, we compared HSCs from baseline to post-treated samples (IFN04, IFN05, IFN06) not on active IFN⍺ therapy (off therapy for ∼3-4 weeks). We observed a residual IFN⍺ response signature and a slight downregulation of the TNF⍺ signaling via NF-κB pathway in the post-therapy samples compared to the baseline, but intriguingly, the TGFβ signaling was upregulated in the post-treated HSCs compared to both baseline and actively treated HSCs (**Fig. 3G**, **fig. S5H**). These data suggested that following IFN⍺ exposure, HSCs actively upregulate the quiescence program, consistent with the report in mice that HSCs re-enter quiescence following activation by type 1 IFN (*10*). Furthermore, post-therapy HSC-IG upregulated the gene expression program that defined HSC-IG (versus HSC1, **Fig. 3G**, **fig. S5H**), suggesting the reinforcement of the inflammatory signature with prior inflammatory exposure. Consistently, the HSC-IG and IGP frequencies at baseline showed a trending increase with age (**fig. S5I**), and HSC-IGs exhibited a higher aging gene signature compared to the other HSCs (**fig. S5J**) (*56*). Altogether, these data implicated inflammatory neutrophil development as a feature of HSC memory – i.e., trained immunity, a field that has largely focused on monocyte development thus far (*57*).

We next determined whether the anti-inflammatory state propagated to the mature immune cells upon IFN⍺ therapy. The mature immune cells displayed a greater degree of heterogeneity in their response to IFN⍺ compared to stem and progenitors, indicating a cell type-specific response to IFN⍺ (**fig. S5K, table S10,11**). Notably, IFN⍺ did not induce cell cycle entry of mature immune cells (**fig. S5K-L**). Nonetheless, IFN⍺ therapy downregulated TNF⍺ signaling in innate immune cells, specifically classic dendritic cells and natural killer cells (**Fig. 3H**, **fig. S5K**). Immune cell composition also reflected this shift to an anti-inflammatory condition, with the expansion of regulatory T (Treg) cells (relative to other T-cell subsets), as well as a diminution of the pro-inflammatory CD16^+^ monocytes (within the monocytic compartment, **fig. S5M-N**) (*58, 59*). These data indicated that the anti-inflammatory effects of IFN⍺ on the stem and progenitor cells are likely compounded by the anti-inflammatory state of the bone marrow immune microenvironment.

Given the induction of IGP differentiation and inflammatory neutrophils with upregulation of AP-1 and NF-κB targets, these findings suggest that isolated IFN⍺ initiates both *anti-inflammatory* and *pro-inflammatory* states – both occurring in parallel within the same individual’s hematopoiesis. Consistently, we observed a significant overlap between the IFN⍺-downregulated (*IFN⍺^DN^*) genes and genes upregulated in the IGPs (*IGP^UP^*, **Fig. 1E**, **3B, table S3,8,** P = 1.6 x 10^-5^, hypergeometric test). The inverse, i.e., a significant intersection of *IFN⍺^UP^* genes and *IGP^DN^* genes, was also observed (P = 1.73 x 10^-69^, hypergeometric test, **table S3,8**). Coherently, the HSC-specific *IFN⍺^UP^* gene signature (displayed in **Fig. 3B**) was significantly downregulated in the IGPs compared to the HSCs while the *IFN⍺^DN^* genes were upregulated in the IGPs (with both cell groups under IFN⍺ treatment; **Fig. 3I**). Altogether these data indicated that IFN⍺ can precipitate opposing cell states within the same HSC population. While previously the heterogeneity of HSC cell states was linked with lineage outputs (*60, 61*), these data revealed that HSC heterogeneity may also mediate polarized anti-and pro-inflammatory responses to regulate specialized immune function.

### IFN⍺ potentiates lymphoid differentiation shift

To determine how the transcriptional remodeling by IFN⍺ may impact the hematopoietic differentiation trajectories, we computed the proportion of stem and progenitor subsets within the CD34^+^ compartment before and following IFN⍺ therapy. In addition to the expansion of the alternate IGP state, IFN⍺ also paradoxically induced a significant expansion of the lymphoid progenitors (**Fig. 4A-B**, **fig. S6A**). While the expansion of lymphoid progenitors was an unexpected finding as inflammatory cytokines have been demonstrated to induce myeloid priming (*62–64*), it was consistent with the downregulation of pro-inflammatory pathways, including those of the AP-1 subunits, associated with myeloid differentiation (*65*). In congruence with downregulation of *VWF* and *CD9* expressions, we observed diminutions of the megakaryocytic-erythroid lineage progenitors, including MkPs and erythroid progenitors (EP, **Fig. 4A-B**; paired analyses from serial samples in **fig. S6A**).

**Figure 4.**
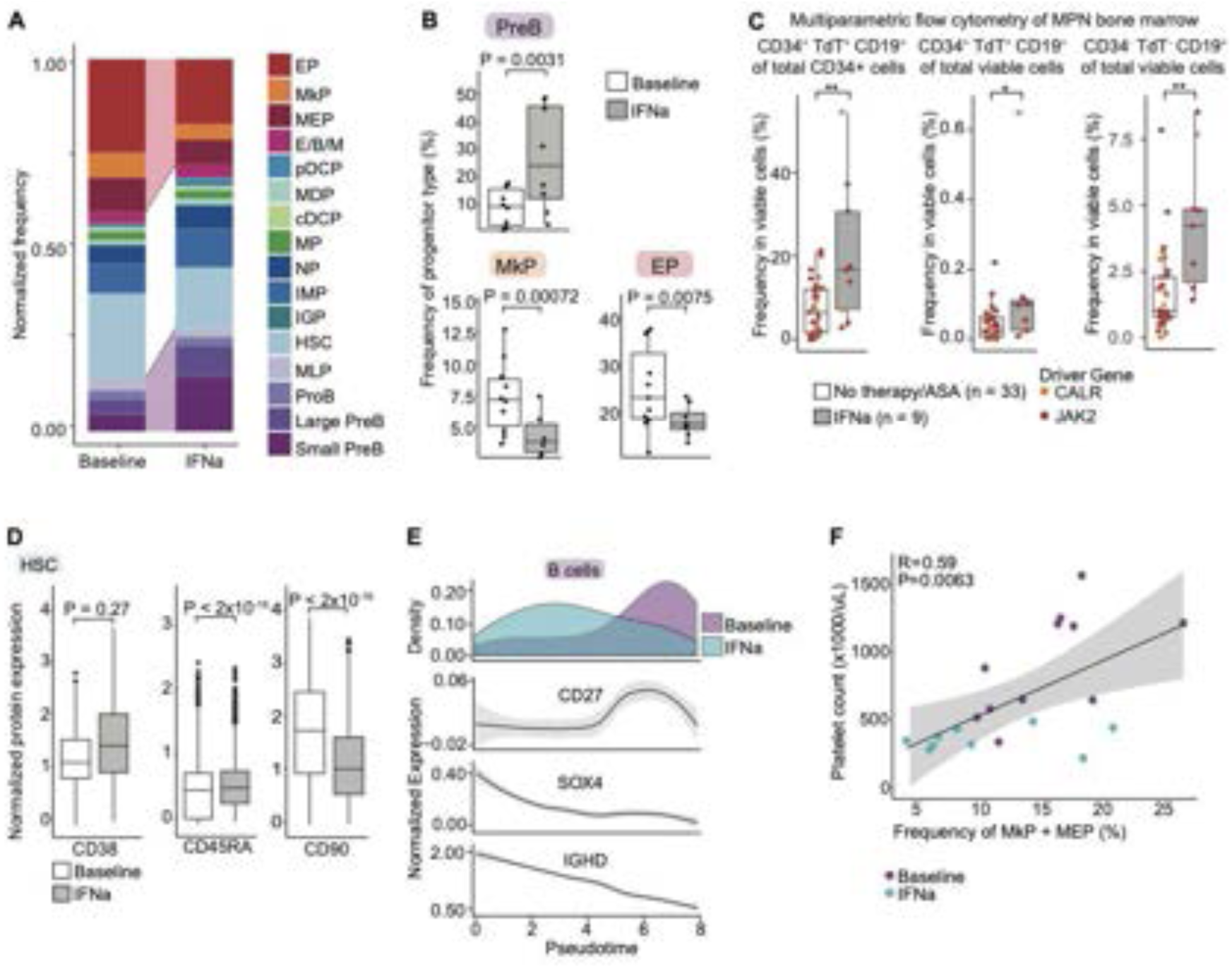
IFN⍺ induces IGP and lymphoid differentiation. **A.** Normalized cell type frequencies at baseline and IFN⍺ treatment. Cells from each sample were down-sampled to the same number (n = 500 cells from each sample, n = 11 baseline samples, n = 9 treated samples). For IFN02, IFN04, IFN05 and IFN08, which have two treated time-points, the time-point powered with greater number of cells was selected (also for panels **B, D**). **B.** Box plot showing normalized cell type frequencies at baseline and IFN⍺ treatment (n = 11 baseline samples, n = 9 treated samples). P-values from likelihood ratio test of linear-mixed modeling (LMM) with/without treatment status. **C.** Box plots showing cell frequencies of B-lymphoid progenitors and B cells from bone marrow of patients with early stage MPN with no treatment or aspirin only (ASA) or with IFN⍺ therapy (n = 33 and 9 samples, respectively), as determined by multiparametric flow cytometry (**table S12**). P-values from Wilcoxon rank sum test, two-sided. **D.** Box plots showing normalized protein expression in HSCs before and after treatment with IFN⍺. P-values from likelihood ratio tests of LMM with/without treatment status (**methods**). **E.** Gene expression of canonical B cell differentiation markers across immature and mature B cells in lymphoid development between baseline and IFN⍺-treated cells. **F.** Platelet counts versus frequencies of MEPs and MkPs. P-value from generalized linear model; Pearson correlation, shading denotes 95% confidence interval.

To validate the generalizability of the lymphoid expansion, we analyzed clinical multi-parametric flow cytometry data of bone marrow aspirates from patients with early-phase MPN with *JAK2* or *CALR* mutations (n = 33 samples without IFN⍺ exposure; n = 9 samples with IFN⍺ therapy). Indeed, we identified that the proportion of TdT^+^, CD19^+^ B-lymphoid progenitors within CD34^+^ HSPCs increased in IFN⍺-treated bone marrow compared to control (**Fig. 4C**, **S6B, table S12)**. We also observed an increased proportion of CD34^+^, TdT^+^, CD19^+^ cells of all viable cells analyzed, as well as an increase in the CD34^-^, TdT ^-^, CD19^+^ B-lymphocytes (**Fig. 4C**), providing evidence for an active lymphoid priming of HSCs leading to an expansion of the lymphoid progenitors. These findings were specific to IFN⍺, and not observed upon hydroxyurea therapies (**fig. S6B, table S12**). In support of lymphoid priming, HSCs exhibited an increased protein expression of CD45RA (used to identify multipotent lymphoid progenitors, MLP) after IFN⍺ treatment (**Fig. 4D**). CD90 was downregulated, consistent with the IFN⍺ effect of driving cell cycle entry and differentiation (**Fig. 4D**). Importantly, CD90, CD38 and CD45RA expressions were coherent with the transcriptional signature of HSCs following IFN⍺ treatment (**fig. S6C**), in contrast to mice progenitors that upregulate an HSC marker Sca-1 upon type 1 IFN treatment (*10*). Consistent with active differentiation bias toward B-lymphoid progenitors, the mature B cell compartment from IFN⍺-treated samples revealed a more immature B cell state compared to the baseline B cells (**Fig. 4E**), in the absence of proliferation of the CD34^-^ B cell compartment (**fig. S5K, S6D**). The major shifts in progenitor output by the HSCs suggested that the clinical improvement in the patients’ platelet count (despite variable molecular responses) may be due to the differentiation skewing away from the megakaryocytic to the lymphoid lineage. Consistent with this model, the proportions of MEPs and MkPs in CD34^+^ cells could help predict the patient’s platelet counts (P = 0.0063, generalized linear model, **Fig. 4F**). These data thus supported a model wherein the imbalance of hematopoietic differentiation landscape caused by somatic mutations in HSCs may be corrected by IFN⍺, as a mode of therapeutic efficacy in hematopoietic neoplasms.

### GoT-ATAC identifies transcription factor regulatory networks that govern IGP differentiation

Chromatin accessibility enables approximation of TF activity based on accessibility of the TF binding sites (*66–70*). Thus, to determine the regulatory networks that govern IFN⍺-induced modulation of inflammatory and differentiation states, we expanded upon GoT to capture somatic mutational status, chromatin accessibility and whole transcriptomes. We adapted the 10x Multiome platform that captures single-nuclei RNA-seq (snRNA-seq) and chromatin accessibility (snATAC-seq) to include somatic genotyping, i.e., GoT-ATAC (**Fig. 5A**). We applied GoT-ATAC to serial bone marrow CD34^+^ cells (n = 23,137 cells) from the clinical trial cohorts (n = 4 baseline, 3 IFN⍺-treated samples). As in GoT-IM, we incorporated time-point specifying barcoded antibodies (*71*) to combine serial samples from the same individuals into a single experiment to remove technical batch effects (**Fig. 5A**, **fig. S7A-E**). After we analytically segregated the baseline and IFN⍺-treated samples (**fig. S7E**), we clustered the cells across samples based on the transcriptomic and chromatin accessibility data and identified the same cell states identified by GoT-IM, including the IGPs (**fig. S8A-E, table S13**).

**Figure 5.**
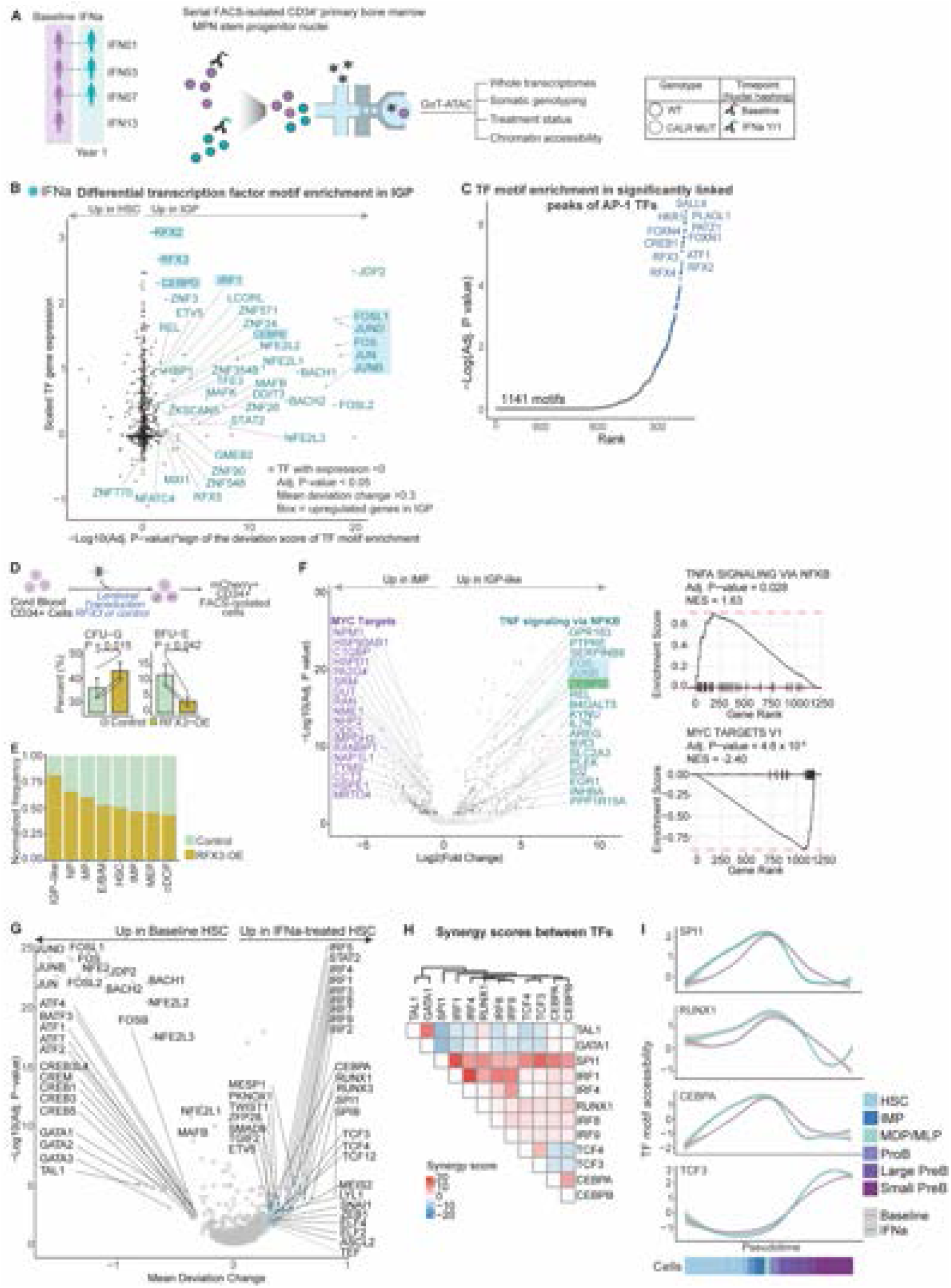
Regulatory networks of IGPs highlight PU.1 as the master regulator of IFN⍺-mediated remodeling of hematopoiesis. **A.** Representation of primary bone marrow samples at baseline and after IFN⍺ treatment (left). Schematic of GoT-ATAC (right). **B.** Motif enrichment and expression of transcription factors (TF) in the IGPs relative to HSCs from IFN⍺-treated samples. Normalized gene expression derived from GoT-IM data within HSCs and IGPs. P-values from Wilcoxon rank sum test, two-sided, with Benjamini-Hochberg FDR-correction (**methods**). **C.** Ranked TF motif enrichment of all positive regulatory peaks of the AP-1 members, relative to background peaks using the hypergeometric test (see **methods**). **D.** Schematic of lentiviral RFX3-overexpression (RFX3-OE) transduction experiment in CD34^+^ umbilical cord blood cells (UCB) (top). Normalized frequency of granulocytic (CFU-G) and erythroid (BFU-E) colonies grown via methylcellulose-based colony-forming unit (CFU) assay compared between control and RFX3-OE CD34^+^ UCBs (n = 3 independent experiments, bottom). **E.** Normalized frequency of RFX3-OE and control cells obtained from scRNA-seq data. **F.** Volcano plot showing genes differentially expressed (DE) between IMP and IGP-like cells from control and RFX3-OE subsets identified via linear mixed model (LMM) with/without cluster identity (left, **methods**). Highlighted are genes enriched in the TNF⍺ signaling via NF-κB (blue) and MYC targets (purple, box representation is same as Fig. 1E). Pre-ranked gene set enrichment analysis of TNF⍺ signaling via NF-κB pathway and MYC targets comparing control and RFX3-OE UCBs (right). **G.** Volcano plot showing TF motifs differentially enriched in IFN⍺-treated versus baseline HSCs (chromVAR). P-values from Wilcoxon rank sum test, two sided, with Benjamini-Hochberg FDR-correction (n = 6 serial samples from 3 individuals, see **table S13** for cell numbers, **methods**). **H.** Heatmap showing synergy scores between TFs as assessed by measuring the excess variability of accessibility for peaks with both TF motifs (*125*). **I.** TF motif accessibility across stem and progenitor subsets in lymphoid development between baseline and IFN⍺-treated cells.

As the IGPs were defined by a robust upregulation in gene expression of immediate early factors and RFX2/3, we determined whether these TFs showed increased activity based on chromatin accessibility of their binding sites. A differential TF motif enrichment analysis between IGPs and HSCs revealed that the same TFs showed enhanced accessibility, including the motifs of the AP-1 family (FOS, JUN, JUND, JUNB, FOSL1, FOSL2), CEBPB/D, and RFX2/3 (**Fig. 5B**, **fig. S8F, table S14**). Consistent with the overexpression of *IRF1* in IGPs, differential motif enrichment analysis also isolated IRF1 as the main interferon regulatory factor active in the IGPs. The differential motif analysis further revealed an upregulation of STAT2 and the proinflammatory REL of the NF-κB complex (**Fig. 5B**, **fig. S8F, table S14**), consistent with the observed gene expression upregulation of the NF-κB pathway (**Fig. 1E**, **fig. S3B**). While TF expression data are often sparse in scRNA-seq, these data demonstrated a high concordance of TF gene expression and their binding motif accessibility, suggesting a rapid induction of the IFN⍺ transcriptional regulatory program.

To determine which TFs upregulated the gene expressions of RFX2/3, we determined TF motifs present in the regulatory peaks that correlated with their gene expression (i.e., linked peaks analysis) (*72*). In the cis-regulatory region of RFX3, we identified the binding motifs of STAT2 and IRF1, as well as FOSB and PU.1, which were upregulated at the gene expression level in IGPs **(fig. S9A-B, table S15**). Likewise, the most significant regulatory regions for RFX2 included motifs for PU.1, KLF factors, NR4A1/2 and AP-1 subunit factors (**fig. S9A,C, table S15**). To determine the regulatory networks that govern the robust upregulation of the AP-1 subunits in the IGPs, we assessed for motif enrichment in the positive regulatory peaks for the AP-1 TFs in aggregate and identified RFX2-4 (which have the same binding motifs) as among the most significant TFs (**Fig. 5C**, **table S16**). Thus, RFX2/3 and AP-1 TF groups positively regulated the expression of the other, synergizing IGP development. These data provided evidence for a model wherein HSC-IG with elevated expression of RFX2/3 and AP-1 subunits and other immediate early response factors were primed toward a robust transcriptional program for IGP differentiation upon IFN⍺ signaling.

Furthermore, as other RFX members (i.e., RFX1 and RFX8) play key regulatory roles in MHC class II expression (*73–75*), we hypothesized that RFX2/3 may downregulate MHC class II in the IGPs. To test this, we determined the significantly linked peaks that negatively regulated *HLA-DRA* expression (**fig. S10A, bottom**) (*72*). We identified a distal regulatory region with four negative regulatory loci (**fig. S10A, inset**). These peaks included motifs for IRF1 and STAT2 as well as immediate response factors including AP-1 and KLF families, but not RFX2/3 (**fig. S10A, inset, table S17**). However, within the same negative regulatory region, we identified an IGP-specific peak that included the binding motif for RFX1-4, KLF factors and IRF1 (**fig. S10A, top, table S17**). These findings identified the immediate response factors, IRF1 and STAT2 as negative regulators of MHC class II genes across cell types, while RFX2/3 binding was specific for MHC class II downregulation during IGP differentiation. Consistently, surface HLA-DR was suppressed in the IGPs in the GoT-IM data (**fig. S10B**). Genes positively regulated by RFX2/3 included *MPO* and genes involved in cell cycle entry (**table S18**), suggesting that these factors play an essential role in activating the IGPs.

As *RFX3* was upregulated in the HSC-IG to IGP transition, we overexpressed *RFX3* in primary cord blood CD34^+^ cells via lentiviral transduction to determine whether RFX3 may regulate granulocytic differentiation. Methylcellulose-based colony forming unit assays of *RFX3* overexpressing (RFX3*-*OE) CD34^+^ cells revealed that *RFX3* overexpression expanded the granulocytic colonies (CFU-G) and diminished the erythroid colonies (CFU-E) compared to control-vector transduced CD34^+^ cells (**Fig. 5D**, **fig. S10C-D)**. To determine whether RFX3 may regulate neutrophil differentiation at the CD34^+^ HSPC stage, we performed single-cell RNA-seq of RFX3-OE CD34^+^ cells with control-vector cells and non-transduced cells. We identified an IGP-like progenitor state that was highly enriched for the RFX3-OE HSPCs (**Fig. 5E**, **fig. S10E-I**). Compared to the IMPs, this IGP-like progenitor state recapitulated the IGP signature including upregulation of AP-1 subunit TFs and *CEBPD* among other TNF⍺ signaling via NF-κB pathway genes, and downregulation of the MYC pathway (**Fig. 5F**, **table S19**). These data highlighted RFX3 as a key transcription factor regulator of IGP development.

### PU.1 underlies hematopoietic differentiation remodeling by IFN⍺

Furthermore, the GoT-ATAC data also confirmed the expansion of the lymphoid progenitors upon IFN⍺ treatment (**fig. S11A**). To determine the regulatory networks that governed the differentiation skewing by IFN⍺, we performed differential motif enrichment analysis between IFN⍺-treated and baseline HSCs. IFN⍺ enhanced the activities of STAT2 and several interferon regulatory factors in HSCs, in contrast to the specific activity of IRF1 in IGPs (**Fig. 5G**, **table S20**). IFN⍺ downregulated the motif accessibility of AP-1 TFs (**Fig. 5G**), consistent with the downregulation of AP-1 TF gene expression associated with TNF⍺ signaling via NF-κB (**Fig. 3B**). Moreover, the accessibility of the TWIST1 motif was enhanced after treatment (**Fig. 5G**), consistent with a previous report that identified TWIST1 as mediating the downregulation of TNF⍺ upon type 1 IFN treatment (*54*). Furthermore, the activity of TGFβ induced factor homeobox 2 (TGIF2), which inhibits TGFβ response genes, was also enhanced (**Fig. 5G**, **table S20**), highlighting TGIF2 as another key factor involved in the downregulation of the observed TGFβ signaling after IFN⍺ treatment.

Importantly, critical TFs involved in hematopoietic differentiation (*76*) were differentially regulated. Notably, motif accessibilities of PU.1 and RUNX1, essential for early lymphoid and granulo-monocytic differentiation (*76*), were enhanced by IFN⍺ (**Fig. 5G**, **table S20**). The activities of GATA1/2, reported to be negatively regulated by PU.1 (*76*), were downregulated as well as those of another critical megakaryocytic-erythroid lineage factor, TAL1 (*77*). These findings were consistent with the differentiation away from the megakaryocytic-erythroid lineages upon IFN⍺ treatment. Furthermore, the accessibilities of the motifs of the critical early B-lymphoid differentiation factors TCF3/4 were enhanced (*71, 78, 79*), highlighting TCF3/4 as TFs that govern the differentiation toward lymphoid progenitors by IFN⍺. CEBPA/B/D, essential for granulo-monocytic development (*80, 81*), were also upregulated in its motif accessibility (**Fig. 5G**, **table S20**). Importantly, the IRFs (*82, 83*), RUNX1 (*84*), CEBPA (*85*) and TCF3 (*86*) have been demonstrated to co-regulate target gene expression with PU.1, suggesting that enhanced PU.1 activities may have enhanced the activities of its co-regulating TFs, such as CEBPA and TCF3. To confirm these co-regulations in IFN⍺-remodeled hematopoiesis, we examined the synergistic activities of the different combinatorial TFs by measuring the excess variability of accessibility for peaks with both TF motifs (compared to peaks with one motif) (*69*). Indeed, PU.1 exhibited synergistic activities with IRF1, RUNX1, CEBPA and TCF3, while displaying antagonism with GATA1 and TAL1 (**Fig. 5H**). To determine the activities of these TFs upon lymphoid differentiation, we assessed the motif accessibilities of the TFs across the early and late progenitors. The activities of PU.1, RUNX1 and CEBPA were enhanced upon IFN⍺ signaling during the early stages of hematopoiesis and diminished with lymphoid development, whereas the activities of TCF3 were more pronounced in the later stages of development (**Fig. 5I**). These data highlighted PU.1 as a master regulator of IFN⍺-induced differentiation.

To determine whether IFN⍺ may directly upregulate the expression of PU.1, we assessed the regulatory peaks of PU.1 gene itself and identified the binding motifs of interferon regulatory factors (**fig. S11B, table S16**). Consistently, PU.1 gene expression increased upon IFN⍺ treatment in stem and early progenitor cells (**fig. S11C**). IFN⍺ treatment of a hematopoietic cell line (K562) in vitro upregulated PU.1 gene and protein expression (**fig. S11D-E**), consistent with a previous report (*87*), demonstrating that IFN⍺ directly upregulates the expression of PU.1. As PU.1 regulatory peaks also included CEBPA binding motifs, we overexpressed CEBPA in K562 cells and observed enhanced expression of PU.1 (**fig. S11F**). We further tested whether enhanced PU.1 may, in turn, co-regulate the expressions of canonical type 1 IFN targets. Overexpression of PU.1 in K562 cells with or without IFN⍺ treatment revealed that PU.1 overexpression led to upregulation of *IRF1* and *B2M* and downregulation of *ISG20* and *IFIH1*, as examples (**fig. S11G-H**). Altogether, these studies revealed that IFN⍺ signaling caused a global rewiring of TF regulatory activities, particularly through PU.1 for both lymphoid and IGP expansion, that underpinned the concerted transcriptional and differentiation remodeling.

### Somatic mutations modify the downstream effects of IFN⍺

Having established the overall effects of IFN⍺ on hematopoiesis, we next determined the differential effects of IFN⍺ on *CALR*-mutated versus wildtype stem and progenitor cells, specifically in relation to the effects of IFN⍺ on the clonal fitness of HSCs (**Fig. 6A**, **fig. S12A**). The high genotyping efficiency of GoT-IM enabled us to track the clone size precisely within the HSCs, revealing that IFN⍺ caused variable changes to the *CALR*-mutated clone size (**Fig. 6B**, **fig. S12B**), consistent with previous reports of bulk sequencing on peripheral blood samples (*12, 13, 15, 17*). Current models of inflammation-induced perturbation to clonal evolution build on the induction of cell cycle entry of clonal HSCs for enhanced fitness (*5*) or differentiation and depletion (*9, 12, 13*). We therefore determined whether differential upregulation of HSC cell cycle entry might predict clonal dynamics. Across the patients regardless of clonal dynamics, IFN⍺ induced greater rates of cell cycle entry of *CALR*-mutated HSCs compared to the wildtype counterparts (**Fig. 6C**, **fig. S12C**). These results indicated that the mutated HSCs were likely primed for a more robust proliferative response, compared to the wildtype counterparts, potentially due to the baseline cell cycle activity enhanced by the *CALR* mutations. In support of this model, IFN⍺ boosted the proliferation to a greater degree in the mutated cells compared to wildtype within the myeloid lineages, especially those of the megakaryocytic-erythroid, but not the lymphoid progenitor compartment—that is, restricted to the progenitor subsets in which the *CALR*-mutation caused enhanced proliferation at baseline (*16*) (**fig. S12C**). We orthogonally validated that IFN⍺ effected greater proliferative rates in the myeloid HSPCs by performing multiplexed *in situ* fluorescent imaging of bone marrow paraffin-embedded sections from patients with *CALR*-mutated MPN with or without IFN⍺ treatment (n = 5 without IFN⍺, n = 4 with IFN⍺, **Fig. 6D**, **fig. S12D**). We determined the protein expression of CD34, CD117, CD38, Ki67 and mutated CALR (*88*) to assess the frequencies of CD34^+^, CD38^+^, CD117^+^ myeloid progenitors that express Ki67, a gold standard of cell cycle entry (*89*). The availability of a mutated CALR-specific antibody enabled us to identify that the frequency of cycling mutated myeloid progenitors was higher compared to that of wildtype with or without IFN⍺ exposure (**Fig. 6D**). Overall, these findings provided evidence that IFN⍺ enhanced the cell cycle entry of the HSPCs to the degree predetermined by baseline priming. Thus, while enhancing absolute rates of cell cycle entry across the stem cells, the relative difference in cell cycle rates between mutated and wildtype HSCs at baseline were preserved upon IFN⍺ exposure. These data indicated that inflammation-induced cell cycle entry rates may be decoupled from clonal dynamics.

**Figure 6.**
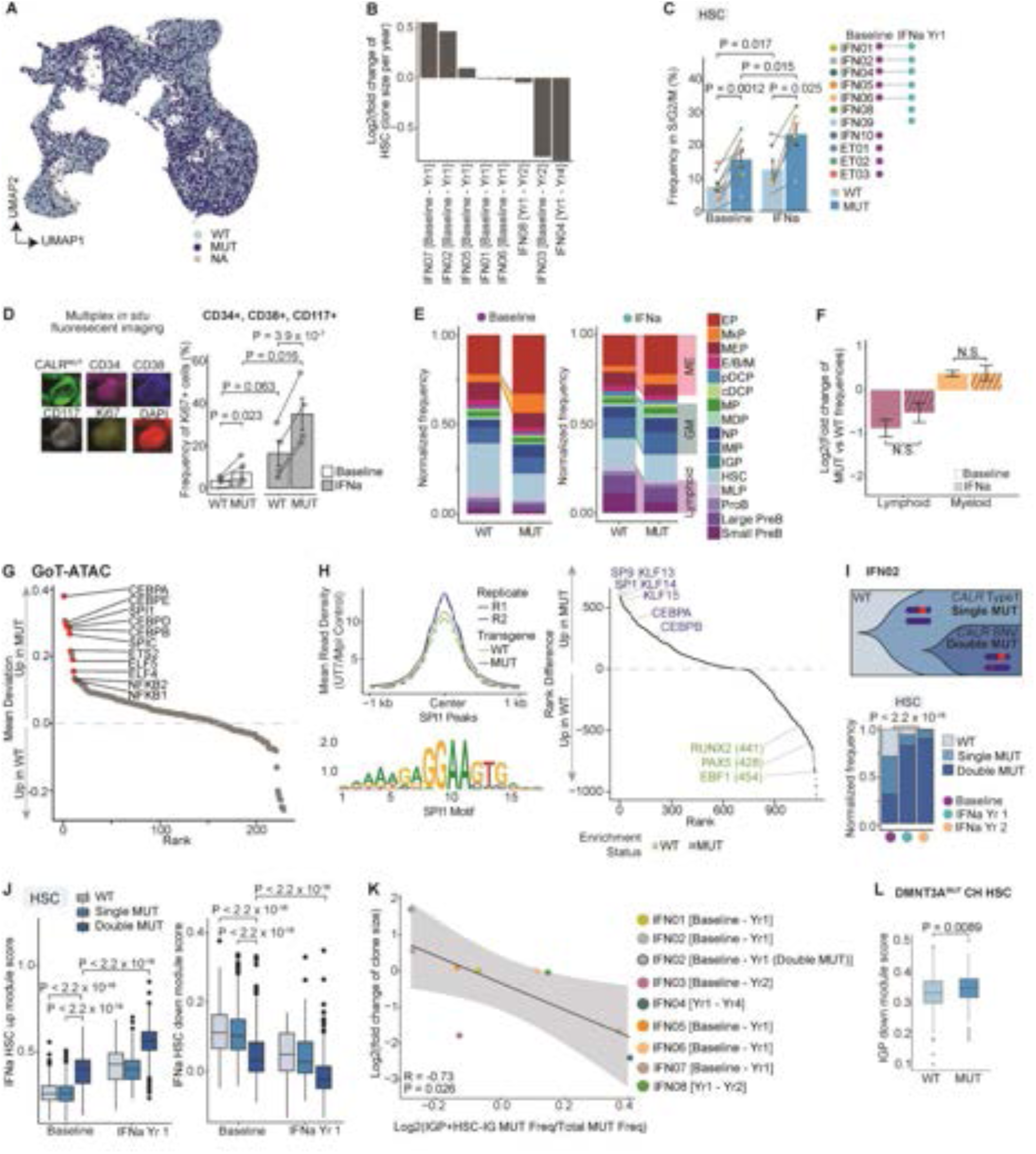
IFN⍺ modulates clonal dynamics via clonal HSC differentiation into IGP differentiation. **A.** UMAP of CD34^+^ cells (n = 65,452 cells) with mutation status highlighted for wildtype (WT; n = 21,354), *CALR-*mutant (MUT; n = 25,529) or unassigned (NA; n =18,569) cells. **B**. Bar plot showing HSC clone size changes at baseline and after IFN⍺ treatment. **C.** Bar plots showing frequencies of cells in G2/M/S phase as assessed in Fig. 3C (n = 11 baseline and 9 IFN⍺-treated, year 1 samples). P-values from likelihood ratio tests of linear mixed model (LMM) with/without treatment status (for comparisons between treatment) or mutation status (for comparisons between genotypes). **D.** Multiplex *in situ* fluorescence imaging of bone marrow biopsy sections from MPN patients with (n = 4) or without (n = 5) treatment with IFN⍺ (**methods**). Representative images of protein markers (top). Bar plots showing frequencies of proliferating myeloid cells as assessed by Ki67 expression (bottom). P-values from likelihood ratio test of LMM with/without treatment status or mutation status. **E.** Normalized cell type frequencies of WT versus MUT cells at baseline (n = 11 samples, left) and following IFN⍺-treatment (n = 9 samples, right). Cells down-sampled to 500 for each sample. **F.** Fold change of normalized cell frequency of MUT versus WT cells in the progenitor groups at baseline and following IFN⍺ treatment. **G.** Transcription factor (TF) motif enrichment in MUT versus WT stem and early progenitor cells (HSCs, IMPs, MDPs/MLPs and MEPs) at baseline. TFs differentially regulated by IFN⍺ were tested (**table S20**). Red = P < 0.05. P-values combined using Fisher’s combined test (**methods**). **H.** Density plot showing PU.1 peak count in WT and MUT cells from the control group of megakaryoblastic cell line expressing TPO receptor (UT7-MPL) with wildtype or mutant *CALR* transgenes (n = 2 independent experiments, left). Differential TF motif enrichment between PU.1 binding sites in WT IFN⍺-treated and MUT IFN⍺-treated UT7-MPL cells (right, **methods**). Analysis conducted with HOMER. TFs enriched in differentially accessible PU.1 peaks for MUT and WT cells are highlighted in blue and green respectively. **I.** Schematic of clonal structure of HSCs from patient IFN02 (top). Bar plot of normalized mutant cell frequencies across treatment time-points (bottom). P-value from pairwise Fisher’s exact test. **J.** Box plot showing HSC-specific IFN⍺-induced signature score in HSC clones at baseline and after one year of treatment. Scores calculated using upregulated or downregulated genes (left and right panels, respectively, **methods**). P-values from Wilcoxon rank sum test, two-sided. **K.** Clone size change with IFN⍺ treatment versus difference in normalized frequency of MUT HSC-IGs and IGPs out of all MUT cells. P-value from F-test, Pearson correlation. Shading denotes 95% confidence interval. **L.** Box plots showing IGP-specific downregulated signature score in wildtype and *DMNT3A*-mutant HSCs from individuals with clonal hematopoiesis (CH) (n = 4 samples, no. cells = 1316 wildtype, 529 mutated cells) (*42*). P-values from likelihood ratio test of linear mixed model with/without mutation status.

To determine whether IFN⍺ exposure may preserve other key features of *CALR*-mutation effects, we compared the gene expression profiles between wildtype and mutated cells, at baseline and following IFN⍺ therapy. Intriguingly, we observed the same differentially expressed pathways between the IFN⍺-treated mutated and wildtype HSCs, including the unfolded protein response (UPR) (*16, 90, 91*), likely due to a heterozygous loss of the wildtype *CALR* that encodes a critical chaperone protein (**fig. S12E-F, table S21-24**). IFN⍺ globally remodeled the differentiation toward lymphoid development, reducing the overall megakaryocytic bias in the mutated cells (**Fig. 6E**). Indeed, the frequencies of mutated MkPs and MEPs better predicted the patients’ platelet counts compared to overall frequencies of MkPs and MEPs (**fig. S12G**). However, the lymphoid expansion was constrained in the mutated compartment due to the relative expansion of the granulo-monocytic and megakaryocytic-erythroid progenitors compared to wildtype HSPCs (consistent with the differential upregulation of cell cycle entry in the mutated myeloid progenitors, **Fig. 6E-F**, **fig. S12C**). Furthermore, computation of the mutant cell frequency across the progenitor subsets before and after treatment revealed that the enrichment of the mutated cells in the myeloid compartments did not change following IFN⍺ treatment (**fig. S12H**). Altogether, these data revealed that phenotypic responses to IFN⍺ are constrained (e.g. lymphoid differentiation) or amplified (e.g. cell cycle entry) by somatic mutations, such that cell state distinctions between the mutated and wildtype cells are preserved upon IFN⍺ signaling.

We therefore hypothesized that *CALR*-mutations may alter the chromatin state of the binding sites of IFN⍺-regulated TFs and thereby modulate their activities following therapy. We tested differential TF motif enrichment via GoT-ATAC between the mutated versus wildtype stem and early progenitors at baseline (**Fig. 5A**). The binding sites of NFKB1/2 were enhanced in the mutated cells, consistent with our previous report of the gene set enrichment of the NF-κB pathway in the *CALR*-mutated early HSPCs (*16*). We also identified that the chromatin accessibility of PU.1 and CEBPA were increased in the mutated cells (**Fig. 6G**, **table S25**). These data suggested that *CALR* mutations alter the chromatin state of key lineage specifying TF binding sites, skewing the lineage-modulating activities of IFN⍺. Specifically, these data indicated that IFN⍺-induced PU.1 activities may be skewed toward granulo-monocytic (versus lymphoid) development via enhanced PU.1 and CEBPA co-activity in the mutated HSPCs.

To directly assess the impact of differential PU.1 activity due to the mutation status on modulating the effects of IFN⍺, we performed a chromatin binding assay (CUT&RUN) (*92*) for PU.1 in UT7 cell lines expressing MPL (thrombopoietin receptor) and either the mutant *CALR* (type 1, L367Tfs*46) or wildtype *CALR* transgene (*93*) treated with IFN⍺ (**fig. S13A, table S26**). We observed that PU.1 binding sites were enriched in the mutated cells compared to wildtype, consistent with the GoT-ATAC data and the myeloid bias induced by the *CALR* mutation (**Fig. 6H**, **left**). Following IFN⍺ treatment in vitro, we observed co-enrichment of PU.1-bound peaks with IFN⍺-specific transcription factor motifs, including IRF4/8, which are known to cooperate with PU.1 in the Ets-IRF composite elements (EICE) to mediate lymphoid and myeloid differentiation (**fig. S13B, table S27**) (*83, 94, 95*). We observed that IFN⍺ enhanced PU.1 binding at distal regulatory regions (**fig. S13C**), the regions that regulate the differential commitment to the lymphoid versus monocytic lineages by PU.1 (*96*). In these distal peaks, PU.1 binding sites were enriched for CEBPA/B co-binding sites in the *CALR* mutated cells compared to the wildtype (**Fig. 6H**, **right, fig. S13D, table S28**). These data demonstrated that *CALR* mutations enhance PU.1 binding activities and alter the preferential cooperating TF partners of PU.1. Overall, these results suggested that somatic mutations may alter the chromatin state of TF regulatory regions, which in turn modulate the downstream effects of inflammatory activation.

### IFN⍺ regulates clonal fitness of HSCs via IGP development

While IFN⍺ induced cycling of *CALR*-mutated HSCs at a higher frequency than wildtype HSCs consistently across patient samples, clonal dynamics following therapy was heterogeneous (**Fig. 6B**). To investigate other factors that may contribute to the effects of IFN⍺ on clonal dynamics, we examined an unusual case in which the mutated clone expanded substantially with IFN⍺ therapy in patient IFN02 (**Fig. 6B**). Interestingly, the HSPCs from patient IFN02 harbored a subclone with another mutation in *CALR*, a single nucleotide variant in the *CALR* allele (M131I, predicted to impact protein structure (*97*)) which is trans to the MPN-causing frameshift mutation (**fig. S14A**). At baseline, the double mutant clone remained subclonal to the dominant clone with the single canonical *CALR* mutation (**Fig. 6I**, **fig. S14B**). Upon IFN⍺ therapy, the double mutant clone overtook the neoplasm and the overall stem cell population, even though the double mutant clone did not exhibit significant difference in cell-cycle entry rates compared to single mutant HSCs at baseline (**Fig. 6I**, **fig. S14C**). The additional insult to CALR activities in the double mutant clone effected a greater UPR activation compared to single mutant HSCs, as expected (**fig. S14D-E, table S29-30**). Surprisingly, however, the predominant signatures of the double mutant HSCs compared to other HSCs at baseline were an upregulation of IFN response genes and decreased TNF⍺ and TGFβ signaling (**fig. S14D-E, table S29-30**), thus recapitulating the HSC response to extrinsic IFN⍺ and supporting a potential causal link between UPR and IFN activation (*98*). Consistently, the double mutant HSCs exhibited increased expression of the HSC-specific *IFN⍺^UP^* gene signature (**Fig. 3B**) at baseline and even higher expression upon IFN⍺ therapy (**Fig. 6J**, **left**). Similarly, the *IFN⍺^DN^* gene signature was downregulated in the double mutant clone at baseline and to a greater degree following treatment (**fig. 6J**, **right**). Further, unbiased clustering and dimensional reduction revealed that the double mutant clone at baseline clustered with the treated HSCs rather than with the other baseline HSCs (**fig. S14F-G**). As the double mutant HSCs share the same microenvironment as the wildtype and single mutant HSCs, the striking similarity of the intrinsically activated IFN⍺ signaling in the double mutant cells at baseline to the extrinsic IFN⍺ effects indicated that the predominant IFN⍺ signaling signatures observed in the IFN⍺-treated HSCs may be largely direct rather than secondarily mediated by the effects of IFN⍺ on other cell types. These findings also highlighted a genotype-specific modulation of HSC fitness by IFN⍺.

As the *IGP^UP^* signature has a significant overlap with the *IFN⍺^DN^* gene signature (including the immediate early response TFs), the significant downregulation of the *IFN⍺^DN^* gene signature of the double mutant cells suggested that the double mutant HSCs may be resistant to IGP development. Consistent with this hypothesis, the double mutant HSCs expressed lower *IGP^UP^* and higher *IGP^DN^* signatures (**fig. S14H**). Altogether, these data raised the hypothesis that the resistance of clonal stem cells to differentiate into the IGPs may dictate its clonal fitness upon IFN⍺ signaling. We therefore computed the fold change in clone size as a function of the difference in the frequencies of mutant cell frequency within the HSC-IG and IGPs relative to that of the total CD34^+^ compartment (**Fig. 6K**). Indeed, the propensity of clonal HSCs to differentiate into IGPs could model clonal dynamics (**Fig. 6K**). These findings identified the alternate IFN⍺-specific route of differentiation into IGPs as an avenue to perturb clonal dynamics, while perturbations to the existing programs did not lead to a net difference in fitness between mutated and wildtype HSCs. To test the generalizability of this model, we examined the *IGP^DN^* gene signatures in *DNMT3A*-mutated HSCs from individuals with clonal hematopoiesis (*42*), as *DNMT3A*-mutated clones are resistant to IFN⍺, frequently expanding upon treatment (*15*). Consistently, the *DNMT3A*-mutated HSCs demonstrated increased expression of the *IGP^DN^* gene signature compared to the admixed wildtype HSCs (**Fig. 6L**), suggesting that the *DNMT3A*-mutated HSCs may be resistant to IGP development. These results therefore support a unified model of clonal dynamics wherein the differential propensity of HSC clones to differentiate into the IGPs determines the clonal composition of the stem cell niche under inflammation.

## Discussion

Here, our studies deconvoluted the pleotropic effects of IFN⍺ in reshaping the differentiation trajectories of human HSCs for normalization of blood counts and modulation of clonal dynamics in myeloproliferative neoplasms. The single-cell multi-omics methods applied to serial sampling enabled us to track clonal evolution over a treatment period together with differential phenotypic remodeling of mutated versus wildtype stem and progenitor cells. We identified that the underlying somatic mutations both amplified the IFN⍺ effects (e.g., cell cycle entry) and constrained others (e.g., lineage skewing). In this way, the relative distinctions between the mutated and wildtype HSPCs were preserved, despite the global remodeling of hematopoiesis. These findings suggested that the relative fitness of clonal HSCs via programs already activated at baseline also remained constant. Thus, differential HSC activation into cell cycle entry alone could not fully explicate the inflammation-induced changes in clonal dynamics.

Instead, we identified an IFN⍺-specific alternate route of differentiation directly from HSCs that predicted clonal dynamics based on priming of clonal stem cells to differentiate into the IGP state, enabling resisting HSC clones to dominate the stem cell niche. Furthermore, identification of the IGP population highlighted an intriguing phenomenon in human HSCs: induction of the IGP differentiation through upregulation of the pro-inflammatory AP-1 and NF-κB activities indicated that IFN⍺ potentiates a pro-inflammatory as well as an overall anti-inflammatory cell states with downregulation of the same AP-1 and NF-κB activities within the same hematopoietic differentiation program. Differential expression levels of *RFX2/3* and immediate early response programs within the HSC populations were highlighted in our data as mediating the polarized response to IFN⍺, revealing a key functional relevance of HSC heterogeneity. These data suggested that HSC-IG with elevated *RFX2/3* and immediate early response gene expressions were poised to a precipitous pro-inflammatory response to stimuli, implicating these HSCs in innate immune memory or trained immunity, i.e. an adapted innate immune response due to a prior inflammatory activation of HSCs (*99, 100*). Trained immunity may thus serve as a non-genetic modifier of IGP priming, reflecting the interpatient variabilities in the rates of IGP development by the *CALR*-mutated stem cells. Notably, these results were highly concordant with a recent report of inflammatory memory HSCs that have significant overlaps with HSC-IG in gene expression and a similar retention of the inflammatory phenotype following resolution of active inflammatory stimulation (*56*). In this study, downregulation of the inflammatory signature by clonal stem cells in the context of CH was also associated with clonal fitness (*56*). These findings therefore highlighted a novel connection between trained immunity and clonal dynamics, further shedding insights into the impact of age-related inflammation to clonal expansion and malignant transformation.

Furthermore, Type 1 IFN, the first FDA-approved immunotherapy (*101*), remains an effective treatment and is being tested across various autoimmune (*102, 103*), cancer (*104*) and infectious disease contexts (*105, 106*), including COVID-19 (*107, 108*). In the context of MPN, IFN⍺ is the only agent to consistently to deplete clonal stem cells. While previous studies have not detected improvement in disease progression in patients treated with IFN⍺ (*20, 109*), large clinical trial cohorts would be required to detect significant differences between the IFN⍺ and control arms, as the rates of disease progression are low in patients with ET. Moreover, our identification of the HSC-specific IFN⍺ response, which includes IFN⍺/γ response upregulation and downregulation of the targets of TNF⍺ and TGFβ signaling pathways, may help clarify its therapeutic efficacy in other disease contexts beyond MPN. In mouse studies, IFN⍺ has been demonstrated to modulate TNF⍺ expression (*110*) with data suggesting either an upregulation (*10*) or downregulation (*54*) of TNF⍺ by IFN⍺. Thus, our studies clarify that the predominant effect of IFN⍺ in human HSPCs is the downregulation of downstream pathways involved in TNF⍺ signaling via AP-1 and NF-κB, consistent with amelioration of the inflammatory state in MPN and multiple sclerosis upon type 1 IFN administration (*17, 107*). It also suggested that individuals with deficiencies in type 1 IFN may have exhibited exaggerated response to COVID-19 infections (*111, 112*) due to the inability to counterbalance the pro-inflammatory effects of the other cytokines. Coherently, *TNF* expression was demonstrated to be decreased in patients with MPN who received IFN⍺ treatment (*55*). The pro-fibrotic TGFβ signaling was also broadly downregulated across progenitor subsets, through a coordinated downregulation of the *TGFB1* gene itself and upregulation of the TGFβ signal inhibiting TGIF2 activity. In this way, IFN⍺ downregulated two key cellular programs involved in the MPN-associated pathology, potentially underlying improved disease states. Moreover, as NF-κB and TGFβ signaling are both implicated in HSC quiescence (*113–116*), the anti-inflammatory effects of IFN⍺ were also linked with HSC exit from quiescence. The robust upregulation of TGFβ signaling in the HSCs following resolution of an acute IFN⍺ exposure supported data in mice of re-entry into quiescence to protect from HSC depletion following inflammation-induced cell cycle entry (*10*).

Another major finding in our work was the remodeling of hematopoietic differentiation toward the lymphoid lineage by IFN⍺. While various differentiation skewing by IFN⍺ has been reported in mice (*117–119*), the interpretation of the results is complicated by the alteration of the HSC-marker Sca-1 upon IFN⍺ exposure in mice (i.e. induction of Sca-1 in the CMPs, GMPs and MEPs, resulting in their inclusion within the Sca-1^+^, Kit^+^ HSC/multipotent progenitor compartment) (*10*). As other inflammatory cytokines, such as TNF⍺, IL-1, and IFNγ, have been demonstrated to induce granulo-monocytic differentiation (*62–64, 120, 121*), IFN⍺ presents as a unique cytokine among the inflammatory milieu to balance the granulo-monocytic differentiation with its positive regulation of B-lymphoid differentiation as another mode of dampening the pro-inflammatory response.

The reshaping of the differentiation landscape by IFN⍺ provided a novel model of therapeutic efficacy in patients with myeloid neoplasms. As MPNs are primarily the result of a defect in homeostatic hematopoietic development, due to an abnormal differentiation skewing and proliferation of the myeloid lineages, IFN⍺-induced lymphoid differentiation may serve to balance the differentiation landscape. IFN⍺ reshaped the major bifurcation divide in MPN, that is, from the JAK2/STAT5-mediated bifurcation at the myeloid (i.e., granulo-monocytic and megakaryocytic-erythroid) versus lymphoid commitment, to the PU.1-mediated bifurcation at granulo-monocytic and lymphoid versus megakaryocytic-erythroid commitments. These two models have been described based on the identification of stem progenitor cells that produce myeloid (*122*) or lymphoid lineages (*123*) versus granulo-monocytic-lymphoid lineages (*124*). Our data provide evidence for a plastic differentiation hierarchy in which PU.1-mediated production of granulo-monocytic and lymphoid lineages at the expense of megakaryocytic-erythroid lineages are prioritized in perturbed settings, such as infection. The ability of IFN⍺ to directly upregulate PU.1 expression suggested that the differentiation modulating effects of IFN⍺ may be largely direct. These findings are consistent with the landmark study from Essers *et al.* that elegantly demonstrated the ability of IFN⍺ to directly activate HSCs into cell cycle entry, via an in vivo cell mixing study in which only a small minority of the bone marrow cells harbored intact IFN⍺ receptors (*9*). The cell cycle entry rate was further enhanced with increasing frequency of bone marrow cells with intact IFN⍺ signaling, suggesting that IFN⍺ effects are both directly and indirectly mediated.

Overall, these studies revealed the pleiotropic modes of therapeutic efficacy of IFN⍺ and principles of clonal dynamics upon inflammatory activation. Importantly, our work motivates the development of novel therapeutic strategies to deplete clonal stem cells by enhancing their differentiation rates into the IGP state upon IFN⍺ therapy, a strategy that may be generalizable across myeloid neoplasms and clonal hematopoiesis.

## Acknowledgments

We thank patients who participated in the MPN-RC-111/2 clinical trials and patients at WCM, who generously donated samples that enabled this research. This work was enabled by the Weill Cornell Epigenomics Core, Genomics Core and Flow Cytometry Core. We thank Dr. Matthew Greenblatt and Dr. Steven Josefowicz (Weill Cornell Medicine) and Dr. Serine Avagyan (University of California, San Francisco) for a critical review of the manuscript. We also thank the funding sources that supported this work:

F30 Predoctoral Fellowship from the NHLBI of the National Institutes of Health (F30HL156496, ND)

Medical Scientist Training Program grant from the National Institute of General Medical Sciences of the National Institutes of Health under award number T32GM152349 to the Weill Cornell/Rockefeller/Sloan Kettering Tri-Institutional MD-PhD Program (DI, ND)

National Institute of Health grant (P01CA108671, RH)

Burroughs Wellcome Fund Career Award for Medical Scientists (ASN)

National Institutes of Health Director’s Early Independence Award (DP5 OD029619, ASN) National Heart Lung and Blood Institute (R01 HL167139, ASN),

Starr Cancer Consortium (ASN)

## Author contributions

Conceptualization: ASN, RH

Methodology: ASN, RH, SP, DAL, PS, EW, LM

Experimental investigation: DI, SR, OS, MSS, RC, NO, MT, MQ, GF

Analytical investigation: ASN, CL, AKM, AXX, SM, NP, MO, AD, PS, PC, HEK, ND, TA, ACD

Funding acquisition: ASN, RH

Project administration: ASN, RH

Resources: ASN, RH, RW, DC, FCT, GAZ, EW, BM

Supervision: ASN, RH

Writing – original draft: ASN, CL, DI

Writing – review & editing: ASN, RH, CL, DI, OS, SR, MT, MSS

All authors reviewed and approved the manuscript.

## Data and Code Availability

Data generated in this study will be available upon publication.

## Corresponding author

Anna S. Nam.

**Correspondence and requests for materials** should be addressed to A.S.N.

## List of Supplementary Materials

Materials and Methods Figs. S1 to S14

Tables S1 to S31 (Tables S1-S31 provided as xlsx files)

List of references only cited in Supplementary Materials: 131-163

## Materials and Methods

### Patient samples

The study was approved by the local ethics committee and by the Institutional Review Board (IRB) of Weill Cornell Medicine. The study was conducted in accordance with the Declaration of Helsinki protocol, and all patients provided informed consent. Cryopreserved bone marrow mononuclear cells were obtained from patients with *CALR*-mutated essential thrombocythemia (ET) treated with weekly pegylated IFN-alfa2a during clinical trials MPN-RC-111 (NCT01259817) (*19*) and MPN-RC-112 (NCT01258856) (*20*). Samples used for CD34^+^ cell analysis include 8 baseline and 13 treated samples (3 of which were collected post-IFN⍺ discontinuation) collected from 10 individuals for GoT-IM. Three baseline ET samples (ET01-ET03) from our previous work (*16*) were included as additional baseline controls with a similar distribution of age, gender, and disease status for treated samples that did not have a paired baseline control. For GoT-ATAC, n = 4 baseline and n = 3 treated samples were included from the clinical trials. For GoT-IM of CD34^-^ mature cell analysis, 5 baseline and 10 treated samples (2 of which were collected following IFN⍺ discontinuation) were collected from 7 individuals. Two hydroxyurea-treated peripheral blood samples from *CALR*-mutated ET patients from MPN-RC-112 were included for scRNA-seq of CD34^+^ cells. **Table S1** includes detailed clinical and sample information collected according to clinical trial protocols.

### Cell preparation

Cryopreserved bone marrow mononuclear cells were thawed and stained using standard procedures with the surface antibody CD34-PE (clone AC136, dilution 1:50, Miltenyi Biotec) and DAPI (Sigma-Aldrich), according to manufacturer’s protocol. To eliminate experimental batch effects, cells were labelled simultaneously with hashing antibodies with time-point-identifying barcodes as described (*21*) using Hashtag Antibodies 1-6 (TotalSeq-A, clone LNH-94, BioLegend) for GoT-IM and Anti-Nuclear Pore Hashing Antibodies 9 and 10 (Clone LNH-94, BioLegend) for GoT-ATAC. To link cell identities to expression of cell surface proteins, cells were also incubated with CITE-seq antibodies (*22*) according to manufacturer protocol (TotalSeq-A, BioLegend, see **table S31** for information on antibodies). For IFN01-IFN10 and IFN13, cells were subsequently sorted for DAPI^−^ and CD34^+^ cells to isolate CD34^+^ populations. For IFN01, IFN02, and IFN09, we sorted for DAPI^−^ and CD34^−^ cells to isolate mature bone marrow cells. For IFN05, IFN08, IFN11, and IFN12, CD34 expression by side scatter was used to enriched for granulocytes. CD34^−^, high side-scatter (SSC) was used to sort for granulocytes, medium SSC to enrich for monocytes/DCs, and low SSC to enrich for lymphocytes. Mature cell populations were then pooled at approximately equal ratios for granulocytes, monocytes/DCs, and lymphocytes for each sample. All FACS sorting was completed using BD Influx at the Weill Cornell Medicine flow cytometry core.

### GoT-IM

To simultaneously capture genotyping data and whole transcriptomic data, Genotyping of Transcriptomes (GoT) was performed by adapting the 10x Genomics platform as previously described (*16*). FACS-sorted CD34^+^ cells for each time-point from the same individual were pooled. The standard 10x Genomics Chromium 3′ (v.3.1 chemistry) was implemented according to the manufacturer’s recommendations up to the cDNA amplification step (10x Genomics, Pleasanton, CA). After cDNA amplification and SPRI bead (Beckman Coulter) cleanup, 10x scRNA-seq and ADT/HTO libraries were generated as recommended. A portion of the cDNA was used for somatic genotyping as previously described (*16*). Briefly, to capture the somatic genotypes of cells, cDNA was amplified with a locus-specific amplification (10-16 PCR cycles), using the generic forward SI-PCR primer and a locus-specific reverse primer for the *CALR* mutations (see **table S31** for primer sequences). The amplified locus-specific cDNAs are then cleaned using SPRI purification to remove unincorporated primers. Finally, the targeted amplicon libraries are generated through a PCR performed with a P5 generic forward PCR primer together with an RPI-x primer (**table S31**). The targeted amplicon libraries were spiked into the remainder of the gene expression and immunophenotyping libraries to be sequenced together on a NovaSeq (Illumina, San Diego, CA). The cycle settings were as follows: 28 cycles for Read 1, 90 cycles for Read 2, 10 cycles for i7 sample index and 10 cycles for i5 sample index.

### GoT-IM scRNA-seq data processing, alignment, cell-type classification and clustering

For single-cell GoT-IM data from IFN01-IFN10 patient CD34^+^ samples and from IFN01, IFN02, IFN05, IFN08, IFN09, IFN11 and IFN12 patient CD34^-^ samples, the pooled scRNA-seq, CITE-seq and hashing libraries were processed with Cell Ranger (v6.1.1 and v6.1.2) using cellranger-multi pipeline (v1). The reads were aligned to the human genome GRCh38 with default parameters. The Seurat package (*126*) (v4.1.0) was used to perform unbiased clustering of the CD34^+^ sorted cells from each patient. In brief, for individual datasets, cells with UMI > or < 3 standard deviations from the mean UMI and mitochondrial gene percentage >10% were filtered (**fig. S1A, S4A**). The HTO data was normalized with centered log-ratio (CLR) transformation and used to assign the time-points for each experiment (*21*). The cells from each time-point were analytically separated into individual datasets based on the HTO counts (**fig. S1B**). These individual datasets (in case of CD34^+^ samples, together with the baseline ET samples from our previous work (*16*)), were integrated and underwent batch-correction within Seurat, which implements reciprocal principal component analysis (RPCA) and the principles of mutual nearest neighbor (*127*). Recommended settings were used for the integration (30 principal components for the anchor determining procedure in IntegrateLayers function). Principal component analysis was performed using variable genes using recommended settings (i.e., top 2000 variable genes using variance stabilizing transformation) (*127*). The first 30 statistically significant principal components were used as inputs to the UMAP algorithm for dimensional reduction and visualization (*128*). Clusters were manually assigned based on differentially expressed genes using the FindAllMarkers function using default settings (using top 2000 variable genes, in a minimum of 10% of cells in either of the two comparison sets as input, and log-transformed fold change of 0.25 as the threshold, using Wilcoxon rank sum test). The clusters were annotated according to canonical lineage markers identified previously in single-cell RNA-seq data of normal hematopoietic progenitor cells (*24*) into 16 main progenitor subsets based on expression of levels of these canonical markers (Fig. 1C, **fig. S2A-C, S4C**). For dimensional reduction and visualization of individual experiments (e.g., **fig. S1H**), top 3000 variable genes were included for principal component analysis, and the first 30 statistically significant principal components were used as inputs to the UMAP algorithm. The CITE-seq data after normalization using the CLR transformation was used to distinguish cell types (**Fig. 2B**, **fig. S2B**) and identify HSCs (**fig. S2D**). Same integration and downstream clustering methods were used to analyze the two hydroxyurea (HU) samples.

### IronThrone-GoT for processing targeted amplicon sequences and mutation calling

Analysis of the GoT library was carried out as described previously (*16*) using IronThrone pipeline V2.1 (*42*). Amplicon reads (Read 2) were screened for the presence of the primer sequence and the shared sequence (i.e., the expected sequence between the primer sequence and the mutation locus). Reads (Read 1) from GoT-IM experiments were also assessed for matching to the cell barcode list of the 10x dataset. A mismatch of 20% was allowed for all sequence matching steps. Only UMIs with at least 2 or more supporting reads were retained for final genotyping assignments, after the UMI collapse algorithm, as previously described (*42, 129*). Filtered cells were then genotyped as follows, as previously described: cells with at least one mutant UMI were categorized as mutant cells whereas cells with no mutant UMI and at least one wildtype UMI were identified as wildtype.

### Gene module scoring, differential expression and gene set enrichment analysis

For statistical analysis, when the variable in question was cell type identity (e.g., IGP vs IMP), cell type identity was entered as the fixed effect and samples as random effects in a linear mixed model. For IGP and IMP comparison, treatment status was also included as a fixed effect. When the variable in question was genotype status (e.g., *CALR* mutant versus wildtype), genotype status was entered as the fixed effect and samples as random effects in a linear mixed model. P-values were obtained from likelihood ratio tests of the full model with the effect in question against the model without the effect in question. Linear mixed effects analysis was performed using the lme4 package (v.1.2-1) (*130*).

Differential gene expression testing between two groups within an individual experiment (e.g., **Fig. 6J**) was performed using the logistic regression framework (*131*) with the FindMarkers function. The tested genes included the top 2,000 variable genes from the CCA integration, which were filtered for those expressed in at least 10% of either group. In aggregated differential gene expression analysis (e.g., treated versus baseline as in **Fig. 3B**), the two groups were compared via the linear mixed model framework, as previously described (*42*). Among CD34^+^ cells with GoT-IM data, IFN01-IFN10 samples at baseline and active treatment were included for the differential expression analysis. For each gene, the variable in question (i.e., treatment or mutation status) was entered as the fixed effect and samples as random effects. P-values were obtained from likelihood ratio tests of the full model with the effect in question against the model without the effect in question. Individual genes with abs(avg_log2FC)> 0.1 and adjusted p-value <0.05 were significant for further analysis such as module scoring for HSC-specific IFN⍺-induced upregulated or downregulated genes.

Pathway enrichment analysis was performed via a pre-ranked gene set enrichment approach (ranking based on the sign of the fold change * -log10(adjusted P-value)) using the msigdbr (v7.2.1) (*132*) and fgsea (v1.12.0) (*133*) R packages, using the canonical Hallmark pathway genes from MsigDB (*134*).

For examining gene module expression (e.g., HSC-specific IFN⍺-upregulated or downregulated gene signature), the function AddModuleScore within the Seurat package (*126*) was used to calculate the relative expression of the genes (that are significantly upregulated or downregulated, as described above) for each cell. To calculate the module expression of cell-cycle related genes, G2M phase and S phase marker genes were used as available in Seurat with CellCycleScoring function. Briefly, expression of a control gene module was calculated and subtracted from the average gene module expression of interest, as previously described (*44*). All analyzed genes were classified based on average expression into 24 bins, and for each gene in the module, 100 control genes are randomly selected from the same expression bin as the gene of interest (*44*). For the overall IGP module score (**Fig. 2C**), the cells were first scored for the upregulated and downregulated genes (**Fig. 1E**); the downregulated gene score was subtracted from the upregulated gene score to obtain the overall IGP score.

### RNA velocity analysis using scVelo and partition-based graph abstraction

RNA velocity was assessed from the spliced and unspliced transcript variants using scVelo (v0.2.5) (*34*). Counts from loom files generated with Velocyto (v0.17.17) (*35*) for each GoT-IM sample (IFN01-IFN09) were normalized and filtered (UMI counts > 100 and UMI count < 20000). The annotated data matrices were combined by using anndata.concat() command and cell-specific annotations such as cell-type, time-point, UMAP and PCA embeddings were imported from the GoT-IM integrated object. Connectivities between the cell clusters were quantified using scvelo.tl.paga() within the partition-based graph abstraction (PAGA) framework (*36*).

### GoT-ATAC

Cryopreserved bone marrow mononuclear cells were thawed and stained using standard procedures with the surface antibody CD34-PE (clone AC136, dilution 1:50, MACS) and DAPI (Sigma-Aldrich), according to manufacturer’s protocol. Cells were subsequently sorted for DAPI^−^, CD34^+^ cells using BD Influx at the Weill Cornell Medicine flow cytometry core. Nuclei were isolated from DAPI^-^, CD34^+^ cells according to 10x Genomics Demonstrated Low Cell Input Nuclei Isolation protocol. Lysis buffer was prepared following manufacturer’s recommendations and then split into aliquots for each serial sample. Either TotalSeq-A Anti-Nuclei Pore Complex Proteins Hashtag 9 or 10 Antibody (1μL at 1:5 dilution; BioLegend) was added to each aliquot of lysis buffer. Low-input nuclei isolation was otherwise performed following manufacturer’s recommendations. Subsequently, nuclei from each time-point were counted and pooled together at approximately equal proportions. For IFN07, additional nuclei from the IFN⍺-treated sample were available to be run on a separate lane. Single-nucleus gene expression (GEX) and chromatin accessibility libraries were constructed from the pooled nuclei according to the Chromium Next GEM Single Cell Multiome User Guide (10x Genomics).

Genotyping libraries targeting the *CALR* mutant transcripts were constructed from the remaining amplified cDNA, similar to the original GoT method. For each PCR, 12.5μL Kapa HiFI HotStart Ready Mix was mixed with 0.75μL of 10uM forward primer, 0.75μL of 10uM reverse primer, 3μL cDNA and nuclease-free water for a total reaction volume of 25μL. In the first PCR, 3μL cDNA was re-amplified with Partial TSO and Partial Read 1 primers (**table S31**), using the following PCR condition: 98°C for 3min; 3 cycles of 98°C for 15sec, 67°C for 20sec and 72°C for 1min; 72°C for 1min. The re-amplified sample was purified and concentrated via 0.7X SPRI cleanup, eluting it into 10μL Buffer EB. To pre-enrich the *CALR* mutation locus, a gene-specific PCR was performed with 3μL of cleaned re-amplified cDNA and Partial Read 1 and gene-specific primers (**table S31**). The following PCR condition was used: 98°C for 3min; 11 cycles of 98°C for 20sec, 60°C for 20sec, and 72°C for 2min; 72°C for 2min. After 0.7X SPRI cleanup, cDNA was eluted into 10μL Buffer EB. *CALR* locus-specific amplification was then performed with 3μL of cleaned gene-specific amplified cDNA and SI-PCR and loci-specific Primers, using the PCR condition: 98°C for 3min; 11 cycles of 98°C for 20sec, 60°C for 20sec, and 72°C for 2min; 72°C for 2min. A 0.7X SPRI cleanup was performed, and cDNA was eluted into 11μL Buffer EB. Finally, to construct the targeted amplicon library, loci-amplified cDNA was mixed with P5 Generic and RPIx indexing primers (**table S31**) and amplified with the PCR condition: 98°C for 3min; 5 cycles of 98°C for 15sec, 60°C for 20sec, and 72°C for 1min; 72°C for 1min. The constructed library was cleaned via 0.8X SPRI cleanup and eluted into 12μL Buffer EB.

At the cDNA amplification stage of the Chromium Next GEM Single Cell Multiome protocol, supernatant from the 0.6X size selection was retained and was used to generate the hashing libraries as per HTO protocol (*21*) with the following modification. For the hashing library construction step, the PCR reaction was prepared with 0.65μL of 10uM SI-PCR primer, 0.65μL of 10uM TruSeq DNA D7xx_s primer (**table S31**), 11.25μL cleaned supernatant and 12.5 μL KAPA HiFi HotStart Ready Mix (Roche, Basel, Switzerland). Hashing, gene expression and genotyping libraries were pooled and sequenced together on a NovaSeq (Illumina) with cycle settings: 28 cycles for Read 1, 90 cycles for Read 2, 10 cycles for i7 sample index and 10 cycles for i5 sample index. The ATAC library was sequenced separately on a NovaSeq, with cycle settings: 50 cycles for Read 1 and Read 2, 8 cycles for i7 sample index and 24 cycles for i5 sample index.

### Data preprocessing, alignment and cell type identification for GoT-ATAC

For single-nuclei GoT-ATAC data from IFN01, IFN03, IFN07 and IFN13 patient CD34^+^ samples, 10x data were processed using Cell Ranger (v6.1.1 and v6.1.2). Multi-omic nuclear data for snATAC-seq and snRNA-seq were processed together with Cell Ranger Arc (v2.0.0). snRNA-seq data was also combined with cell hashing data (HTO) and run using the Cell Ranger Multi pipeline (v1). The reads were aligned to the human reference genome GRCh38. The downstream analysis of the processed data was performed using Seurat (v4.1.0) (*126*) and Signac (v1.5.0) (*135*) packages. For the ATAC analysis, we called peaks on individual samples using MACS2 peak caller (*136*). Gene annotations from EnsDb.Hsapiens.v86 and motifs annotations from Cis-BP (*137*) for TF binding motifs were utilized. Cells with blacklist ratio >0.02, TSS enrichment < 2 and nucleosome signal >4 were filtered out. For the RNA data, cells with UMI > or < 3 standard deviations from the mean UMI or mitochondrial gene percentage > 25%, were filtered (**fig. S7A-D**). The nuclei hashing data processing was performed as for the GoT-IM data (**fig. S7E**). As nuclear hashing is known to be noisier than cell hashing (*135*), the nuclear hashing data was used in combination with cell clustering data, as the cells cluster based on treatment status. After the cells were segregated analytically based on time-point, the datasets were integrated as described for the GoT-IM data. snRNA-seq data were integrated where they underwent batch correction with canonical correlation analysis (CCA) within Seurat and the principles of nearest neighbor (*127*). Recommended settings were used for the integration (30 canonical correlation vectors for canonical correlation analysis in the FindIntegrationAnchors function and 30 principal components for the anchor weighting procedure in IntegrateData function). Principal component analysis and cell type assignments were performed as described in GoT-IM. Integration via the ATAC-seq data was performed by normalizing the merged counts using first term frequency inverse document frequency (TFIDF) normalization with RunTFDIF followed by linear dimensional reduction using latent semantic indexing (LSI). The first 2:30 dimensions were retained, and batch-correction was performed with runHarmony (Harmony, v0.1.0) which iteratively learns cell-specific linear correction function to account for batch effect. Pseudotime analysis was performed by constructing cell lineage trajectory with Monocle 3 (*138–140*). We used as.cell_data_set function from SeuratWrappers package to convert Seurat object to CellDataSet object used by Monocle 3. IronThrone-GoT protocol was used to determine mutation calling within the GoT-ATAC assay, as described for GoT-IM. A requirement of at least four reads supporting a UMI was implemented for GoT-ATAC. To capture genotyping reads within the snRNA-seq data, we called *CALR* variants using the gene expression BAM derived from the standard Multiome workflow. BAMQL (*141*) and the CIGAR string of the BAM file were used to parse reads containing the expected wildtype and mutant sequences. Cells expressing at least one mutant UMI were categorized as mutant and cells with just wildtype UMI were assigned as wildtype. Genotyping assignments were then appended to calls made by IronThrone.

### Identification of distal regulatory elements with gene-peak cis-association

For each GoT-ATAC sample, we examined all ATAC peaks within ± 500kb of all annotated TSS to identify regulatory networks of genes using LinkPeaks function (*72*). Pearson correlation between gene expression and accessibility of the peaks located in the window was calculated after correcting for bias arising from GC content, overall accessibility and peak size. Recommended settings were implemented (200 background peaks per peak with similar GC content and accessibility, P-value < 0.05 and min.cell = 10).

### Motif enrichment analysis in GoT-ATAC

Per-cell TF motif activity score (chromatin accessibility) was calculated by running chromVAR (v1.18.0) (*69*). We used the curated Cis-BP motif database (*137*) which contains 1141 human TF motif position frequency matrices (PFMs). The function matchMotifs was first called to identify which peaks contain which motifs (P-value = 5 x 10^-5^). A set of background peaks that are similar to a peak in GC content and average accessibility was internally picked and used for normalizing the deviation scores. Deviation Z-scores, namely bias-corrected deviation z-scores in accessibility from the expected accessibility based on the average of all the cells, were then calculated for each TF motif and each cell.

To perform differential motif enrichment analysis, within a sample and on the deviation z-score computed by chromVAR, we applied the function FindMarkers in Signac (Wilcoxon Rank Sum test), where the average difference in z-score between the groups was calculated. For integrated data, we combined the P-values (Fisher’s method) and calculated weighted mean deviation score across individual samples. P-values were adjusted by the Benjamini-Hochberg method.

To find over-enriched motifs for a group of genomic features, the FindMotifs function was used, accounting for accessibility and GC content bias by selecting 5000 accessible background peaks with similar GC content for each feature set. To identify motifs enriched in a singular genomic range, FIMO (v5.4.1) from MEME suite (*142*) was used to scan for Cis-BP TF motifs along the nucleotide sequence from human reference genome GRCh38 with a p-value threshold of 0.0001. We de-prioritized zinc fingers (ZNFs) in the list of motifs specific to genomic regions as each individual ZNF binds to three bp motif leading to more frequent matches and higher match scores (*143*). chromVAR was used to compute the synergy between pairs of TF motifs, where synergy is defined as the excess variability of chromatin accessibility for peaks sharing both motifs compared to a random subsample of the same size of peaks with one motif. High synergy usually indicated a cooperative binding relationship between pairs of TFs. The function getAnnotationSynergy was called to calculate synergy scores (*69*).

### Retrospective flow cytometry data analysis

Retrospective flow cytometry data analysis was performed in accordance with relevant guidelines, regulations and approval by the Institutional Review Board at Weill Cornell Medicine (IRB #1007011151). Patient flow cytometry data selected corresponded to patients with myeloproliferative neoplasms and which had the same antibody panel analyzed by flow cytometry. Patients with a diagnosis of ET or polycythemia vera with no increase in myelofibrosis or blast counts were included in the study. The antibody panel chosen for evaluation was a modified version of the EuroFlow AML/MDS tube #4 (*144, 145*) that consisted of the following antibody-fluorophore pairs, in addition to forward scattering and side scattering pulse area and width measurements (FSC-A, FSC-H, SSC-A, and SSC-H): cytoplasmic TdT/FITC (clone HT-6, Agilent/Dako, cat. F7139), CD56/PE (clone C5.9, Cytognos, cat. CYT-56PE), CD34/PerCP-Cy5.5 (clone 8G12, BD Biosciences, cat. 347213), CD117/PE-Cy7 (clone 104D2D1, Beckman Coulter/Immunotech, cat. IM3698), CD7/APC (clone GP40 [Leu-9], Invitrogen, cat. 17-0079-42), Fixable Viability Stain 700 (BD Biosciences, cat. 564997), CD19/APC-H7, HLA-DR/Pacific Blue (clone SJ25C1, BD Biosciences, cat. 643078), and CD45/V500 (clone 2D1, BD Biosciences, cat. 347213). Data was collected using BD LSR II flow cytometers, with approximately 500,000 events collected per antibody panel per sample to generate the raw data FCS files.

Custom software was developed using python, FlowKit (*146*) and umap-learn (*128*) to detect the antibody panels that were used to generate each raw data FCS file and to determine the unused flow cytometer channels that should be disregarded using the self-contained metadata for each file. Subsamples from each FCS file were combined to make an “ensemble FCS file” that could be used to create the UMAP embedding that could be applied to each of the individual files. Each subsample consisted of the same number of randomly selected flow cytometer events such that the combined total number of events was approximately 250,000 for each unique antibody panel. The various channels were normalized and processed using UMAP to calculate the normalization constants and UMAP embedding that were then applied to all FCS files of the given antibody panel. Of note, the UMAP calculations included the forward scatter height (FSC-H), side scatter height (SSC-H) and each of the defined fluorescence channels. The normalization factors and UMAP embedding were then applied to all the individual files. Modified FCS files were created that included the UMAPs as additional channels for subsequent evaluation and gating using FlowJo software (v10.8.1, FlowJo LLC, Ashland, Oregon, USA).

Using FlowJo, appropriate gates based on the UMAP plot were determined using the ensemble FCS file. Additional standard gating was also performed using the original data channels (gating using the other channels is essential to determine the identities of the various cell clusters within the UMAP plots). UMAP gates were based on data after gating out doublets and non-viable cells via standard gating approaches. Once the UMAP gates were determined to adequate satisfaction (sufficient segregation of cell subpopulations and verified to encompass cells of the same or similar type), they were then applied to all the FCS files of the given antibody panel. The FCS files were divided into untreated/only aspirin treated cohort (n = 33), an interferon-treated cohort (n = 9), a hydroxyurea-treated cohort (n = 10). The ratios and absolute numbers of cells, as well as other summary statistics were then calculated, and the values exported as CSV files. Statistics included numbers of CD34^+^ blasts; CD19^+^, cTDT^+^, CD34^+^ lymphoid progenitors; CD19^+^ lymphocytes; CD19-negative lymphocytes/NK cells; and monocytes. Relevant distributions of cells for each cohort are plotted, and the Wilcoxon p-statistic was calculated for various compared distributions.

### Multiplexed Immunofluorescence

Multiplexed immunofluorescence (mIF) was performed using the Opal system (Akoya Biosciences) by staining 4 micron-thick Bouin-fixed, paraffin-embedded whole-tissue sections from decalcified human bone marrow core biopsy specimens in a Bond RX automated tissue stainer (Leica Biosystems, Buffalo Grove, IL), as described previously (*147, 148*). Briefly, tissue sections were first deparaffinized prior to EDTA-based antigen retrieval (Leica ER2 solution, 20min). A cyclical staining protocol was then performed, with horseradish peroxidase-mediated deposition of tyramide-Opal fluorophore constructs (Akoya Biosciences) in each cycle, with intervening application of heat, citrate-based epitope retrieval solution (Leica ER1), and Bond Wash Solution (Leica) to execute stripping of primary/secondary antibody complexes between staining cycles. Finally, 4’, 6-diamidino-2-phenylindole (Spectral DAPI, Akoya Biosciences) was applied per provided protocols to label nuclei. The following panel of primary antibody/fluorophore pairs was applied to all cases, in a sequential order as shown: 1) Opal 480/anti-mutant CALR (1:120, CAL2, Dianova), 2) Opal 520/anti-CD38 (1:50, 38C03 (SPC32), Invitrogen), 3) Opal 570/anti-CD117 (1:100, D3W6Y, Cell Signaling), 4) Opal 620/anti-TdT (1:8, SEN28, Invitrogen), 5) Opal 690/anti-CD34 (1:100, QBEND/10, Invitrogen), 6) Opal 780/anti-Ki67 (Ready-to-use, MM1, Leica). Slides were cover-slipped using ProLong™ Diamond Antifade Mountant (Invitrogen). Whole slide scans were subsequently obtained at 20X magnification using the Vectra Polaris Automated Quantitative Pathology Imaging System (Akoya Biosciences) to generate a collection of tiled images, which were subsequently spectrally unmixed in InForm (v2.4.8, Akoya Biosciences). Unmixed tiles were finally fused together in HALO (v3.3.2541.231, Indica Labs) to generate a multi-layered TIFF image file for each sample, which was used in downstream analyses.

### Image Analysis with PathML

Vectra whole-slide images (WSIs) were digitized using digital whole-slide scanners and stored in tiff format. WSIs of bone marrow sections were captured (n = 9 samples). Each sample was stained based on 8 cell markers including DAPI, mutant-specific CALR, CD38, CD117, TdT, CD34, Ki67, and auto-fluorescence. The image contrast was enhanced using histogram equalization in Fiji. To analyze the Fiji-preprocessed WSIs, PathML, a toolkit for computational image analysis (*149*), was applied to images. Images were loaded and divided into equal-sized tiles on which we ran our preprocessing pipeline. This pipeline starts with coercing the tile shape into the standard x,y,c format followed by segmentation. Nuclear and cellular segmentation were performed using the Mesmer (*150*) deep learning segmentation model implemented in PathML with DAPI and autofluorescence used as nuclear and cell membrane markers, respectively. Subsequently, we used the ‘QuantifyMIF’ function from PathML to convert the segmented images into count matrix which includes the intensity of each marker in each segmented cell along with cell coordinates, size, and eccentricity. To remove the noise from the count matrix data, any cell with raw intensity less than 50 for DAPI and auto-fluorescence markers was excluded from the analysis. Next, the thresholds were obtained to find the positive and negative expression level for each cell marker. The thresholds were manually set for each marker based on examination by a board-certified hematopathologist.

### CUT&RUN Assay

UT7 cell lines expressing MPL (thrombopoietin receptor) and either the mutant *CALR* (type 1, L367Tfs*46) or wildtype *CALR* transgene (*93*) were seeded in 10cm plates in DMEM supplemented with 10% FBS (Thomas Scientific) and 5 ng/ml GM-CSF (Miltenyi-Biotec). Cells were treated with 0.1μg/ml Recombinant Human IFN-alpha2a (RC217-14) or PBS-1X control for 24hrs. After treatment, cells were washed 1x with PBS and collected for use with the CUTANA™ CUT&RUN assay (EpiCypher) according to the manufacturer’s protocol. 0.5μg of anti-human rabbit monoclonal PU.1 antibody (ab76543) and IgG antibody (ab172730) were used in the assay.

CUT&RUN data were down-sampled to have same number of reads across all conditions within the sample replicate to account for difference in sequencing depth. Among the down-sampled reads, low-quality reads were filtered out using Trimmomatic (*151*), resulting reads were aligned with hg38 genome with Bowtie2 (*152*), and PCR duplicates were removed with SAMtools (*153*) with a previously described workflow (*154*). Aligned reads were then used for peak calling with MACS3 (*136*) with a q-value threshold of 0.01 and reads from IgG antibody as the control. SPI1 over-enrichment in the called peaks was confirmed with Simple Enrichment Analysis (SEA, v5.5.5) (*155*). Overlapping peaks between replicates of the same condition were kept for further processing. The peaks were processed with multiBamSummary function from deepTools (*156*) to obtain counts matrix per sample for downstream analysis.

Differential peak enrichment analysis was run between conditions (MUT vs WT and IFN⍺-treated vs control) using DESEQ2 (*157*) for distal PU.1 peaks. Differential peaks in each condition were centered around SPI1 motif and motif enrichment was performed for regions +/- 250 bp from SPI1 motif using HOMER (v.4.1.1) (*96*). Differential ranking between each motif two conditions was calculated (**Fig. 6H**, **right, fig. S12B, D**).

### Single-cell Differentiation Assay

One day before sorting HSCs, flat-bottom 96-well plates were coated with 60μL 0.2% gelatine (Sigma) per well for 1 hour and then removed. Low passage murine MS5 stroma cells were plated at a density of 1500 cells/well in 100μL Myelocult H5100 medium (Stem Cell Technologies), 1% Penicillin-Streptomycin (Pen/Strep, 10,000 U/mL, Gibco) and 1% glutamine (Gibco). On the day of the assay, the medium was changed to lympho-myeloid media consisting on 100μL/well Myelocult H5100 medium, 1% Pen/Strep and 1% glutamine supplemented with the following cytokines: IL-2 10ng/mL, IL-6 20ng/mL, IL-7 20ng/mL, SCF 100ng/mL, TPO 50ng/mL, G-CSF 20ng/mL, FLT3L 10ng/mL and GM-CSF 20ng/mL (all from Bio-Techne). Cryopreserved bone marrow mononuclear cells (n = 2 baseline and 3 IFN⍺-treated samples) were thawed and CD34^+^ cells were isolated using the EasySep™ Human CD34 Positive Selection Kit II (StemCell Technologies #17856) following manufacturer’s protocol. CD34^+^ isolated cells were then stained with FITC CD45RA (1:50), APC-Cy7 CD34 (1:200), APC CD90 (1:50), Pe-Cy7 FLT3 (1:100), PerCP-Cy5.5 HLA-DR (1:100), and BV786 CD41 (1:100) (antibody details in **table S31**). CD34^+^ CD90^+^ cells were index-sorted by FACS directly into 96-well plates with pre-plated MS5 stromal cells. Eight days after sorting, 70μL of media were removed from the top of the plate without disturbing the colonies, and 170μL of IMDM medium (Gibco) with 10% BIT 9500 (StemCell Technologies), 1% Pen/Strep, 1% glutamine, supplemented with IL-2 10ng/mL, IL-3 20ng/mL, IL-6 10ng/mL, IL-7 10ng/mL, SCF 100ng/mL, TPO 50ng/mL, G-CSF 10ng/mL, FLT3L 10ng/mL, SDF1 5ng/mL and 2-ME (1.8μL per 50mL), were added. Colonies were analyzed under the microscope 16-18 days after sorting and all visible colonies were detached by pipetting from the stromal cell layer and transferred into a 96-well U-bottom plate using a plate filter (Pall AcroPrep), to prevent the carryover of MS5 cells. Cells were stained for 1 hour at 4°C with FITC CD66b (1:100), PE CD16 (1:100), PerCP-Cy5.5 CD184 (1:100), Pe-Cy7 CD83 (1:100), APC CD14 (1:100), V500 CD45 (1:100), BV650 CD71 (1:100), and BV711 CD33 (1:100) (antibody details in **table S31**) with 50μL/well antibody mix and washed afterwards with 200μL PBS and 2.5% FBS. Immunophenotype of the colonies was assessed in a BD LSRFortessa Analyzer. Flow cytometric analysis was performed using FlowJo and OMIQ. To generate differentiation plots, OMIQ was used to gate live cells and calculate UMAP dimensional reduction based on expression of CD14, CD34, CD45, CD71, CD33, CD16, and CD66b after integrating cells across every colony using recommended settings. A total of 177 colonies were included in the analysis from 5 samples (after filtering for colonies with <350 live cells). GEMM colonies showed CD71^+^, CD14^+^, and CD66b^+^ populations. GM colonies showed both CD14^+^ and CD66^+^ populations (at least 1% each). Neutrophil-only colonies showed <1% CD14^+^ cells and at least 95% CD66b^+^ cells. To increase the stringency of the neutrophil-only assignment, a minimum of 4000 events was required for this assignment. The remainder of the colonies were labeled as early myeloid (EM) colonies showing a large CD33^+^ population (>40%) that lacked either CD14 or CD66b expression.

### Western Blot

1x10^6 K562 cells were seeded in triplicate in a 10cm plate in RPMI 10% FBS 1% Pen/Strep. Recombinant Human IFN-alpha2a (RC217-14) was diluted in PBS and added to treatment condition plates at a concentration of 2000U/ml. Corresponding control plates were treated with equal volumes of PBS. Cells were kept in media with or without IFN⍺ for 24hrs or 48hrs, after which cells were collected and cell count and viability was recorded. Cells were then centrifuged at 300g for 5min at 4°C. Pellet was resuspended in cold PBS and centrifuged at 300g for 5min at 4°C. Dry cell pellets were frozen at -80°C until ready for use. Cell pellets were lysed in RIPA Buffer (ThermoFisher #89901) with Protease Inhibitor (ThermoFisher #78420) and Phosphatase Inhibitor (ThermoFisher #78429) for 15min on a shaker at 4°C. Total protein was quantified using colorimetric BioRad DC Protein Assay (#5000113). Samples were run on a Novex Tris-Glycine Mini Protein Gel (ThermoFisher) according to manufacturer’s protocol and transferred via standard wet transfer protocol. After blocking with 5% milk for 1hr at room temperature, blot was stained with anti-human mouse PU.1 monoclonal antibody (1:1000) (CST #89136) and anti-human rabbit Vinculin monoclonal antibody (1:10000) (ab129002) diluted in Intercept T20 Antibody Diluent (LI-COR) overnight at 4°C. After washing with TBS-T (1X TBS, 0.1% Tween 20) buffer, blot was stained with secondary antibodies including anti-mouse IRDye 800CW Goat anti-Mouse IgG Secondary Antibody (1:5000) AND IRDye 680RD Goat anti-Rabbit IgG Secondary Antibody (1:5000) for 1hr at room temperature (antibody details in **table S31**). Blot was then imaged using LI-COR machine and absorbance quantified using LI-COR software.

### Lentiviral Constructs for Overexpression Experiments

Lentiviral overexpression (OE) vectors and their corresponding control vectors were designed and obtained from VectorBuilder. All constructs included a lentivirus backbone (pLV) with gene-specific inserts, a fluorophore and an antibiotic resistance insert. The RFX3-OE insert consisted of human RFX3 ORF (NM_001377999.1) driven by the human phosphoglycerate kinase (hPGK) high-expression promoter, with CMV-mCherry-T2A-Puro for selection. The SPI1-OE and CEBPA-OE inserts consisted of the human SPI1 ORF (NM_001080547.2) and human CEBPA ORF (NM_001287424.2) respectively, driven by the hPGK promoter, with CMV-EGFP-T2A-Puro for selection. The control vectors contained the hPGK promoter driving an empty ORF stuffer, followed by CMV-mCherry-T2A-Puro or CMV-EGFP-T2A-Puro. The plasmid vectors were generated and amplified by VectorBuilder.

### Lentivirus Production

HEK-293T cells were seeded at a density of 2x10^6^ cells in DMEM, 10% FBS and 1% Pen/Strep in a 10cm plate. After 24 hours, media was changed to DMEM and 10% FBS. 9μg OE or control plasmid was added to 1mL Opti-MEM (ThermoFisher) containing 3μg pMD2.G plasmid and 8μg psPAX2 plasmid and incubated with Lipofectamine 2000 (ThermoFisher) in 1mL of Opti-MEM for 15min at room temperature. Mixture was then added dropwise to the cells and incubated at 37°C for 24 hours. Media containing virus was collected over the next 48 hours. Lentivirus was then concentrated using Lenti-X Concentrator (Clontech #631231) according to manufacturer’s protocol. Viral titer was determined using the qPCR Lentivirus Titer Kit (Applied Biological Materials LV900).

### Lentiviral RFX3 Overexpression of CD34^+^ Cells

Frozen umbilical cord blood mononuclear cells were purchased from the NYC Blood Center. After cells were thawed rapidly at 37°C, red blood cell lysis was performed using ACK Lysis Buffer (ThermoFisher) on ice for 7min. Cells were then centrifuged at 300g for 5min at 4°C. Supernatant was discarded, and cell pellet was resuspended in cold MACS buffer (PBS-1X with 0.5% BSA). Cells were centrifuged at 300g for 5min at 4°C, and resuspended in cold MACs buffer at a concentration of 1x10^7 cells/mL. CD34^+^ cells were then isolated with the EasySep™ Human CD34 Positive Selection Kit II (StemCell Technologies #17856) according to manufacturer’s protocol. After isolation, cells were counted and plated in 96-well round-bottom plates (Falcon) in StemSpan™ SFEM II media (StemCell Technologies) with StemSpan™ CD34^+^ Expansion Supplement (StemCell Technologies) and 1% Pen/Strep. After 24 hours in culture, cells were spun down and resuspended in minimal media consisting of StemSpan StemSpan™ SFEM II media with StemSpan™ CD34^+^ Expansion Supplement, 1% Pen/Strep, 10μM prostaglandin E2 (StemCell Technologies #72192) and 100ng/μl poloxamer 407 (Millipore Sigma P2164030). Cells were split into 2 technical replicates each for untransduced condition, mCherry-control vector transduction, and RFX3-OE vector transduction. Lentivirus or PBS control was added at an MOI of 100 directly into the well. Cells were then spinoculated at 300g, 32°C for 1 hour. After the spin, cells were incubated in same lentivirus-containing media for 24 hours at 37°C before resuspension in fresh media consisting of StemSpan™ SFEM II media with StemSpan™ CD34^+^ Expansion Supplement and 1% Pen/Strep. After 48 hours of incubation, cells were stained with CD34-APC antibody (1:50) (**table S31**). Cells were then FACS isolated for the CD34^+^ population in the untransduced condition and the CD34^+^ mCherry^+^ populations in the RFX3-OE and mCherry-control conditions. Three independent transduction experiments were completed with new units of umbilical cord blood cells.

Untransduced, mCherry-Control, and RFX3-OE CD34^+^ cells were stained with Cell Hashing antibodies (HTO TotalSeq-A, BioLegend) for 30 min at 4°C. Cells were then washed with FACs Buffer three times, pooled, and counted for loading. scRNA-sequencing was then performed using the 10x 3’ v3.1 platform according to manufacturer’s protocol. HTO demultiplexing and downstream analyses such as differential gene expression and gene set enrichment were performed as in GoT-IM for CD34^+^ bone marrow samples. scRNA-seq experiment was completed on one replicate of three independent transduction experiments.

The FACS-sorted cells were also used for colony-forming unit (CFU) assays. MethoCult total media were prepared using MethoCult™ H4034 Optimum (StemCell Technologies # 04044) supplemented with 10ng/ml human FLT3L and IL-6 (Miltenyi-Biotec) and 1% Pen/Strep. 250 cells per group (Untransduced, mCherry-Control, and RFX3-OE) were added to 3mL MethoCult total and vortexed thoroughly. Tubes were then left at room temperature until bubbles rose to the surface. 1.25mL of MethoCult with cell suspension was transferred per well of a 6-well plate (2 replicates per group) via blunt-edge 16g needle. Plate was incubated at 37°C for 14 days after which colonies were counted and identified by morphology. CFU assay was completed on all three of the independent transduction experiments.

### Lentiviral Transduction of K562 Cells

K562 cells were seeded at density of 3x10^5^ cells/mL in 24-well tissue-culture treated plates (Falcon) in 500μL of K562 media (RPMI with 10% FBS and 1% Pen/Strep) in triplicate for each of the following conditions: untransduced, mCherry-Control, RFX3-OE, EGFP-control, SPI1-OE, and CEBPA-OE. Concentrated lentivirus (or PBS-1X control for untransduced condition) was added directly to each well at MOI of 20, shaken gently and incubated at 37°C for 24 hours. After 24 hours in lentivirus-containing media, plates were spun at 300g for 5min at room temperature, and supernatant media was replaced with fresh K562 media. Cells were then incubated at 37°C for 72 hours to allow for vector expression. Post-72 hours, cells were pooled per condition and washed with FACS buffer (PBS-1X with 2% FBS). mCherry+ or EGFP+ cells per condition were then FACS-isolated and plated in K562 media in tissue-culture treated T25 flasks and allowed to recover for 48 hours before use in subsequent experiments. Two independent transduction experiment replicates of mCherry-Control, RFX3-OE and SPI1-OE and three independent replicates of EGFP-control and CEBPA-OE were performed.

### *In vitro* IFN⍺ Treatments of SPI1-OE and CEBPA-OE Cells

5x10^6^ untransduced, EGFP-control and SPI-OE K562 cells were seeded in 24-well TC-treated plates in 1mL of K562 media (RPMI with 10% FBS and 1% Pen/Strep). Recombinant Human IFN-alpha2a (RC217-14) was diluted in PBS-1X and added to treatment condition plates at a concentration of 2000U/mL. Equal volume of PBS-1X was added to untreated controls. After gentle shaking, cells were incubated at 37°C for 24 hours. Post-incubation, cell suspension was collected and spun at 350g for 5min at 4°C. Supernatant was aspirated and cell pellet was washed with 1mL cold PBS. Cells were spun again at 350g for 5min at 4°C. Supernatant was aspirated and dry cell pellets were frozen at -80°C until RNA was extracted.

### RNA extraction and RT-QPCR

Total RNA was prepared using the RNeasy Plus Micro or RNeasy Micro kit (Qiagen, #74034/ 74004) according to the manufacturer’s instruction. RT-QPCR assays were performed using a QuantStudio™ 5 Real-Time PCR System (Applied Biosystems). One-step qRT-PCR assays were performed using the Power SYBR™ Green RNA-to-CT™ 1-Step Kit (Thermo Fisher Scientific, #4389986) according to manufacturer’s instructions. The thermal cycling conditions comprised a reverse transcription step at 48°C for 30min, an initial denaturation (enzyme activation) step at 95°C for 10min and 40 cycles at 95°C for 15s and 60°C for 1min. Transcripts of the TBP gene encoding the TATA box-binding protein (a component of the DNA-binding protein complex TFIID) were quantified as an endogenous RNA control. Quantitative values were obtained from the cycle number (Ct value), according to the manufacturer’s manuals (Applied Biosystems) and 2^-ΔCT^ values were calculated relative to TBP. Sequences of primers used for QPCR are listed in **table S31**.

### Statistics and reproducibility

Linear mixed modeling (LMM) was implemented using the lme4 R package (v.1.2-1). In all cases, LMMs were generated with/without cell mutational status, treatment status or cell types, as specified in the figure legends. This allowed inclusion of random effects to account for biological variation. We included patient sample as random effects in our statistical comparisons. P-values were calculated by analysis of variance with likelihood ratio test using the Stats R package (v.3.5.1) between two models (with or without the fixed variable of interest). P-value adjustments were done with Benjamini-Hochberg FDR-correction unless specified otherwise.

For all box plots presented, the box represents the interquartile range; upper and lower whiskers represent the largest and smallest values within 1.5 times the interquartile range above the 75th or below the 25th percentile, respectively; the central line represents the median. Dots represent outlier values or data value distributions. For all violin plots, the violin represents the kernel probability density of the data and dots represent the observed values.

---

**fig. S1 (related to Fig. 1).**
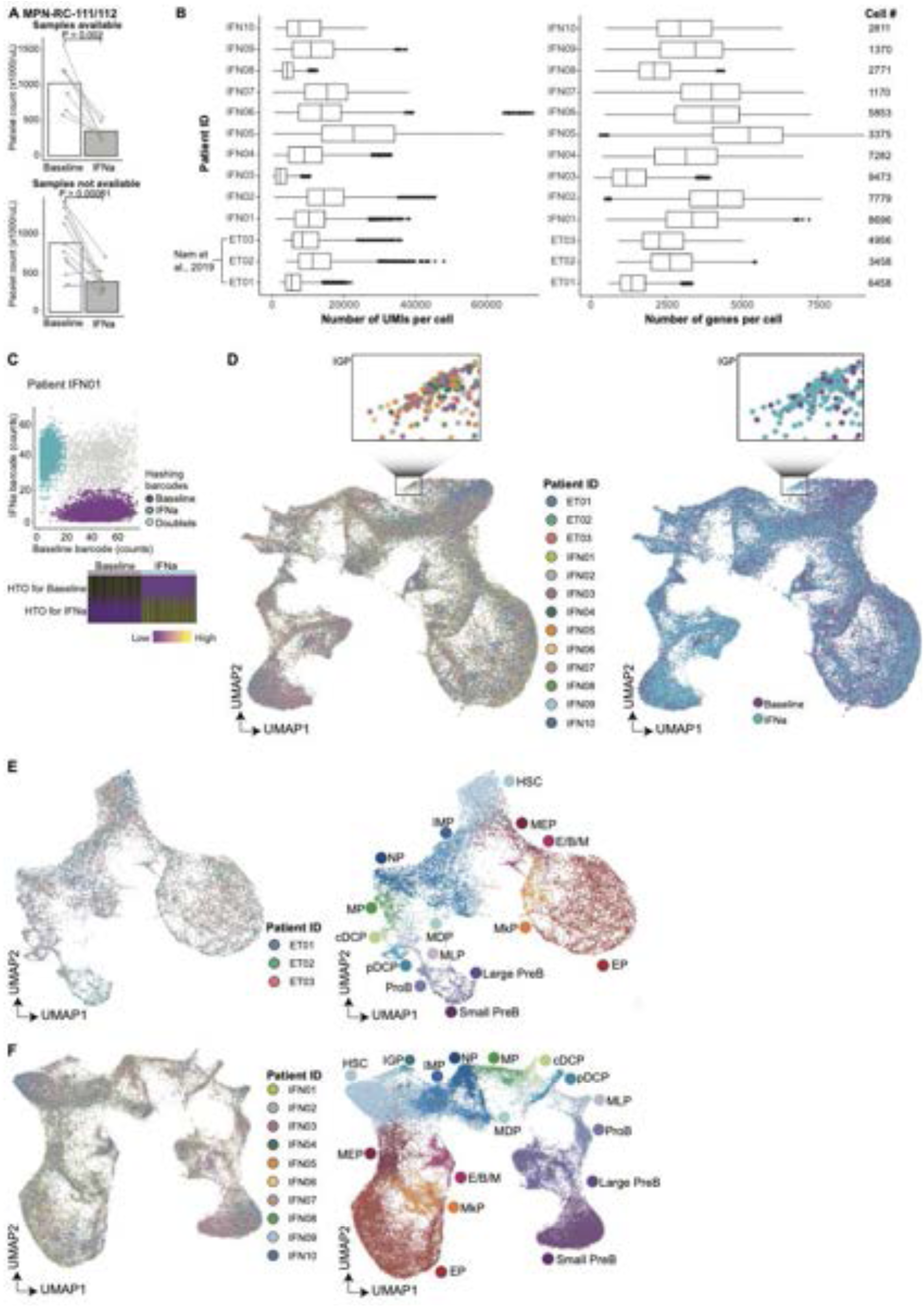
Genotyping of Transcriptomes integrated with immunophenotyping (GoT-IM) profiles thousands of CD34^+^ cells at baseline and following treatment with IFN⍺. **A.** Change in platelet frequency at baseline and one year upon IFN⍺ treatment for samples (IFN01-IFN13) included in the study (top) and samples that were not available (bottom) from MPN-RC-111/112 trial. Patient samples with both baseline and year 1 were included. P-values from likelihood ratio test of LMM with/without treatment status (methods). **B.** Box plots showing number of UMIs (left panel), and genes (right panel) detected per cell in sorted CD34^+^ hematopoietic progenitors from each patient after filtering based on quality control (QC) metrics (**methods**). **C.** Top: Time-point assignment (data demultiplexing) using time-point specifying barcodes. Cells in which both barcodes are detected are considered as doublets and excluded. Representative patient sample IFN01 (n = 8,696 cells) shown. Bottom: Heatmap showing HTO expression level for baseline and IFN⍺-treated cells from IFN01. **D.** Uniform Manifold Approximation and Projection (UMAP) of sorted CD34^+^ progenitors (n = 65,452 cells, samples from this study and Nam et al., 2019) highlighted by patient ID (n = 13 individuals, left) and treatment status (right) after integration with zoomed in view of the region with IGP cluster. **E.** Integrated UMAP of sorted CD34^+^ progenitors from Nam et al., 2019 (n = 14,872 cells) highlighted by patient ID (n = 3 individuals, left) and cell type clusters (right). **F.** Integrated UMAP of sorted CD34^+^ progenitors of ET patients from the MPN-RC-111/112 trials (n = 50,580 cells) highlighted by patient ID (n = 10 individuals, left) and cell type clusters (right).

**fig. S2 (related to Fig. 1).**
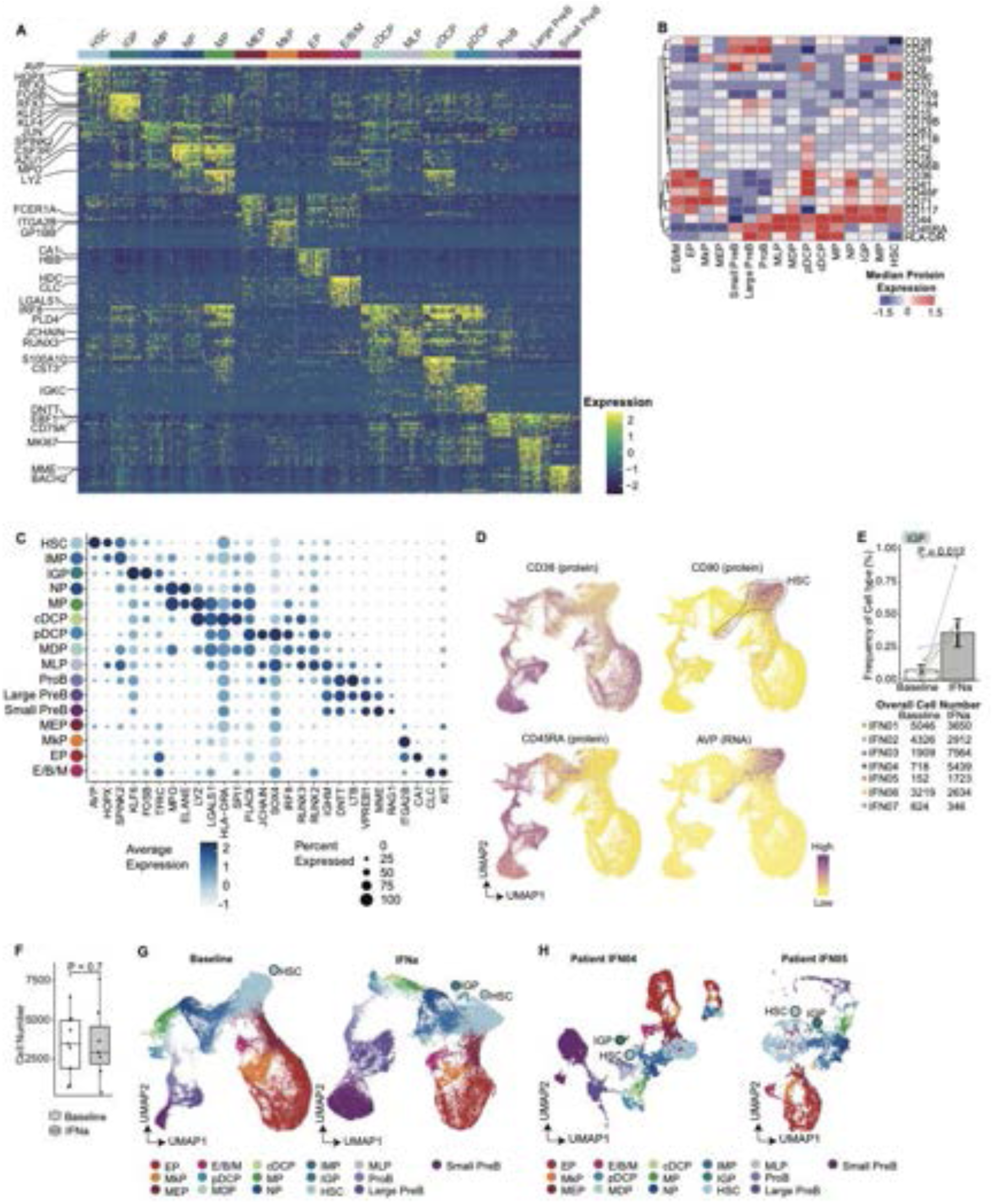
GoT-IM of baseline and IFN⍺-treated samples identifies HSPC populations. **A.** Heatmap of top 15 differentially expressed genes for each progenitor cell type. Cells of each progenitor type were down-sampled to the same number (n = 100 cells per cluster) for visualization. **B.** Heatmap showing median scaled expression of canonical HSPC protein markers from a representative patient IFN03. **C.** Dot plot showing expression levels of cell type specific gene markers in each progenitor subset. **D.** UMAP of sorted CD34^+^ HSPCs (n = 65,452 cells) highlighting CD38, CD90 and CD45RA protein expression and *AVP* RNA expression. **E.** Bar plot showing normalized IGP frequency from paired samples (n = 7 individuals) at baseline and IFN⍺ treatment. P-value from likelihood ratio test of LMM with/without treatment status (**methods**). **F.** Box plot showing the cell frequency at baseline and upon IFN⍺ treatment per sample (right). For IFN02, IFN04 and IFN05, the time-point powered with greater number of cells was selected. P-value from likelihood ratio test of LMM with/without treatment status (**methods**). **G.** UMAP of sorted CD34^+^ HSPCs overlaid with cell type assignment after separate integration for baseline (left, n = 33,877 cells) and IFN⍺-treated (right, n = 31,575 cells) samples. **H.** UMAP of CD34^+^ HSPCs overlaid with cell type assignment from representative samples IFN04 (n = 7,282 cells, left) and IFN05 (n = 3, 375 cells, right).

**fig. S3 (related to Fig. 1).**
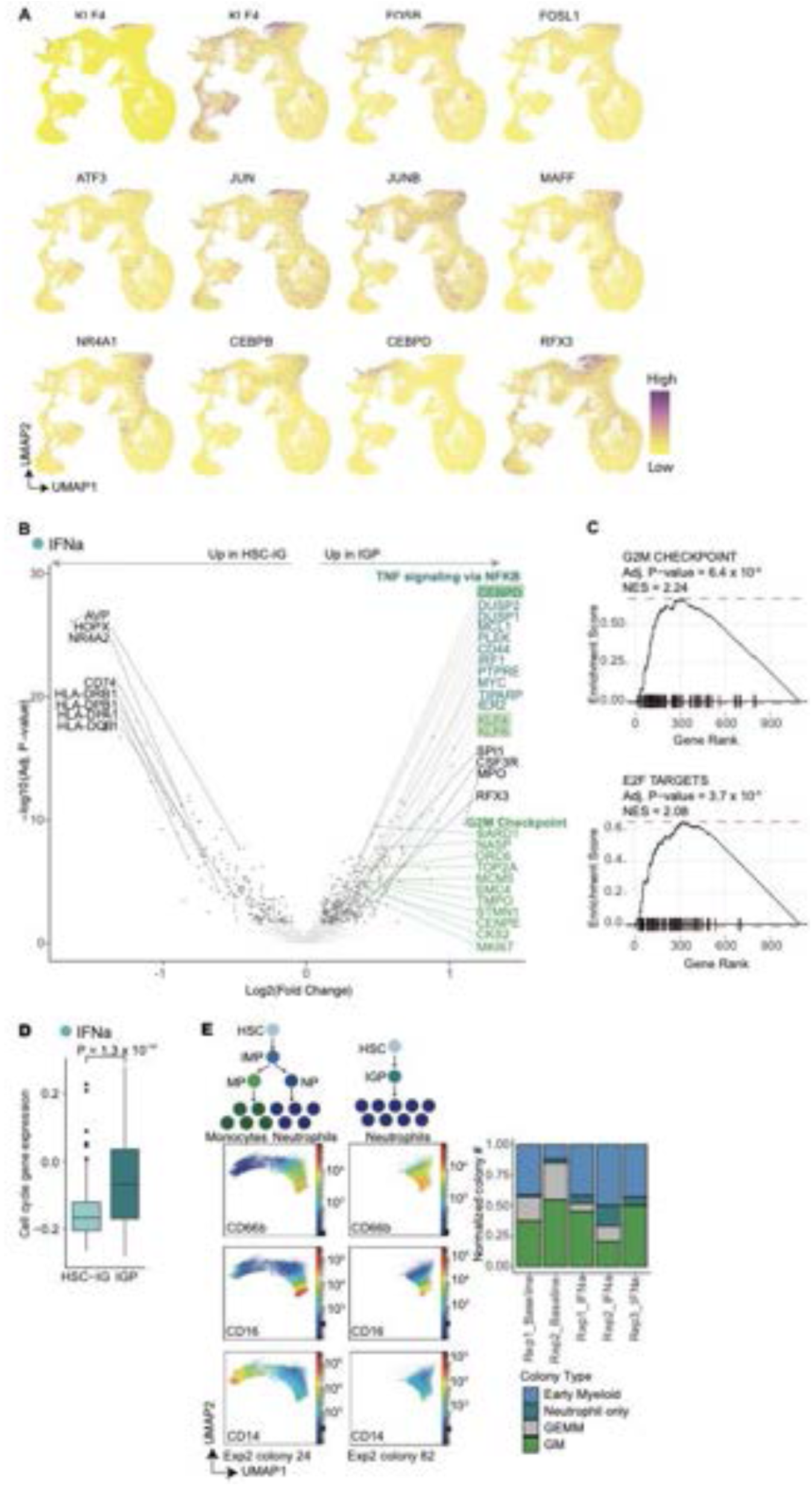
IFN⍺ induces a novel inflammatory granulocytic progenitor state. **A.** UMAP showing gene expression levels of differentially upregulated TFs in IGPs compared to IMPs**. B.** Left: Volcano plot of differentially expressed genes between IFN⍺-treated IGPs and HSC-IGs. P-values from likelihood ratio test of linear mixed modeling with/without cluster identity as a fixed effect variable (**methods**). Genes in blue represent genes enriched in the TNF⍺ signaling via NF-κB and those in green enriched in the G2M Checkpoint pathway (box representation is same as **Fig. 1E**). **C.** Pre-ranked gene set enrichment analysis using the MSigDB Hallmark collection of G2M Checkpoint and E2F Targets gene set. **D.** Cell cycle gene expression in IFN⍺-treated HSC-IGs vs IFN⍺-treated IGPs. P-value from likelihood ratio test of LMM with/without cluster identity. **E.** Schematic showing proposed model of HSC vs HSC-IG differentiation (top-left). UMAPs generated from flow-cytometric data showing representative colonies from single-cell differentiation assay from baseline and IFN⍺-treated BM HSCs **(methods)** depicting differentiation trajectories (bottom-left). Normalized frequency of colonies including early myeloid (EM), granulocytic (G), mixed granulocytic-monocytic (GM), mixed granulocytic, monocytic, early myeloid (GEMM) across single-cell differentiation assays from sorted baseline (n=2) and IFN⍺-treated (n=3) samples **(methods)** (right).

**fig. S4 (related to Fig. 2).**
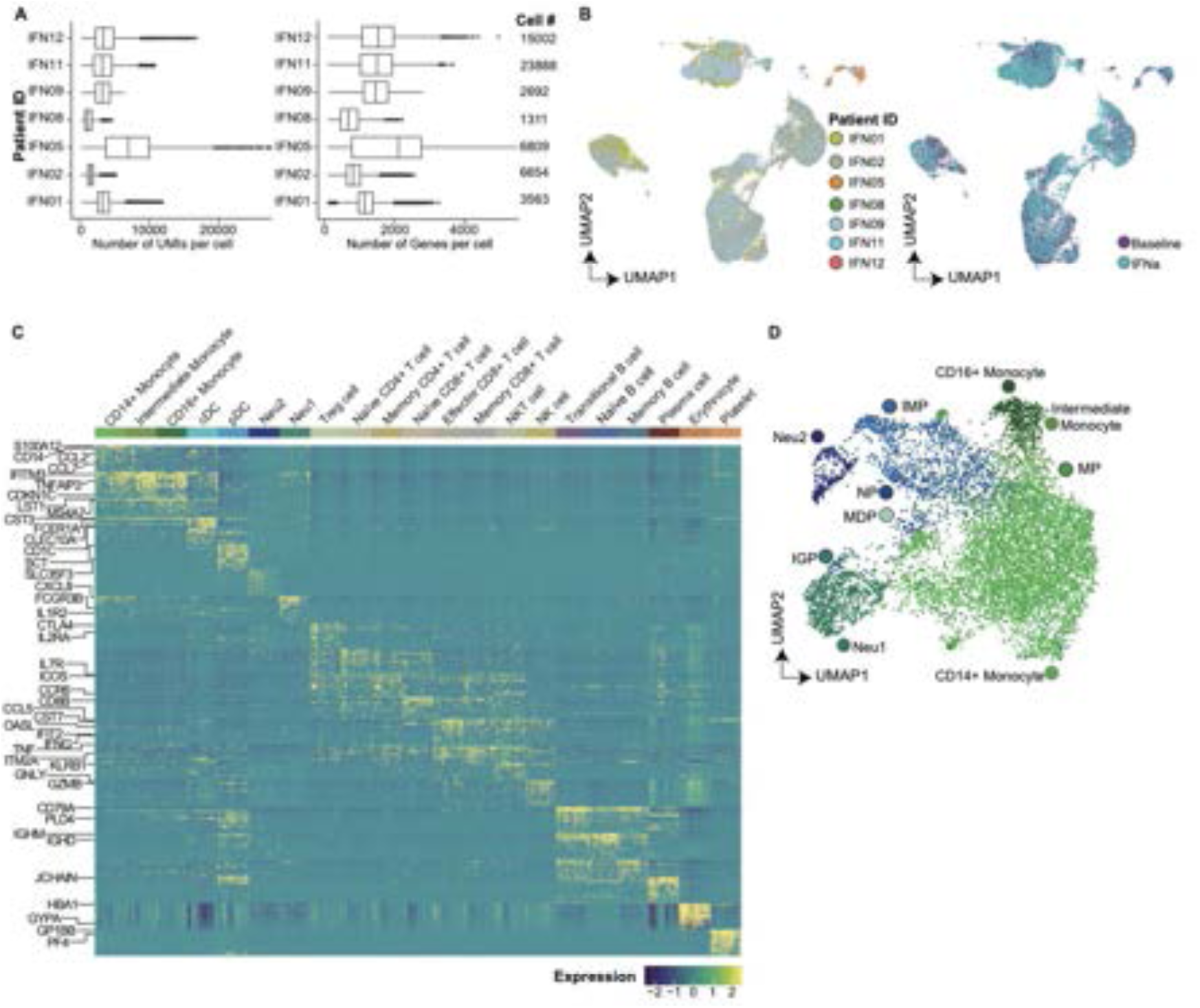
Genotyping of Transcriptomes integrated with immunophenotyping (GoT-IM) profiles thousands of CD34^-^ cells at baseline and following treatment with IFN⍺. **A.** Box plots showing number of UMIs (left panel), and genes (right panel) detected per cell in sorted CD34^-^ mature immune cells from each patient after filtering based on quality control (QC) metrics (**methods**). **B.** UMAP of sorted CD34^-^ immune cells (n = 59,912 cells) highlighted by patient ID (n = 7 individuals, left) and treatment status (right) after integration. **C.** Heatmap of top 15 differentially expressed genes for each immune cell type. Cells from each group was down-sampled to the same number (n = 75 cells per cluster) for visualization. **D.** Integrated UMAP of sorted CD34^+^ progenitors and CD34^-^ mature immune cells from representative samples (n = 4 individuals, IFN09-IFN12) highlighted by cell type assignment.

**fig. S5 (related to Fig. 3).**
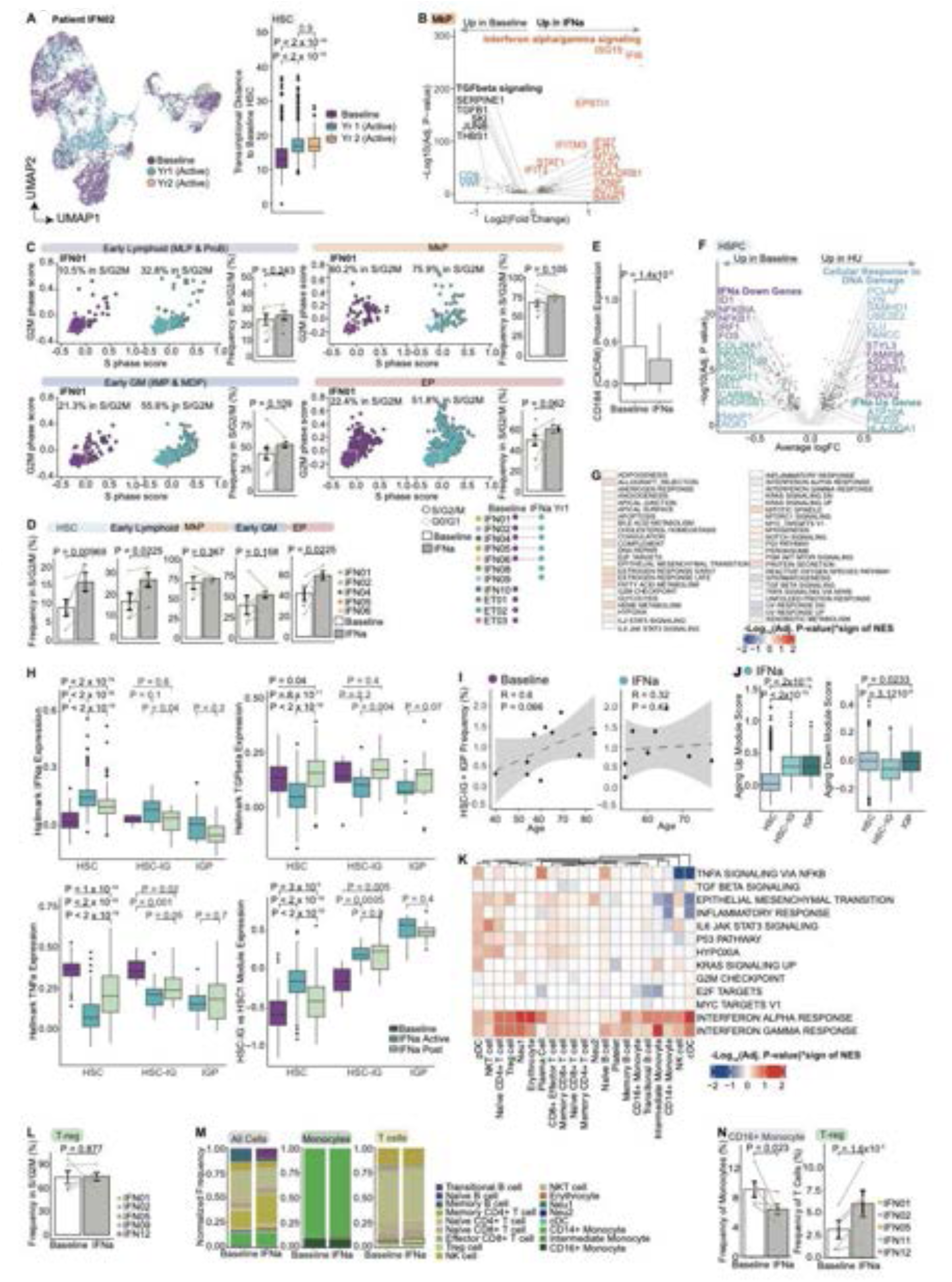
IFN⍺ induces HSC cell cycle entry. **A.** UMAP of CD34^+^ cells from patient IFN02 highlighting treatment status (n = 4,326 baseline cells, 2,912 IFN⍺-treated cells at year 1 and 541 IFN⍺-treated cells at year 2, left). Box plot showing transcriptional distance measurements between HSCs from each time-point and HSCs at baseline (right). Transcriptional distance corresponds to Euclidean distance of the first thirty principal components. P-values from Wilcoxon rank sum test, two-sided. **B.** Volcano plot showing differentially expressed genes between baseline and IFN⍺-treated MkPs. P-values from likelihood ratio test of linear mixed modeling (LMM) with/without treatment status (**methods**). Genes in black are enriched in the TGFβ signaling and those in orange enriched in the IFN⍺/γ response. Enrichment is based on pre-ranked gene set enrichment analysis (GSEA) using the MSigDB Hallmark collection. **C.** Cell cycle status of progenitor cells. For each cell type, left subpanel: Cell cycle gene expression in progenitor cells (representative patient IFN01, see **table S2** for cell numbers). Right subpanel: Frequencies of cells in G2/M/S phase as assessed in left subpanel (n = 11 baseline and 9 treated samples). P-values were derived from likelihood ratio test of LMM with/without treatment status. **D.** Frequencies of cells in G2/M/S phase in HSCs and progenitor cells at baseline and upon IFN⍺ treatment (n = 12 paired samples from 6 individuals). P-values were derived from likelihood ratio test of LMM with/without treatment status. **E.** *CXCR4* (CD184) protein expression in stem and early progenitor subsets (HSCs, IMPs, MLPs, MEPs and MDPs) at baseline and under IFN⍺ treatment. P value from likelihood ratio test LMM with/without treatment status. **F.** Volcano plot showing DE genes between baseline and HU treated HSPCs highlighting genes in cellular response to DNA damage (light-blue) the MSigDB GOBP collection, IFN⍺*^UP^* genes (teal) and IFN⍺*^DN^*genes (purple) from. P-values from likelihood ratio test of LMM with/without treatment status. **G.** Heatmap showing results of the pre-ranked gene set enrichment analysis comparing baseline and HU treated HSPCs. Values show the sign of the normalized enrichment score (NES) multiplied by -log10(Adjusted P-value). **H.** Gene expression of IFN⍺ response signature, TGFβ pathway and TNF⍺ signaling from the MSigDB Hallmark collection and HSC-IG vs HSC1 (**Fig. 1G**) signature at baseline, during and post-IFN⍺-treatment in HSCs (HSC1 and HSC2 subclusters from **Fig. 1G**), HSC-IGs and IGPs from a representative sample IFN05. **I.** Scatterplot showing correlation between patients’ age and their normalized HSC-IG and IGP frequency at baseline (left) and upon IFN⍺ treatment (right). **J.** Boxplots showing module gene expression of previously identified aging-specific upregulated genes (left) and downregulated (right) genes (*56*) in IFN⍺-treated HSCs (HSC1 and HSC2 subsets from **Fig. 1G**), HSC-IGs and IGPs. P-values from likelihood ratio test of LMM with/without cluster identity (**methods**). (*56*) in IFN⍺-treated HSCs (HSC1 and HSC2 subsets from **Fig. 1G**), HSC-IGs and IGPs. P-values from likelihood ratio test of LMM with/without cluster identity (**methods**). **K.** Heatmap showing results of the pre-ranked gene set enrichment analysis comparing baseline and during IFN⍺ treatment across CD34^-^ mature cells. Values show the sign of the normalized enrichment score (NES) multiplied by -log10(Adjusted P-value). **L.** Frequency of regulatory T cells in G2/M/S phase as assessed based on cell cycle gene expression at baseline and upon IFN⍺ treatment. P-values were derived from likelihood ratio test of LMM with/without treatment status. **M.** Normalized frequency of cell types within all CD34^-^ mature immune cells (left), monocytes (middle) and T cells (right) at baseline and active IFN⍺ treatment. Cells from each treatment status and individual were down-sampled to the same number (n = 500 cells per treatment status per sample, 10 baseline and IFN⍺-treated paired samples from 5 individuals). For IFN02 and IFN05, treated time-point powered with greater number of cells was selected. **N.** Normalized frequency of CD16^+^ monocytes and regulatory T (Treg) cells at baseline and upon IFN⍺ treatment (n = 10 samples from 5 individuals). For IFN02 and IFN05, the treated time-point powered with greater number of cells was selected. P-values from likelihood ratio test of LMM with/without treatment status.

**fig. S6 (related to Fig. 4).**
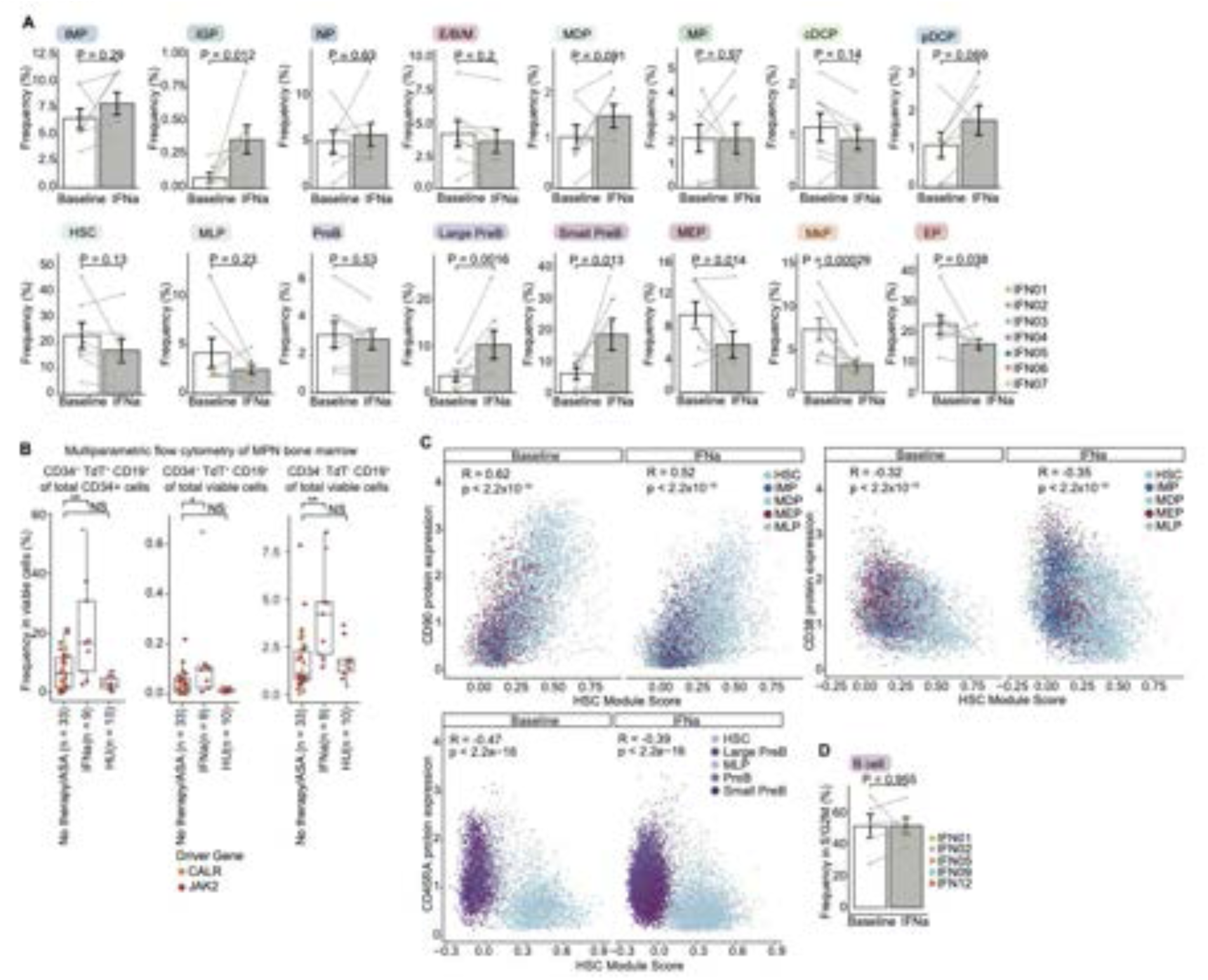
IFN⍺ induces lymphoid differentiation. **A.** Bar plot showing normalized cell type frequencies at baseline and during IFN⍺ treatment (n = 14 paired samples from 7 individuals). For IFN02, IFN04 and IFN05, the time-point powered with greater number of cells was selected. P-values from likelihood ratio test of linear-mixed modeling (LMM) with/without treatment status (**methods**). **B.** Box plots showing cell frequencies of B-lymphoid progenitors and B cells from bone marrow of patients with early stage MPN treated with IFN⍺ and HU treatment (n = 9 and 10 respectively) and without treatment (n = 33 samples), as determined by multiparametric flow cytometry. P-values from Wilcoxon rank sum test, two-sided. **C.** Scatter plot showing correlation between HSC module expression (based on differentially expressed genes in HSC cluster, **fig. S2A**) and protein expression of canonical stem/progenitor markers i.e., CD90 (top-left), CD38 (top-right) and CD45RA (bottom) in stem and progenitor subsets. P-value from F-test, Pearson correlation. Shading denotes 95% confidence interval. **D.** Frequency of B cells in G2/M/S phase as assessed based on cell cycle gene expression at baseline and upon IFN⍺ treatment. P-values were derived from likelihood ratio test of LMM with/without treatment status.

**fig. S7 (related to Fig. 5).**
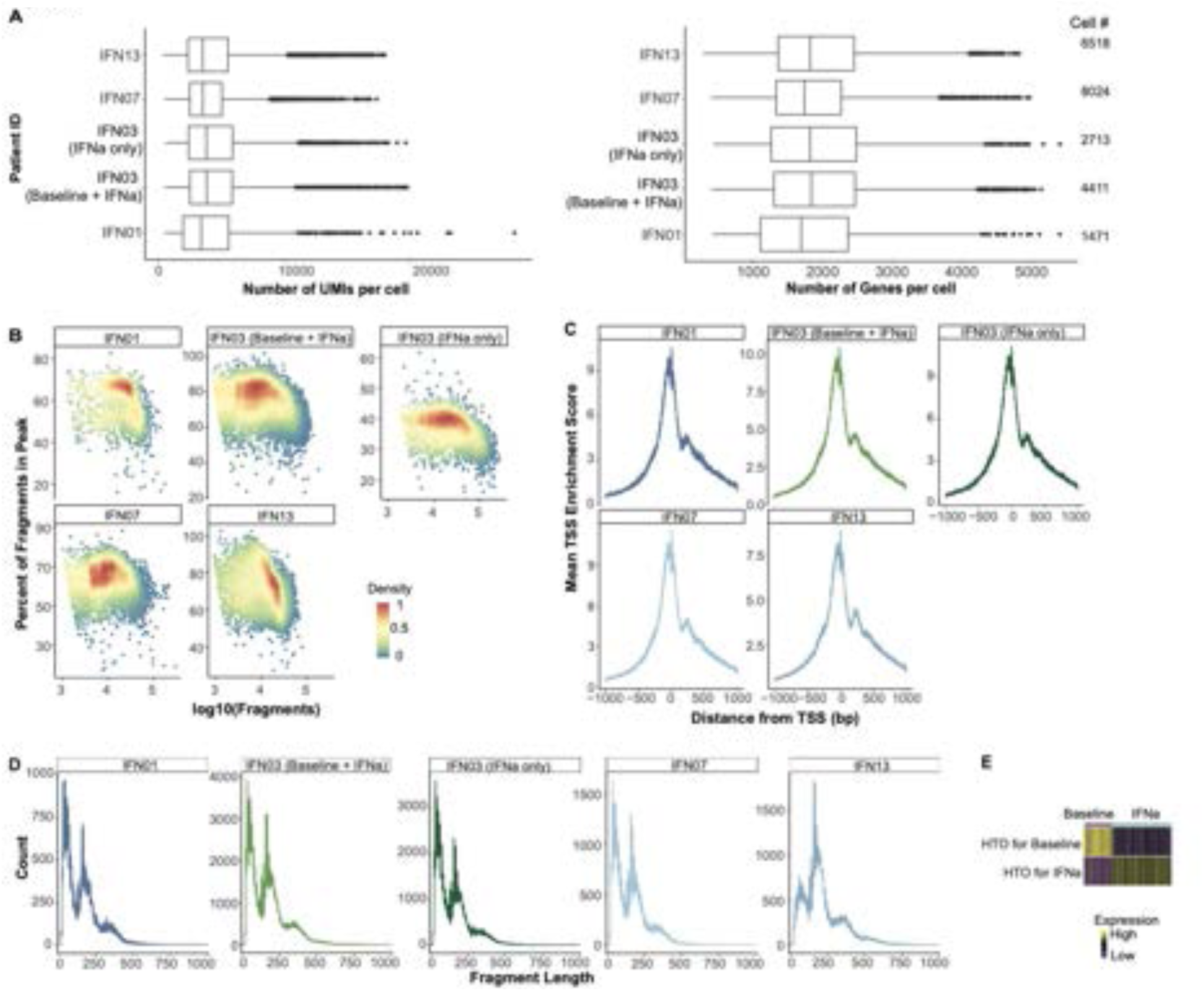
GoT-ATAC captures genotyping, snRNA-seq and snATAC-seq data for CD34^+^ HSPCs. **A.** Box plots showing number of UMIs (left) and genes (right) detected per cell in sorted CD34^+^ hematopoietic stem and progenitors from each patient after filtering based on quality control (QC) metrics (**methods**) from GoT-ATAC experiments. **B.** Density plot comparing percentage of snATAC fragments within peaks to the total number of fragments detected per sample (n = 7 samples from 4 individuals, additional IFN03 IFN⍺-treated cells also sequenced separately). **C.** Distribution of mean TSS enrichment score at each position relative to the TSS per sample. **D.** Average distribution of fragment length per sample. **E.** Heatmap showing HTO expression level for baseline and IFN⍺-treated cells from representative IFN01 (n = 1471 cells).

**fig. S8 (related to Fig. 5).**
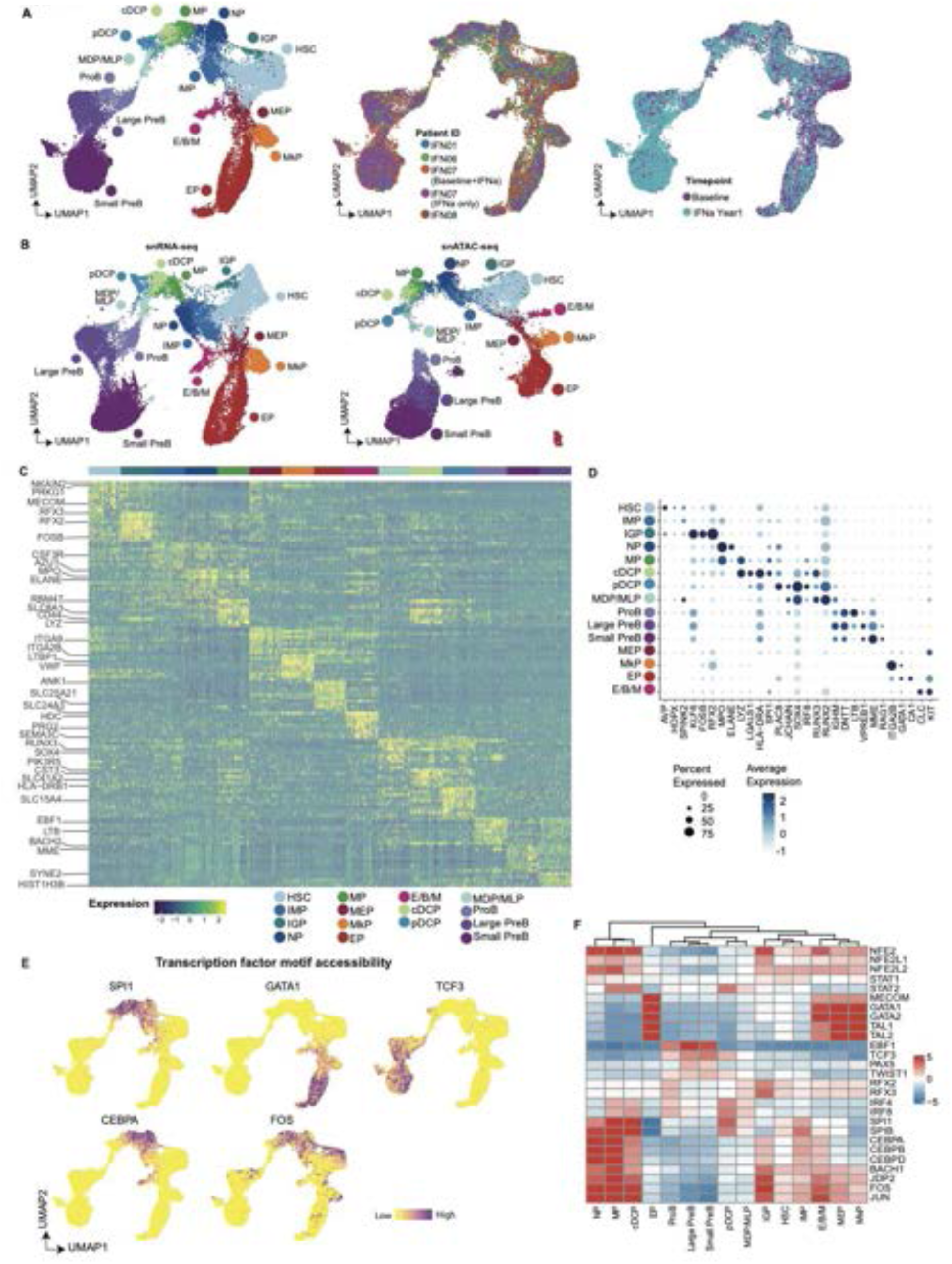
GoT-ATAC identifies the novel inflammatory granulocytic progenitor population. **A.** UMAP of sorted CD34^+^ stem and progenitors (n = 23,137 cells, 7 samples from 4 individuals), with cell type (left), patient ID (middle) and treatment status (right) using weighted-nearest neighbor (WNN) analysis of snRNA-seq and snATAC-seq data (**methods**). **B.** UMAP of CD34^+^ cells based on snRNA-seq (left) and snATAC-seq (right) data, overlaid with cell type assignment. **C.** Heatmap of top 15 differentially expressed genes for each HSPC type. Cells of each progenitor type were down-sampled to the same number (n = 100 cells per cluster). **D.** Dot plot showing expression levels of cell type-specific gene markers in each progenitor subset. **E.** UMAP based on weighted nearest neighbor (WNN) analysis (n = 23,137 cells) highlighting TF motif accessibility. TF accessibility scores added with AddChromatinModule function in Signac. **F.** Heatmap showing cell type specific TF accessibility scores.

**fig. S9 (related to Fig. 5).**
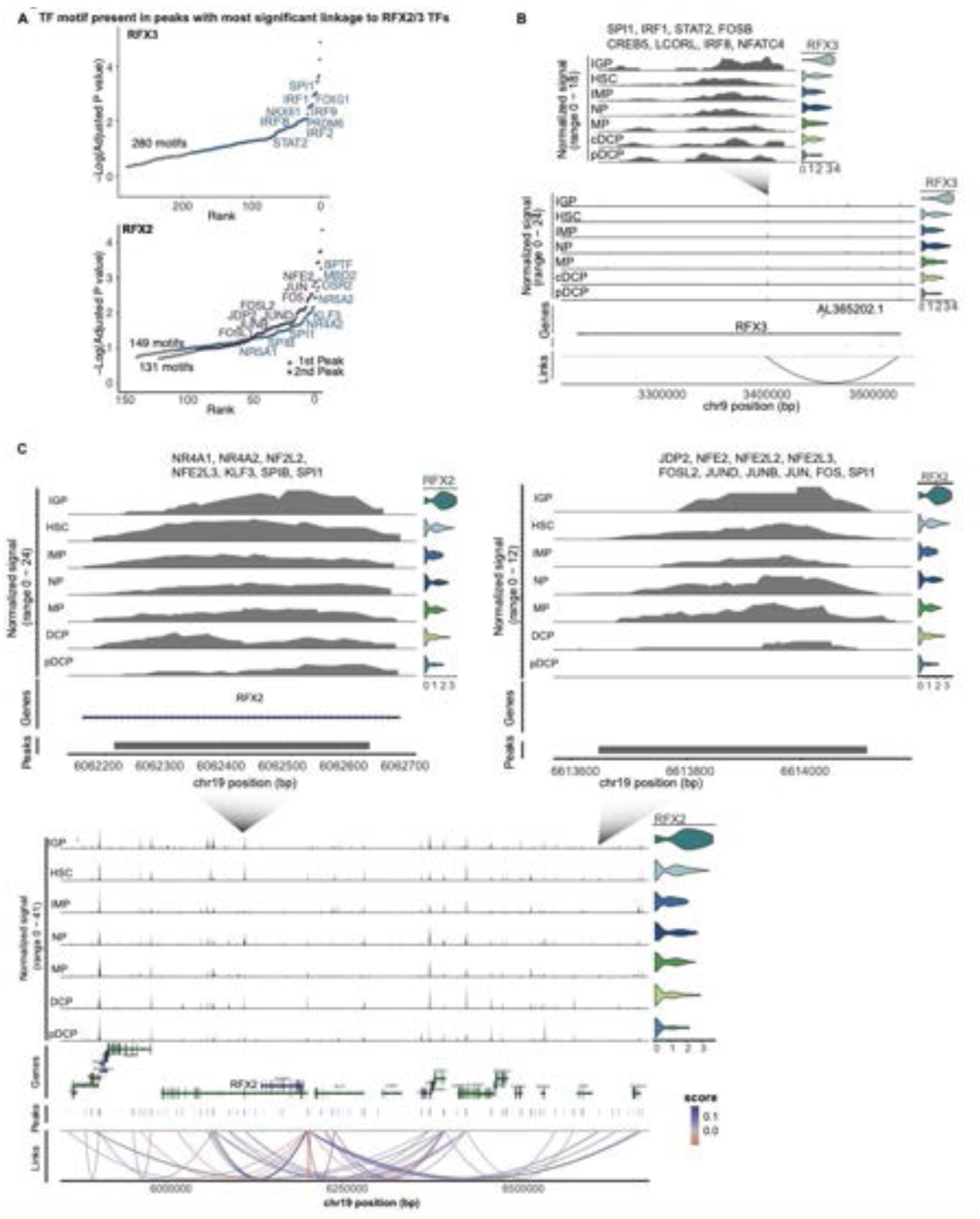
Transcription factor activities in inflammatory granulocytic progenitors. **A.** Ranked transcription factor (TF) motifs in the most significant individual positive regulatory peak of RFX3 (top) and RFX2 (bottom, 1st peak: upstream of TSS, 2nd peak: downstream of TSS) identified using motif scanning with FIMO for a representative sample (IFN03, **methods)**. **B.** Chromatin accessibility track (left) of the regulatory region of *RFX3* (representative example from IFN03). Violin plots (right) display gene expression level of *RFX3*. **C.** Chromatin accessibility tracks of regulatory regions of *RFX2* (representative example from IFN03, bottom) and distal region enriched with the two most significant positively regulating loci (top-left, top-right). Violin plots display gene expression level of *RFX2*.

**fig. S10 (related to Fig. 5).**
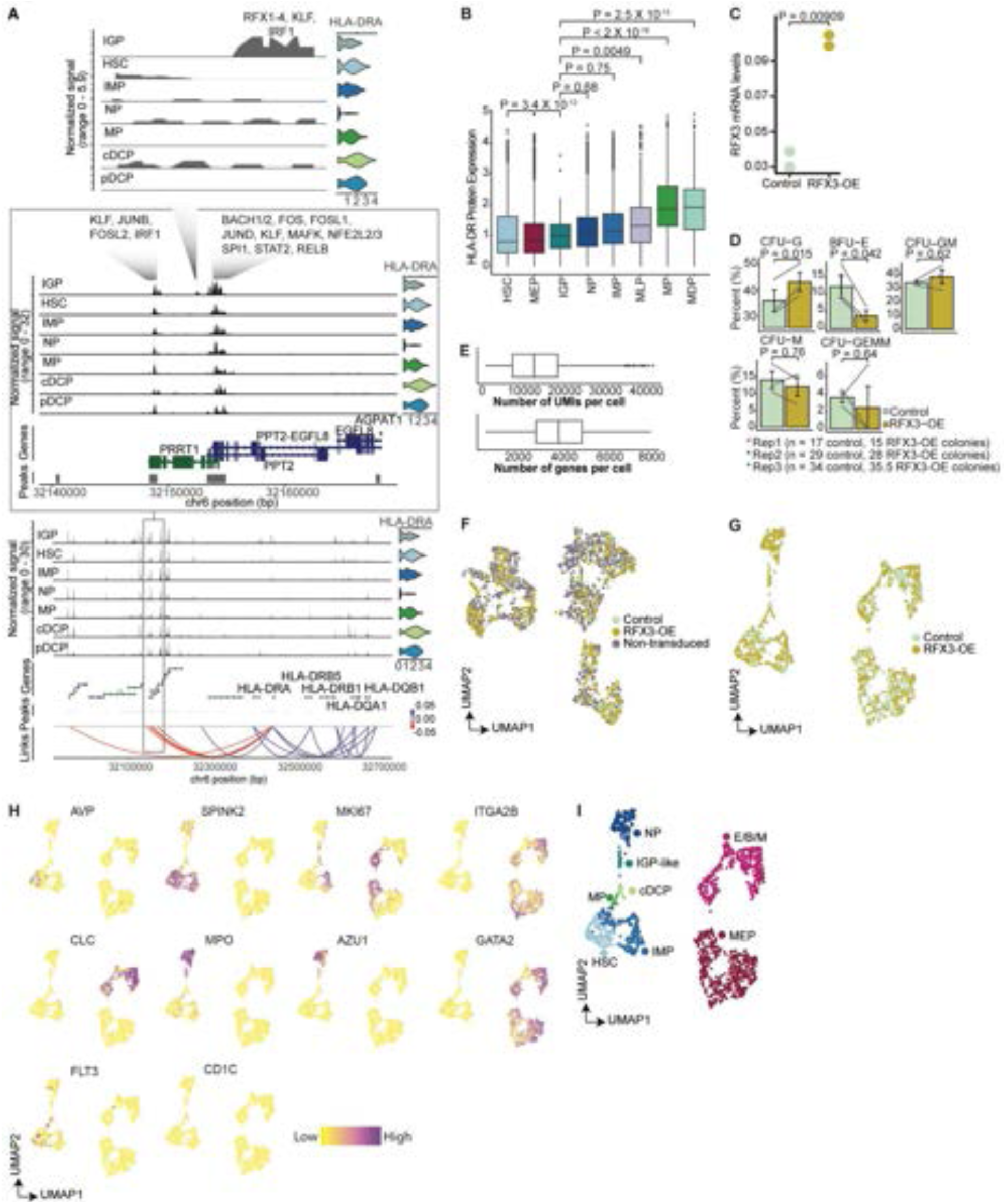
RFX3 overexpression induces an IGP-like cell state. **A.** Chromatin accessibility tracks of regulatory regions of *HLA-DRA1* (bottom), distal region enriched with negative regulatory loci (inset), and IGP-specific regulatory locus (top, representative example from IFN07). Violin plots display gene expression level of *HLA-DRA1*. **B.** Box plots showing normalized expression of HLA-DR protein expression. P-values from likelihood ratio tests of linear-mixed modeling (LMM) with/without cell type identity. **C.** RFX3 overexpression lentiviral vector validated in K562 cells by assessing *RFX3* mRNA levels by RT-QPCR (n = 2 independent experiments). mRNA levels correspond to RFX3 Ct values normalized to TBP Ct values. P-value from t-test. **D.** Normalized frequency of erythroid (BFU-E), granulocytic (CFU-G), granulo-monocytic (CFU-GM), monocytic (CFU-M) and myeloid (CFU-GEMM) colonies grown from RFX3-OE CD34^+^ umbilical cord blood (UCB) cells in methylcellulose-based CFU assays compared to control CD34^+^ UCB cells. P-value from likelihood ratio test linear-mixed modeling (LMM) with/without RFX3 overexpression. **E.** Box plots showing number of UMIs (top) and genes (bottom) detected per cell in sorted CD34^+^ UCB cells after filtering based on quality control (QC) metrics (**methods**) from GoT-IM experiments. **F.** UMAP of sorted CD34^+^ UCB cells (n = 2,609 cells) highlighted by three transduction statuses. **G.** UMAP of RFX3-OE and mCherry (control) subsets (n = 1,520 cells) highlighted by their status. **H.** UMAP of RFX3-OE and control subset showing gene expression levels of cell type specific gene markers for HSPCs. **I.** UMAP of RFX3-OE and control subset, highlighting cell type assignments.

**fig. S11 (related to Fig. 5).**
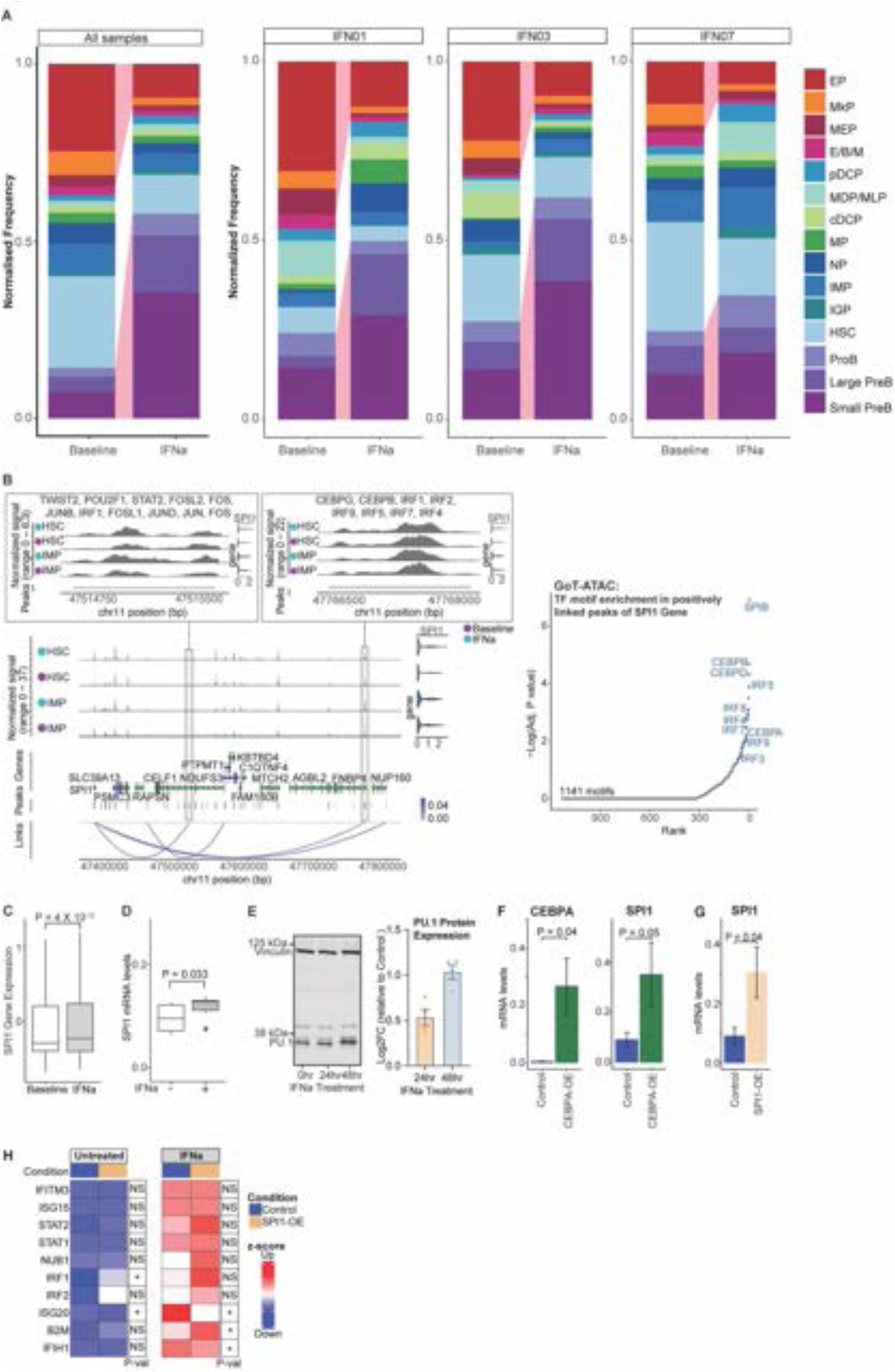
PU.1 is the master regulator of IFN⍺-mediated lymphoid differentiation and remodeling of hematopoiesis. **A.** Left: Normalized cell frequencies of progenitor subsets at baseline and after IFN⍺ treatment from all GoT-ATAC samples (n = 4 individuals). Cells from each treatment status and individual were down-sampled to the same number (n = 100 cells per treatment status per sample). Right: Cell frequency distribution as in left panel for patients IFN01, IFN03 and IFN07. **B.** Chromatin accessibility tracks of regulatory regions of *SPI1* (representative example from IFN07, bottom-left), distal region enriched with the two most significant positively regulating loci (top-left). Violin plots display gene expression level of *SPI1*. Ranked TF motif enrichment of all positive regulatory peaks of the SPI1 gene, relative to background peaks using the hypergeometric test across three samples IFN01, IFN03 and IFN07 (right, see **methods**). **C.** SPI1 gene expression in stem and early progenitors (HSCs, IMPs, MLPs, MEPs and MDPs) at baseline and upon IFN⍺ treatment. P-value from likelihood ratio test linear-mixed modeling (LMM) with/without treatment status. D. *SPI1* mRNA levels in K562 cells treated with IFN⍺ *in vitro* for 24 hours (assessed by RT-QPCR, n=6 independent experiments). mRNA levels correspond to SPI1 Ct values normalized to TBP Ct values. P-values from likelihood ratio test of LMM with/without treatment status. **E.** Representative western blot showing PU.1 protein levels in K562 cells treated with IFN⍺ *in vitro* across for 0, 24 and 48 hours with vinculin as loading control (left). Log-fold change analysis of quantified PU.1 protein levels based on luminescence intensity of western blot bands across three replicates (right) (**methods**). F. *CEBPA* and *SPI1* mRNA levels in K562 cells upon expression of control or CEBPA-OE lentiviral vectors (assessed by RT-QPCR, n=3 independent experiments). mRNA levels correspond to Ct values of gene target normalized to TBP Ct values (2^−ΔΔCt^). P-values from likelihood ratio test of LMM with/without CEBPA overexpression. G. *SPI1* mRNA levels in K562 cells upon expression of control or SPI1-OE lentiviral vectors (assessed by RT-QPCR, n=2-3 independent experiments). mRNA levels correspond to Ct values of gene target normalized to TBP Ct values (2^−ΔCt^). P-values from likelihood ratio test of LMM with/without SPI1overexpression. **H.** Heatmap depicting scaled mRNA levels(2^−ΔCt^) of IFN-related genes in K562 cells upon expression of control or SPI1-OE lentiviral vectors at baseline and after 24-hour IFN⍺ treatment (assessed by RT-QPCR, n = 4-7 independent experiments). Scaled across all data points for each gene. P-values from likelihood ratio tests of LMM with/without SPI1 overexpression status.

**fig. S12 (related to Fig. 6).**
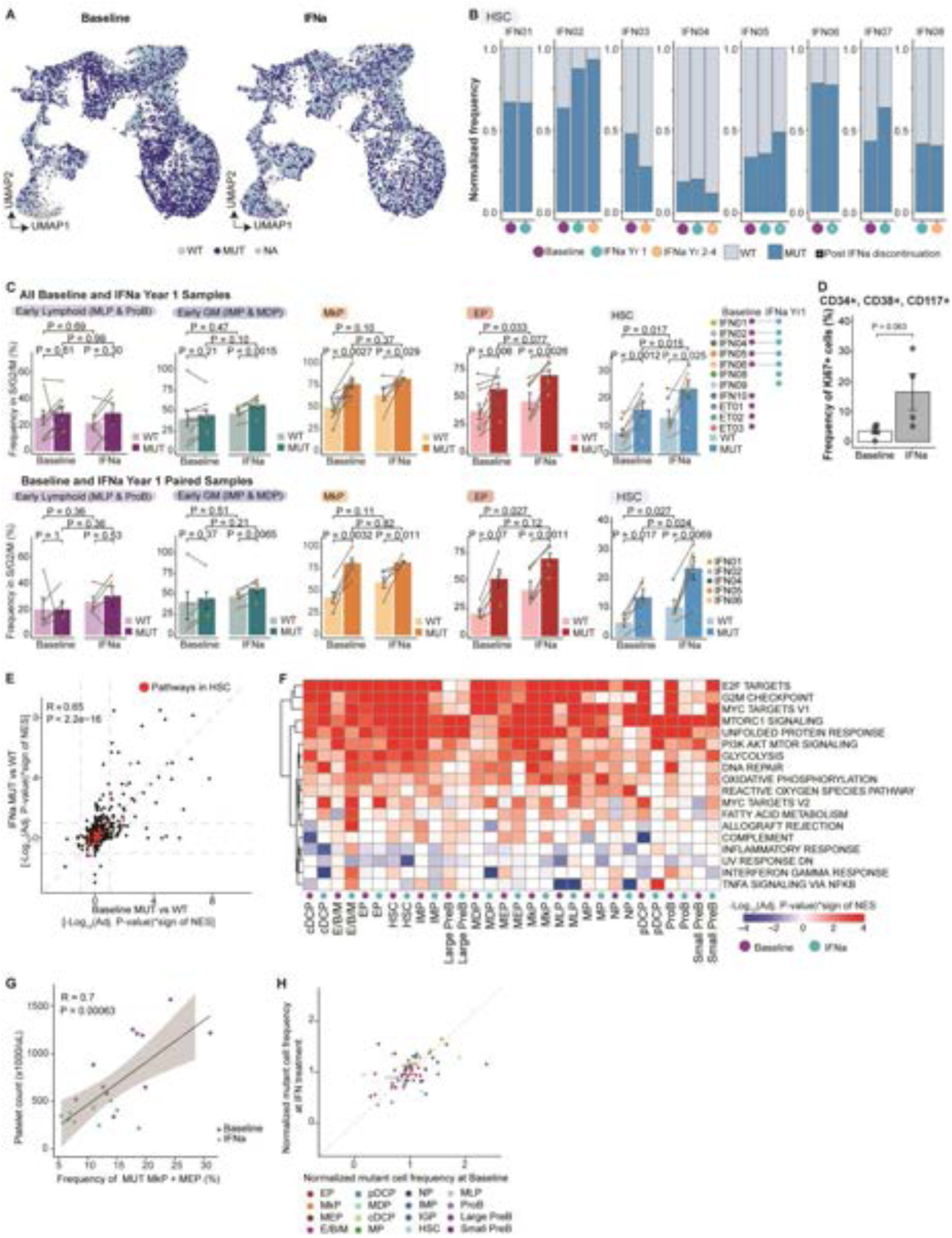
*CALR* mutations modify the effects of IFN⍺ signaling. **A.** UMAP of sorted CD34^+^ stem and progenitors at baseline and after IFN⍺ treatment from GoT-IM experiments with *CALR* mutation status highlighted. Cells from each sample were down-sampled to the same number for each mutation status (n = 1,000 cells from each mutation status per sample). **B.** Normalized frequencies of CALR-mutated (MUT) and wildtype (WT) HSCs at each time-point (n= 8 individuals with at least 2 time-points). **C.** Bar plots showing frequencies of MUT and WT cells in G2/M/S phase as assessed in **Fig. 3C** for all CD34^+^ GoT-IM samples (n = 11 baseline and 9 IFN⍺ year 1 samples, top) and paired CD34^+^ GoT-IM samples (n = 12 samples from 6 individuals, bottom). P-values were derived from likelihood ratio test of linear mixed modeling (LMM) with/without treatment status. **D.** Frequencies of Ki67^+^ myeloid cells before and after IFN⍺ treatment. P-values from Wilcoxon rank sum test, two-sided. **E.** Scatter plot showing pathways from pre-ranked DE gene set enrichment analysis comparing mutated (MUT) versus wildtype (WT) cells at baseline and after IFN⍺ treatment. Values show the sign of the normalized enrichment score (NES) multiplied by -log10(Adjusted P-value). Pathways in red are present in HSCs. P-value from F-test, Pearson correlation. Shading denotes 95% confidence interval. **F.** Heatmap showing results of the pre-ranked gene set enrichment analysis of genes DE between MUT and WT cells at baseline and after IFN⍺ treatment. **G.** Platelet counts versus frequencies of MUT MEPs and MkPs. P-value from F-test, Pearson correlation. Shading denotes 95% confidence interval. **H.** Normalized mutant cell frequency at baseline versus after IFN⍺ treatment for GoT-IM CD34^+^ samples (cell type clusters with at least 10 genotyped cells were within each sample were used).

**fig. S13 (related to Fig. 6).**
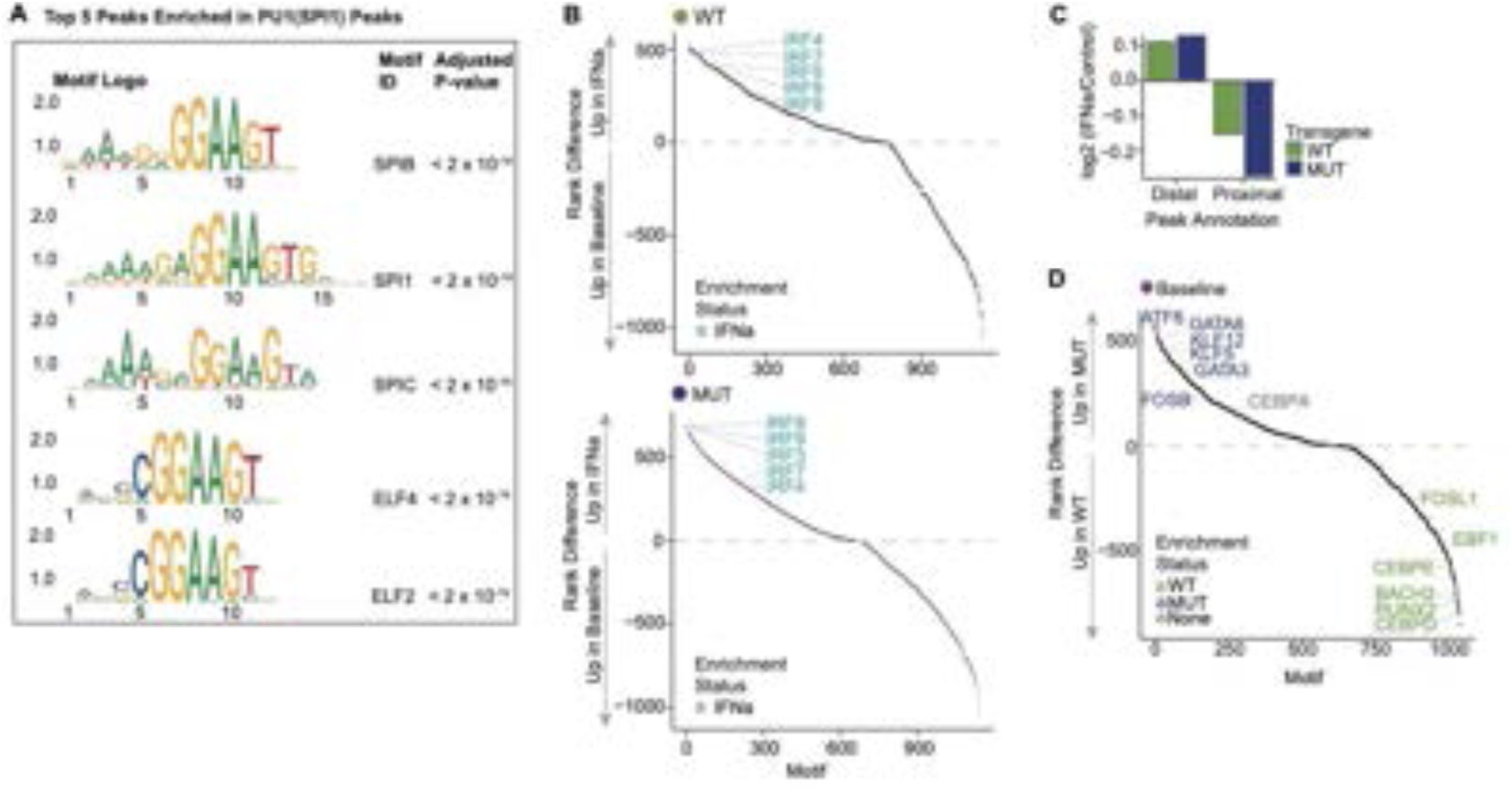
*CALR* mutated cells display enhanced PU.1 binding activity at regulatory regions with distinct cooperating TF binding sites. **A.** Top five hits from Ranked transcription factor (TF) motif enrichment of peaks captured from CUT&RUN targeting PU.1 motif with SEA v5.5.5. **B.** Differential TF motif enrichment between baseline and IFN⍺-treated focusing on exclusive PU.1 peaks in WT (top) and MUT cells (bottom). Analyses with HOMER. Highlighted are IRF TFs (blue). **C.** Bar plot showing difference in the number of distal and proximal PU.1 peaks between control and IFN⍺-treated UT7-MPL cells. Peaks less than 500 bp from TSS were considered proximal. **D.** Differential TF motif enrichment between MUT and WT exclusive PU.1 peaks in baseline cells. TFs enriched in differentially accessible PU.1 peaks for MUT and WT cells are highlighted in blue and green respectively.

**fig. S14 (related to Fig. 6).**
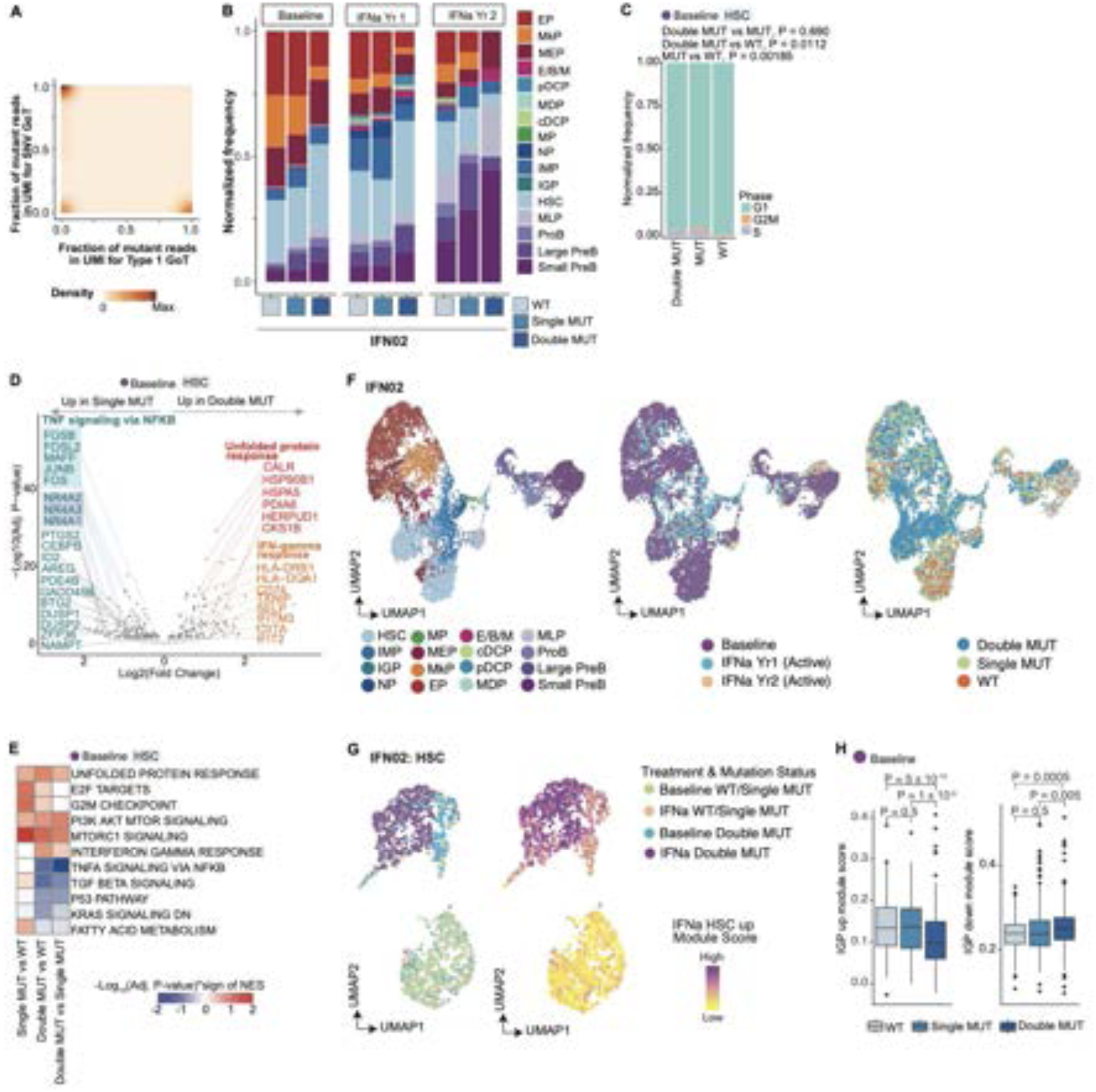
IFN⍺ perturbs clonal evolution via the IGP differentiation program. **A.** Relative density of proportion of Type 1-mutant reads versus SNV-mutant reads in UMIs captured via GoT, showing mutual exclusivity. **B.** Normalized frequencies of each progenitor subset among WT, single MUT (Type 1 *CALR*) and double MUT (Type 1 and SNV mutations in *CALR*) cell populations at each time-point for IFN02 (n = 7,779 cells). **C.** Bar plots showing normalized frequency of baseline HSCs in G1/G2/M/S phases as assessed in **Fig. 3C**. P-values from Fisher’s exact test between mutation status and cell-cycle entry status (G2/M/S vs G1). **D.** Volcano plot showing genes differentially expressed (DE) between single mutant and double mutant HSCs at baseline (n = 890 genotyped HSCs). DE genes identified using logistic regression model (**methods**). Genes highlighted in blue are enriched in the TNF⍺ signaling via NF-κB, red in the unfolded protein response, and orange in the IFNγ response (box representation is same as **Fig. 1E**). **E.** Heatmap showing results of the pre-ranked gene set enrichment analysis of genes DE between HSC clones at baseline. Values correspond to the sign of the normalized enrichment score (NES) multiplied by the -log10(Adjusted P-value)**. F.** UMAP of IFN02 patient based on scRNA-seq data highlighting cell types (left), treatment status (middle) and mutation status (right). **G.** UMAP of HSCs from IFN02 (n = 1,879 cells) overlaid with treatment status and HSC clones (left) and module score for HSC-specific IFN⍺-induced upregulated genes (right). **H.** Box plots showing IGP-specific signature score in HSC clones at baseline in IFN02. Scores calculated using IGP-upregulated or downregulated genes (left and right panels, respectively, **methods**). P-values from Wilcoxon rank sum test, two-sided.

## Supplementary Tables

**Table S1.** Summary of patients’ clinical history, pathology and laboratory data, and mutation status.

**Table S2.** Number of cells for each cell type in GoT-IM CD34^+^ compartment.

**Table S3.** Differential gene expression analyses for IGP versus IMP and treated IGP versus treated HSC1 via the linear mixed modeling framework.

**Table S4.** Gene set enrichment analysis of genes differentially expressed between IGPs versus IMPs and treated IGPs versus treated HSC-IG.

**Table S5.** RNA velocity analysis with scVelo between HSPC subpopulations. Cluster-to-cluster transition and connectivity scores calculated based on velocity graph based on pseudotime values. **Table S6.** Number of cells for each cell type in GoT-IM CD34^-^ compartment.

**Table S7.** Differential gene expression analyses for Neu1 versus Neu2 subsets via the linear mixed modeling framework. Gene set enrichment analysis of genes differentially expressed between Neu1 versus Neu2 subsets.

**Table S8.** Differential gene expression analysis between baseline and IFN⍺-treated HSPC subsets. **Table S9.** Gene set enrichment analysis of genes differentially expressed between baseline and IFN⍺-treated HSPC subtypes.

**Table S10.** Differential gene expression analysis between baseline and IFN⍺-treated CD34^-^ mature cells subsets.

**Table S11.** Gene set enrichment analysis of genes differentially expressed between baseline and IFN⍺-treated CD34^-^ mature cells subsets.

**Table S12.** Clinical data multi-parametric flow cytometry data of bone marrow aspirates from patients with early-phase MPN with IFN⍺/HU and without treatment.

**Table S13.** Number of cells identified for each cell type in each sample via GoT-ATAC.

**Table S14.** Differential transcription factor motif enrichment analysis between IGPs and HSCs under IFN⍺ treatment.

**Table S15.** Motifs identified in peaks linked with RFX2/3 via motif scanning with FIMO.

**Table S16.** Over-enrichment motif analysis for peaks linked with AP-1 genes and SPI1 gene.

**Table S17.** Motifs identified in peaks linked with MHC class II genes.

**Table S18.** Genes positively regulated by RFX2 and RFX3 using gene-peak cis-association.

**Table S19.** Differential gene expression analyses for IGP-like vs IMP subsets in CD34^+^ cells with RFX3 overexpression. Gene set enrichment analysis of genes differentially expressed between IGP-like vs IMP subsets.

**Table S20.** Differential transcription factor motif enrichment analysis between baseline and IFN⍺-treated HSCs.

**Table S21.** Differential gene expression analysis between *CALR*-mutated and wildtype cells at baseline in HSPC subsets.

**Table S22.** Gene set enrichment analysis of genes differentially expressed between *CALR*-mutated and wildtype cells at baseline in HSPC subsets.

**Table S23.** Differential gene expression analysis between *CALR*-mutated and wildtype cells after IFN⍺ treatment in HSPC subsets.

**Table S24.** Gene set enrichment analysis of genes differentially expressed between *CALR*-mutated and wildtype cells after IFN⍺ treatment in HSPC subsets.

**Table S25.** Differential transcription factor motif enrichment analysis between MUT vs WT stem and early progenitor cells at baseline and upon IFN⍺ treatment.

**Table S26.** Motifs identified in PU.1 bound peaks captured from CUT&RUN data of CALR-mutated versus WT UT-7 cells.

**Table S27.** Differential TF motif enrichment in MUT and WT exclusive PU.1 peaks at baseline and upon IFN⍺ treatment.

**Table S28.** Differential TF motif enrichment in IFN⍺ and Baseline exclusive PU.1 peaks for MUT and for WT.

**Table S29.** Differential gene expression analysis between HSC clones from IFN02 at baseline. **Table S30.** Gene set enrichment analysis of genes differentially expressed between HSC clones from IFN02.

**Table S31.** List of antibodies used for FACS, CITE-seq, and Cell Hashing and primer sequences used in GoT-IM, GoT-ATAC, and RT-PCR.

## References

1. S. Jaiswal et al., Clonal Hematopoiesis and Risk of Atherosclerotic Cardiovascular Disease. N Engl J Med 377, 111–121 (2017).

2. S. Jaiswal et al., Age-related clonal hematopoiesis associated with adverse outcomes. N Engl J Med 371, 2488–2498 (2014).

3. L. Busque et al., Recurrent somatic TET2 mutations in normal elderly individuals with clonal hematopoiesis. Nat Genet 44, 1179–1181 (2012).

4. S. Avagyan et al., Resistance to inflammation underlies enhanced fitness in clonal hematopoiesis. Science 374, 768–772 (2021).

5. A. Heyde et al., Increased stem cell proliferation in atherosclerosis accelerates clonal hematopoiesis. Cell 184, 1348–1361.e1322 (2021).

6. M. Meisel et al., Microbial signals drive pre-leukaemic myeloproliferation in a Tet2-deficient host. Nature 557, 580–584 (2018).

7. Z. Cai et al., Inhibition of Inflammatory Signaling in Tet2 Mutant Preleukemic Cells Mitigates Stress-Induced Abnormalities and Clonal Hematopoiesis. Cell Stem Cell 23, 833–849.e835 (2018).

8. J. J. Trowbridge, D. T. Starczynowski, Innate immune pathways and inflammation in hematopoietic aging, clonal hematopoiesis, and MDS. J Exp Med 218, (2021).

9. M. A. Essers et al., IFNalpha activates dormant haematopoietic stem cells in vivo. Nature 458, 904–908 (2009).

10. E. M. Pietras et al., Re-entry into quiescence protects hematopoietic stem cells from the killing effect of chronic exposure to type I interferonsIFN-1 effects on HSC function. J Exp Medicine 211, 245–262 (2014).

11. M. T. Baldridge, K. Y. King, N. C. Boles, D. C. Weksberg, M. A. Goodell, Quiescent haematopoietic stem cells are activated by IFN-gamma in response to chronic infection. Nature 465, 793–797 (2010).

12. T. N. Rao et al., JAK2-V617F and interferon-α induce megakaryocyte-biased stem cells characterized by decreased long-term functionality. Blood 137, 2139–2151 (2021).

13. M. Mosca et al., Inferring the dynamics of mutated hematopoietic stem and progenitor cells induced by IFNα in myeloproliferative neoplasms. Blood 138, 2231–2243 (2021).

14. J. L. Spivak, Myeloproliferative Neoplasms. N Engl J Med 377, 895–896 (2017).

15. T. A. Knudsen et al., Genomic profiling of a randomized trial of interferon-α vs hydroxyurea in MPN reveals mutation-specific responses. Blood Adv 6, 2107–2119 (2022).

16. A. S. Nam et al., Somatic mutations and cell identity linked by Genotyping of Transcriptomes. Nature 571, 355–360 (2019).

17. J. Czech et al., JAK2V617F but not CALR mutations confer increased molecular responses to interferon-α via JAK1/STAT1 activation. Leukemia 33, 995–1010 (2019).

18. T. Klampfl et al., Somatic mutations of calreticulin in myeloproliferative neoplasms. N Engl J Med 369, 2379–2390 (2013).

19. A. Yacoub et al., Pegylated interferon alfa-2a for polycythemia vera or essential thrombocythemia resistant or intolerant to hydroxyurea. Blood 134, 1498–1509 (2019).

20. J. Mascarenhas et al., A randomized phase 3 trial of interferon-α vs hydroxyurea in polycythemia vera and essential thrombocythemia. Blood 139, 2931–2941 (2022).

21. M. Stoeckius et al., Cell Hashing with barcoded antibodies enables multiplexing and doublet detection for single cell genomics. Genome Biol 19, 224 (2018).

22. M. Stoeckius et al., Simultaneous epitope and transcriptome measurement in single cells. Nat Methods 14, 865–868 (2017).

23. A. Butler, P. Hoffman, P. Smibert, E. Papalexi, R. Satija, Integrating single-cell transcriptomic data across different conditions, technologies, and species. Nat Biotechnol 36, 411–420 (2018).

24. L. Velten et al., Human haematopoietic stem cell lineage commitment is a continuous process. Nat Cell Biol 19, 271–281 (2017).

25. D. M. Popescu et al., Decoding human fetal liver haematopoiesis. Nature 574, 365–371 (2019).

26. S. Triana et al., Single-cell proteo-genomic reference maps of the hematopoietic system enable the purification and massive profiling of precisely defined cell states. Nat Immunol 22, 1577–1589 (2021).

27. S. B. Hay, K. Ferchen, K. Chetal, H. L. Grimes, N. Salomonis, The Human Cell Atlas bone marrow single-cell interactive web portal. Exp Hematol 68, 51–61 (2018).

28. B. Psaila et al., Single-Cell Analyses Reveal Megakaryocyte-Biased Hematopoiesis in Myelofibrosis and Identify Mutant Clone-Specific Targets. Mol Cell 78, 477–492.e478 (2020).

29. A. Rodriguez-Meira et al., Unravelling Intratumoral Heterogeneity through High-Sensitivity Single-Cell Mutational Analysis and Parallel RNA Sequencing. Mol Cell 73, 1292–1305.e1298 (2019).

30. D. Van Egeren et al., Reconstructing the Lineage Histories and Differentiation Trajectories of Individual Cancer Cells in Myeloproliferative Neoplasms. Cell Stem Cell 28, 514–523.e519 (2021).

31. T. E. Khoyratty et al., Distinct transcription factor networks control neutrophil-driven inflammation. Nat Immunol 22, 1093–1106 (2021).

32. H. Hirai et al., C/EBPbeta is required for ’emergency’ granulopoiesis. Nat Immunol 7, 732–739 (2006).

33. S. A. Pawar et al., C/EBPδ deficiency sensitizes mice to ionizing radiation-induced hematopoietic and intestinal injury. PLoS One 9, e94967 (2014).

34. V. Bergen, M. Lange, S. Peidli, F. A. Wolf, F. J. Theis, Generalizing RNA velocity to transient cell states through dynamical modeling. Nat Biotechnol 38, 1408–1414 (2020).

35. G. La Manno et al., RNA velocity of single cells. Nature 560, 494–498 (2018).

36. F. A. Wolf et al., PAGA: graph abstraction reconciles clustering with trajectory inference through a topology preserving map of single cells. Genome Biol 20, 59 (2019).

37. R. H. Land et al., The orphan nuclear receptor NR4A1 specifies a distinct subpopulation of quiescent myeloid-biased long-term HSCs. Stem Cells 33, 278–288 (2015).

38. P. R. Freire, O. M. Conneely, NR4A1 and NR4A3 restrict HSC proliferation via reciprocal regulation of C/EBPα and inflammatory signaling. Blood 131, 1081–1093 (2018).

39. F. Notta et al., Distinct routes of lineage development reshape the human blood hierarchy across ontogeny. Science 351, aab2116 (2016).

40. I. García-García et al., Pharmacokinetic and pharmacodynamic comparison of two "pegylated" interferon alpha-2 formulations in healthy male volunteers: a randomized, crossover, double-blind study. BMC Pharmacol 10, 15 (2010).

41. D. Van Egeren et al., Reconstructing the Lineage Histories and Differentiation Trajectories of Individual Cancer Cells in Myeloproliferative Neoplasms. Cell Stem Cell 28, 514–523 e519 (2021).

42. A. S. Nam et al., Single-cell multi-omics of human clonal hematopoiesis reveals that DNMT3A R882 mutations perturb early progenitor states through selective hypomethylation. Nat Genet, (2022).

43. P. van Galen et al., Single-Cell RNA-Seq Reveals AML Hierarchies Relevant to Disease Progression and Immunity. Cell 176, 1265–1281.e1224 (2019).

44. I. Tirosh et al., Dissecting the multicellular ecosystem of metastatic melanoma by single-cell RNA-seq. Science 352, 189–196 (2016).

45. Y. Nie, Y. C. Han, Y. R. Zou, CXCR4 is required for the quiescence of primitive hematopoietic cells. J Exp Med 205, 777–783 (2008).

46. F. Le Naour et al., Upregulation of CD9 expression during TPA treatment of K562 cells. Leukemia 11, 1290–1297 (1997).

47. D. Clay et al., CD9 and megakaryocyte differentiation. Blood 97, 1982–1989 (2001).

48. R. Månsson et al., Molecular evidence for hierarchical transcriptional lineage priming in fetal and adult stem cells and multipotent progenitors. Immunity 26, 407–419 (2007).

49. A. Sanjuan-Pla et al., Platelet-biased stem cells reside at the apex of the haematopoietic stem-cell hierarchy. Nature 502, 232–236 (2013).

50. S. O. Ciurea et al., Pivotal contributions of megakaryocytes to the biology of idiopathic myelofibrosis. Blood 110, 986–993 (2007).

51. H. Chagraoui et al., Prominent role of TGF-beta 1 in thrombopoietin-induced myelofibrosis in mice. Blood 100, 3495–3503 (2002).

52. S. Badalucco et al., Involvement of TGFβ1 in autocrine regulation of proplatelet formation in healthy subjects and patients with primary myelofibrosis. Haematologica 98, 514–517 (2013).

53. A. Ehninger et al., Posttranscriptional regulation of c-Myc expression in adult murine HSCs during homeostasis and interferon-α-induced stress response. Blood 123, 3909–3913 (2014).

54. M. N. Sharif et al., Twist mediates suppression of inflammation by type I IFNs and Axl. J Exp Medicine 203, 1891–1901 (2006).

55. T. Tashi et al., Pegylated interferon Alfa-2a and hydroxyurea in polycythemia vera and essential thrombocythemia: differential cellular and molecular responses. Leukemia 32, 1830–1833 (2018).

56. A. G. X. Zeng et al., Identification of a human hematopoietic stem cell subset that retains memory of inflammatory stress. bioRxiv, 2023.2009.2011.557271 (2023).

57. S. Saeed et al., Epigenetic programming of monocyte-to-macrophage differentiation and trained innate immunity. Science 345, 1251086 (2014).

58. K. U. Belge et al., The proinflammatory CD14+CD16+DR++ monocytes are a major source of TNF. J Immunol 168, 3536–3542 (2002).

59. M. Frankenberger, T. Sternsdorf, H. Pechumer, A. Pforte, H. W. Ziegler-Heitbrock, Differential cytokine expression in human blood monocyte subpopulations: a polymerase chain reaction analysis. Blood 87, 373–377 (1996).

60. E. M. Fast et al., External signals regulate continuous transcriptional states in hematopoietic stem cells. Elife 10, (2021).

61. A. E. Rodriguez-Fraticelli et al., Clonal analysis of lineage fate in native haematopoiesis. Nature 553, 212–216 (2018).

62. E. M. Pietras et al., Chronic interleukin-1 exposure drives haematopoietic stem cells towards precocious myeloid differentiation at the expense of self-renewal. Nat Cell Biol 18, 607–618 (2016).

63. K. A. Matatall, C. C. Shen, G. A. Challen, K. Y. King, Type II interferon promotes differentiation of myeloid-biased hematopoietic stem cells. Stem Cells 32, 3023–3030 (2014).

64. M. Yamashita, E. Passegué, TNF-α Coordinates Hematopoietic Stem Cell Survival and Myeloid Regeneration. Cell Stem Cell, (2019).

65. K. A. Lord, A. Abdollahi, B. Hoffman-Liebermann, D. A. Liebermann, Proto-oncogenes of the fos/jun family of transcription factors are positive regulators of myeloid differentiation. Mol Cell Biol 13, 841–851 (1993).

66. T. I. Lee et al., Transcriptional regulatory networks in Saccharomyces cerevisiae. Science 298, 799–804 (2002).

67. X. Y. Li et al., The role of chromatin accessibility in directing the widespread, overlapping patterns of Drosophila transcription factor binding. Genome Biol 12, R34 (2011).

68. R. Pique-Regi et al., Accurate inference of transcription factor binding from DNA sequence and chromatin accessibility data. Genome Res 21, 447–455 (2011).

69. A. N. Schep, B. Wu, J. D. Buenrostro, W. J. Greenleaf, chromVAR: inferring transcription-factor-associated accessibility from single-cell epigenomic data. Nat Methods 14, 975–978 (2017).

70. J. D. Buenrostro et al., Single-cell chromatin accessibility reveals principles of regulatory variation. Nature 523, 486–490 (2015).

71. J. T. Gaublomme et al., Nuclei multiplexing with barcoded antibodies for single-nucleus genomics. Nat Commun 10, 2907 (2019).

72. S. Ma et al., Chromatin Potential Identified by Shared Single-Cell Profiling of RNA and Chromatin. Cell 183, 1103–1116.e1120 (2020).

73. W. Reith et al., MHC class II regulatory factor RFX has a novel DNA-binding domain and a functionally independent dimerization domain. Genes Dev 4, 1528–1540 (1990).

74. L. Pugliatti et al., The genes for MHC class II regulatory factors RFX1 and RFX2 are located on the short arm of chromosome 19. Genomics 13, 1307–1310 (1992).

75. C. A. Siegrist, B. Mach, Antisense oligonucleotides specific for regulatory factor RFX-1 inhibit inducible but not constitutive expression of all major histocompatibility complex class II genes. Eur J Immunol 23, 2903–2908 (1993).

76. P. Burda, P. Laslo, T. Stopka, The role of PU.1 and GATA-1 transcription factors during normal and leukemogenic hematopoiesis. Leukemia 24, 1249–1257 (2010).

77. M. A. Hall et al., The critical regulator of embryonic hematopoiesis, SCL, is vital in the adult for megakaryopoiesis, erythropoiesis, and lineage choice in CFU-S12. Proc Natl Acad Sci U S A 100, 992–997 (2003).

78. C. L. Semerad, E. M. Mercer, M. A. Inlay, I. L. Weissman, C. Murre, E2A proteins maintain the hematopoietic stem cell pool and promote the maturation of myelolymphoid and myeloerythroid progenitors. Proc Natl Acad Sci U S A 106, 1930–1935 (2009).

79. Y. Zhuang, P. Cheng, H. Weintraub, B-lymphocyte development is regulated by the combined dosage of three basic helix-loop-helix genes, E2A, E2-2, and HEB. Mol Cell Biol 16, 2898–2905 (1996).

80. D. E. Zhang et al., Absence of granulocyte colony-stimulating factor signaling and neutrophil development in CCAAT enhancer binding protein alpha-deficient mice. Proc Natl Acad Sci U S A 94, 569–574 (1997).

81. H. S. Radomska et al., CCAAT/enhancer binding protein alpha is a regulatory switch sufficient for induction of granulocytic development from bipotential myeloid progenitors. Mol Cell Biol 18, 4301–4314 (1998).

82. S. Marecki, C. J. Riendeau, M. D. Liang, M. J. Fenton, PU.1 and multiple IFN regulatory factor proteins synergize to mediate transcriptional activation of the human IL-1 beta gene. J Immunol 166, 6829–6838 (2001).

83. W. Huang, E. Horvath, E. A. Eklund, PU.1, interferon regulatory factor (IRF) 2, and the interferon consensus sequence-binding protein (ICSBP/IRF8) cooperate to activate NF1 transcription in differentiating myeloid cells. J Biol Chem 282, 6629–6643 (2007).

84. M. Hoogenkamp et al., Early chromatin unfolding by RUNX1: a molecular explanation for differential requirements during specification versus maintenance of the hematopoietic gene expression program. Blood 114, 299–309 (2009).

85. C. Yeamans et al., C/EBPalpha binds and activates the PU.1 distal enhancer to induce monocyte lineage commitment. Blood 110, 3136–3142 (2007).

86. C. Le Coz et al., Constrained chromatin accessibility in PU.1-mutated agammaglobulinemia patients. J Exp Med 218, (2021).

87. P. Gutierrez, M. D. Delgado, C. Richard, F. Moreau-Gachelin, J. Leon, Interferon induces up-regulation of Spi-1/PU.1 in human leukemia K562 cells. Biochem Biophys Res Commun 240, 862–868 (1997).

88. H. Stein et al., A new monoclonal antibody (CAL2) detects CALRETICULIN mutations in formalin-fixed and paraffin-embedded bone marrow biopsies. Leukemia 30, 131–135 (2016).

89. S. Cuylen et al., Ki-67 acts as a biological surfactant to disperse mitotic chromosomes. Nature 535, 308–312 (2016).

90. J. S. Jutzi et al., Whole-genome CRISPR screening identifies N-glycosylation as a genetic and therapeutic vulnerability in CALR-mutant MPNs. Blood 140, 1291–1304 (2022).

91. D. Prins et al., The stem/progenitor landscape is reshaped in a mouse model of essential thrombocythemia and causes excess megakaryocyte production. Sci Adv 6, (2020).

92. P. J. Skene, S. Henikoff, An efficient targeted nuclease strategy for high-resolution mapping of DNA binding sites. Elife 6, (2017).

93. S. Elf et al., Mutant Calreticulin Requires Both Its Mutant C-terminus and the Thrombopoietin Receptor for Oncogenic Transformation. Cancer Discov 6, 368–381 (2016).

94. A. L. Brass, E. Kehrli, C. F. Eisenbeis, U. Storb, H. Singh, Pip, a lymphoid-restricted IRF, contains a regulatory domain that is important for autoinhibition and ternary complex formation with the Ets factor PU.1. Genes Dev 10, 2335–2347 (1996).

95. J. M. Pongubala et al., Effect of PU.1 phosphorylation on interaction with NF-EM5 and transcriptional activation. Science 259, 1622–1625 (1993).

96. S. Heinz et al., Simple combinations of lineage-determining transcription factors prime cis-regulatory elements required for macrophage and B cell identities. Mol Cell 38, 576–589 (2010).

97. C. Kopanos et al., VarSome: the human genomic variant search engine. Bioinformatics 35, 1978–1980 (2019).

98. F. Hu et al., ER stress and its regulator X-box-binding protein-1 enhance polyIC-induced innate immune response in dendritic cells. Eur J Immunol 41, 1086–1097 (2011).

99. M. G. Netea, J. Quintin, J. W. van der Meer, Trained immunity: a memory for innate host defense. Cell Host Microbe 9, 355–361 (2011).

100. S. Naik et al., Inflammatory memory sensitizes skin epithelial stem cells to tissue damage. Nature 550, 475–480 (2017).

101. I. Gresser, C. Bourali, Exogenous interferon and inducers of interferon in the treatment Balb-c mice inoculated with RC19 tumour cells. Nature 223, 844–845 (1969).

102. N. Tweezer-Zaks, E. Rabinovich, M. Lidar, A. Livneh, Interferon-alpha as a treatment modality for colchicine-resistant familial Mediterranean fever. J Rheumatol 35, 1362–1365 (2008).

103. S. L. Hauser et al., Ocrelizumab versus Interferon Beta-1a in Relapsing Multiple Sclerosis. N Engl J Med 376, 221–234 (2017).

104. E. Aricò, L. Castiello, I. Capone, L. Gabriele, F. Belardelli, Type I Interferons and Cancer: An Evolving Story Demanding Novel Clinical Applications. Cancers (Basel) 11, (2019).

105. N. G. Sandler et al., Type I interferon responses in rhesus macaques prevent SIV infection and slow disease progression. Nature 511, 601–605 (2014).

106. V. Suppiah et al., IL28B is associated with response to chronic hepatitis C interferon-alpha and ribavirin therapy. Nat Genet 41, 1100–1104 (2009).

107. G. Schreiber, The Role of Type I Interferons in the Pathogenesis and Treatment of COVID-19. Front Immunol 11, 595739 (2020).

108. I. A. Darazam et al., Role of interferon therapy in severe COVID-19: the COVIFERON randomized controlled trial. Sci Rep-uk 11, 8059 (2021).

109. J. J. Kiladjian et al., Long-term outcomes of polycythemia vera patients treated with ropeginterferon Alfa-2b. Leukemia 36, 1408–1411 (2022).

110. L. Huys et al., Type I interferon drives tumor necrosis factor-induced lethal shock. J Exp Med 206, 1873–1882 (2009).

111. J. Hadjadj et al., Impaired type I interferon activity and inflammatory responses in severe COVID-19 patients. Science 369, 718–724 (2020).

112. Q. Zhang et al., Inborn errors of type I IFN immunity in patients with life-threatening COVID-19. Science 370, eabd4570 (2020).

113. M. Ohta, J. S. Greenberger, P. Anklesaria, A. Bassols, J. Massagué, Two forms of transforming growth factor-beta distinguished by multipotential haematopoietic progenitor cells. Nature 329, 539–541 (1987).

114. S. J. Stein, A. S. Baldwin, Deletion of the NF-κB subunit p65/RelA in the hematopoietic compartment leads to defects in hematopoietic stem cell function. Blood 121, 5015–5024 (2013).

115. M. Zhao et al., Megakaryocytes maintain homeostatic quiescence and promote post-injury regeneration of hematopoietic stem cells. Nat Med 20, 1321–1326 (2014).

116. J. M. Scandura, P. Boccuni, J. Massagué, S. D. Nimer, Transforming growth factor beta-induced cell cycle arrest of human hematopoietic cells requires p57KIP2 up-regulation. Proc Natl Acad Sci U S A 101, 15231–15236 (2004).

117. A. Mullally et al., Depletion of Jak2V617F myeloproliferative neoplasm-propagating stem cells by interferon-α in a murine model of polycythemia vera. Blood 121, 3692–3702 (2013).

118. S. Haas et al., Inflammation-Induced Emergency Megakaryopoiesis Driven by Hematopoietic Stem Cell-like Megakaryocyte Progenitors. Cell Stem Cell 17, 422–434 (2015).

119. N. Khan et al., M. tuberculosis Reprograms Hematopoietic Stem Cells to Limit Myelopoiesis and Impair Trained Immunity. Cell 183, 752–770.e722 (2020).

120. Y. Ueda, D. W. Cain, M. Kuraoka, M. Kondo, G. Kelsoe, IL-1R type I-dependent hemopoietic stem cell proliferation is necessary for inflammatory granulopoiesis and reactive neutrophilia. J Immunol 182, 6477–6484 (2009).

121. C. Chen, Y. Liu, P. Zheng, Mammalian target of rapamycin activation underlies HSC defects in autoimmune disease and inflammation in mice. J Clin Invest 120, 4091–4101 (2010).

122. K. Akashi, D. Traver, T. Miyamoto, I. L. Weissman, A clonogenic common myeloid progenitor that gives rise to all myeloid lineages. Nature 404, 193–197 (2000).

123. M. Kondo, I. L. Weissman, K. Akashi, Identification of clonogenic common lymphoid progenitors in mouse bone marrow. Cell 91, 661–672 (1997).

124. J. Adolfsson et al., Identification of Flt3+ lympho-myeloid stem cells lacking erythro-megakaryocytic potential a revised road map for adult blood lineage commitment. Cell 121, 295–306 (2005).

125. J. M. Granja et al., Single-cell multiomic analysis identifies regulatory programs in mixed-phenotype acute leukemia. Nat Biotechnol 37, 1458–1465 (2019).

126. Y. Hao et al., Integrated analysis of multimodal single-cell data. Cell 184, 3573–3587.e3529 (2021).

127. T. Stuart et al., Comprehensive Integration of Single-Cell Data. Cell 177, 1888–1902.e1821 (2019).

128. L. McInnes, J. Healy, J. Melville. (Cornell University, UMAP: Uniform Manifold Approximation and Projection for Dimension Reduction. arXiv preprint *arXiv:1802.03426*, 2018).

129. T. Smith, A. Heger, I. Sudbery, UMI-tools: modeling sequencing errors in Unique Molecular Identifiers to improve quantification accuracy. Genome Res 27, 491–499 (2017).

130. B. M. Bolker et al., Generalized linear mixed models: a practical guide for ecology and evolution. Trends Ecol Evol 24, 127–135 (2009).

131. V. Ntranos, L. Yi, P. Melsted, L. Pachter, A discriminative learning approach to differential expression analysis for single-cell RNA-seq. Nat Methods 16, 163–166 (2019).

132. I. Dolgalev. (msigdbr: MSigDB Gene Sets for Multiple Organisms in a Tidy Data Format. (Manual 2022), https://igordot.github.io/msigdbr/, 2022).

133. G. Korotkevich et al., in Fast gene set enrichment analysis, bioRxiv, Ed. (doi: 10.1101/060012, 2021).

134. A. Liberzon et al., The Molecular Signatures Database (MSigDB) hallmark gene set collection. Cell Syst 1, 417–425 (2015).

135. T. Stuart, A. Srivastava, S. Madad, C. A. Lareau, R. Satija, Single-cell chromatin state analysis with Signac. Nat Methods 18, 1333–1341 (2021).

136. Y. Zhang et al., Model-based analysis of ChIP-Seq (MACS). Genome Biol 9, R137 (2008).

137. M. T. Weirauch et al., Determination and inference of eukaryotic transcription factor sequence specificity. Cell 158, 1431–1443 (2014).

138. A. T. Satpathy et al., Massively parallel single-cell chromatin landscapes of human immune cell development and intratumoral T cell exhaustion. Nat Biotechnol 37, 925–936 (2019).

139. X. Qiu et al., Reversed graph embedding resolves complex single-cell trajectories. Nat Methods 14, 979–982 (2017).

140. J. Cao et al., The single-cell transcriptional landscape of mammalian organogenesis. Nature 566, 496–502 (2019).

141. A. P. Masella et al., BAMQL: a query language for extracting reads from BAM files. BMC Bioinformatics 17, 305 (2016).

142. C. E. Grant, T. L. Bailey, W. S. Noble, FIMO: scanning for occurrences of a given motif. Bioinformatics 27, 1017–1018 (2011).

143. A. V. Persikov et al., A systematic survey of the Cys2His2 zinc finger DNA-binding landscape. Nucleic Acids Res 43, 1965–1984 (2015).

144. J. J. van Dongen et al., EuroFlow antibody panels for standardized n-dimensional flow cytometric immunophenotyping of normal, reactive and malignant leukocytes. Leukemia 26, 1908–1975 (2012).

145. T. Kalina et al., EuroFlow standardization of flow cytometer instrument settings and immunophenotyping protocols. Leukemia 26, 1986–2010 (2012).

146. S. White et al., FlowKit: A Python Toolkit for Integrated Manual and Automated Cytometry Analysis Workflows. Front Immunol 12, 768541 (2021).

147. C. D. Carey et al., Topological analysis reveals a PD-L1-associated microenvironmental niche for Reed-Sternberg cells in Hodgkin lymphoma. Blood 130, 2420–2430 (2017).

148. S. S. Patel et al., Multiparametric in situ imaging of NPM1-mutated acute myeloid leukemia reveals prognostically-relevant features of the marrow microenvironment. Mod Pathol 33, 1380–1388 (2020).

149. J. Rosenthal et al., Building Tools for Machine Learning and Artificial Intelligence in Cancer Research: Best Practices and a Case Study with the PathML Toolkit for Computational Pathology. Mol Cancer Res 20, 202–206 (2022).

150. N. F. Greenwald et al., Whole-cell segmentation of tissue images with human-level performance using large-scale data annotation and deep learning. Nat Biotechnol 40, 555–565 (2022).

151. A. M. Bolger, M. Lohse, B. Usadel, Trimmomatic: a flexible trimmer for Illumina sequence data. Bioinformatics 30, 2114–2120 (2014).

152. B. Langmead, S. L. Salzberg, Fast gapped-read alignment with Bowtie 2. Nat Methods 9, 357–359 (2012).

153. H. Li et al., The Sequence Alignment/Map format and SAMtools. Bioinformatics 25, 2078–2079 (2009).

154. M. Bale. (GitHub).

155. T. L. Bailey, C. E. Grant. (BioRxiv, 2021).

156. F. Ramírez et al., deepTools2: a next generation web server for deep-sequencing data analysis. Nucleic Acids Res 44, W160–165 (2016).

157. M. I. Love, W. Huber, S. Anders, Moderated estimation of fold change and dispersion for RNA-seq data with DESeq2. Genome Biol 15, 550 (2014).

